# Genotyping, sequencing and analysis of 140,000 adults from the Mexico City Prospective Study

**DOI:** 10.1101/2022.06.26.495014

**Authors:** Andrey Ziyatdinov, Jason Torres, Jesús Alegre-Díaz, Joshua Backman, Joelle Mbatchou, Michael Turner, Sheila M. Gaynor, Tyler Joseph, Yuxin Zou, Daren Liu, Rachel Wade, Jeffrey Staples, Razvan Panea, Alex Popov, Xiaodong Bai, Suganthi Balasubramanian, Lukas Habegger, Rouel Lanche, Alex Lopez, Evan Maxwell, Marcus Jones, Humberto García-Ortiz, Raul Ramirez-Reyes, Rogelio Santacruz-Benítez, Abhishek Nag, Katherine R. Smith, Regeneron Genetics Center, Mark Reppell, Sebastian Zöllner, Eric Jorgenson, William Salerno, Slavé Petrovski, John Overton, Jeffrey Reid, Timothy Thornton, Goncalo Abecasis, Jaime Berumen, Lorena Orozco-Orozco, Rory Collins, Aris Baras, Michael R Hill, Jonathan R Emberson, Jonathan Marchini, Pablo Kuri-Morales, Roberto Tapia-Conyer

## Abstract

The Mexico City Prospective Study (MCPS) is a prospective cohort of over 150,000 adults recruited two decades ago from the urban districts of Coyoacán and Iztapalapa in Mexico City. We generated genotype and exome sequencing data for all individuals, and whole genome sequencing for 10,000 selected individuals. We uncovered high levels of relatedness and substantial heterogeneity in ancestry composition across individuals. Most sequenced individuals had admixed Native American, European and African ancestry, with extensive admixture from indigenous groups in Central, Southern and South Eastern Mexico. Native Mexican segments of the genome had lower levels of coding variation, but an excess of homozygous loss of function variants compared with segments of African and European origin. We estimated population specific allele frequencies at 142 million genomic variants, with an effective sample size of 91,856 for Native Mexico at exome variants, all available via a public browser. Using whole genome sequencing, we developed an imputation reference panel which outperforms existing panels at common variants in individuals with high proportions of Central, South and South Eastern Native Mexican ancestry. Our work illustrates the value of genetic studies in populations with diverse ancestry and provides foundational imputation and allele frequency resources for future genetic studies in Mexico and in the United States where the Hispanic/Latino population is predominantly of Mexican descent.

## Introduction

Latin American populations harbor extensive genetic diversity reflecting a complex history of migration throughout the Americas, post-Colonial admixture between continents, and more recent population growth^1–3^. The distinct patterns of genomic variation that exist in these populations have led to key insights into the genetic architecture of rare and common diseases. Founder populations are prevalent throughout Latin America and analyses of deleterious variants that segregate at higher frequency in these groups have identified clinically-relevant novel variants ^4–9^. Moreover, Latin American populations include a significant fraction of Native American indigenous subpopulations that have mostly remained genetically uncharacterized. Admixture between European, Native American and African ancestry groups can result in allele frequency distributions that diverge substantially from ancestral populations. Variants that are rare in one ancestry group but common in another may therefore segregate at a higher frequency in an admixed population, leading to opportunities for novel discoveries in these populations that may be missed when studying single ancestry groups^10, 11^. For example, in one study of Mexican mestizo adults a haplotype in the *SLC16A11* locus that is common in Native Americans but rare in Europeans was strongly associated with type 2 diabetes^12^. In addition to increasing opportunities for variant discovery, genetic analysis of admixed populations can also result in improvements in fine-mapping due to differences in patterns of linkage disequilibrium^10, 13–15^.

Unfortunately, despite the numerous opportunities afforded from studying Latin American populations, Hispanic/Latino individuals from such populations comprise less than 1% of all individuals in genetic population research (despite comprising nearly 10% of the global population). By contrast, European populations comprise over 80% of participants in genomic databases but make up less than 20% of people worldwide. Recent initiatives targeting specific populations^13, 16^ or involving large biobanks (such as the Million Veterans Program^17, 18^ and TOPMed (**URLs**)) have increased the number of Hispanic/Latino individuals included in genetic research, but a sizable gap remains. Additional large genetic studies of Latin American populations are therefore needed to help bridge this gap and enable the implementation of precision medicine in these populations.

Between 1998 and 2004, 159,755 participants aged at least 35 years from two contiguous urban districts of Mexico City (Coyoacán and Iztapalapa) were recruited into the Mexico City Prospective Study (MCPS)^19^. In this study we describe genome-wide array genotyping and whole exome sequencing (WES) on the entire cohort, as well as high-coverage whole genome sequencing (WGS) on a subset of 9,950 participants. We provide a comprehensive genetic profile of the MCPS cohort that reveals complex patterns of relatedness, identity-by-descent (IBD) sharing and runs of homozygosity. By incorporating genotypes from 716 indigenous individuals from 60 of the 68 recognized ethnic groups in Mexico, we apply a range of scalable techniques to finely characterize population structure, continental admixture, and local ancestry in the MCPS cohort.

We also provide a survey of variants according to annotation and frequency, with a particular emphasis on genes that exhibit homozygous loss of function variation. Moreover, we estimate ancestry specific allele frequencies from America, Africa and Europe at 142 million variants, a 10-fold increase over existing resources, made available through a public browser (**URLs**). Lastly, we use the phased WGS dataset as a reference panel to impute genotypes into the full cohort and examine the quality of this imputed dataset compared with the exome sequencing dataset and a TOPMed imputed version of the cohort. The phased WGS dataset will soon be available as a reference panel through the Michigan Imputation Server (**URLs**).

### Overview and comparison of genetic datasets

Of the 159,755 MCPS participants, a blood sample was successfully taken, processed and stored for 155,453 (97.3%). Of these, DNA was successfully extracted for 146,068 (94.0%) and sent for genotyping and exome sequencing. After initial QC procedures (see **Methods**) genotyping array data was available for 138,511 participants and exome data was available for 141,046. (**Supplementary Table 1** provides key baseline characteristics of the 141,046 participants with exome data.) The exomes were sequenced with 98.7% of the samples having 90% of the targeted bases covered at 20X or higher. After applying machine-learning methods to filter out low-quality variants, we identified a total of 9.3 million variants including 4.0 million variants across the coding regions of 19,110 genes. 98.7% of the coding variants were rare (minor allele frequency (MAF) < 1%) (**Table 1, Supplementary Table 2, Methods**) and 1.4 million were unique to MCPS when compared with variants discovered by the UK Biobank (UKB) Exome sequencing study^20^, TOPMed ^16^ and gnomAD ^21^ (**Supplementary Table 3)**. Among the coding variants identified were 1,233,054 (median of 14,900 alleles per individual) synonymous, 2,526,776 (13,585 alleles per individual) missense and 233,650 (354 alleles per individual) putative loss-of-function (pLOF) variants (**Table 1**). The proportion of singletons (30.9%) was much lower than observed in other datasets (e.g., 46.8% in UK Biobank Exomes^20^) due to the way in which households of participants in close neighborhoods were recruited. As expected, the proportion of singletons increased to 36.5% when we restricted to individuals related less than 1^st^ degree, and further to 39.2% when we restricted to individuals related less than 3rd-degree. In addition, we observed more homozygous pLOF variants in MCPS compared with a sample size matched version of the UK Biobank exome dataset (**Supplementary Table 4**).

**Table 1:**
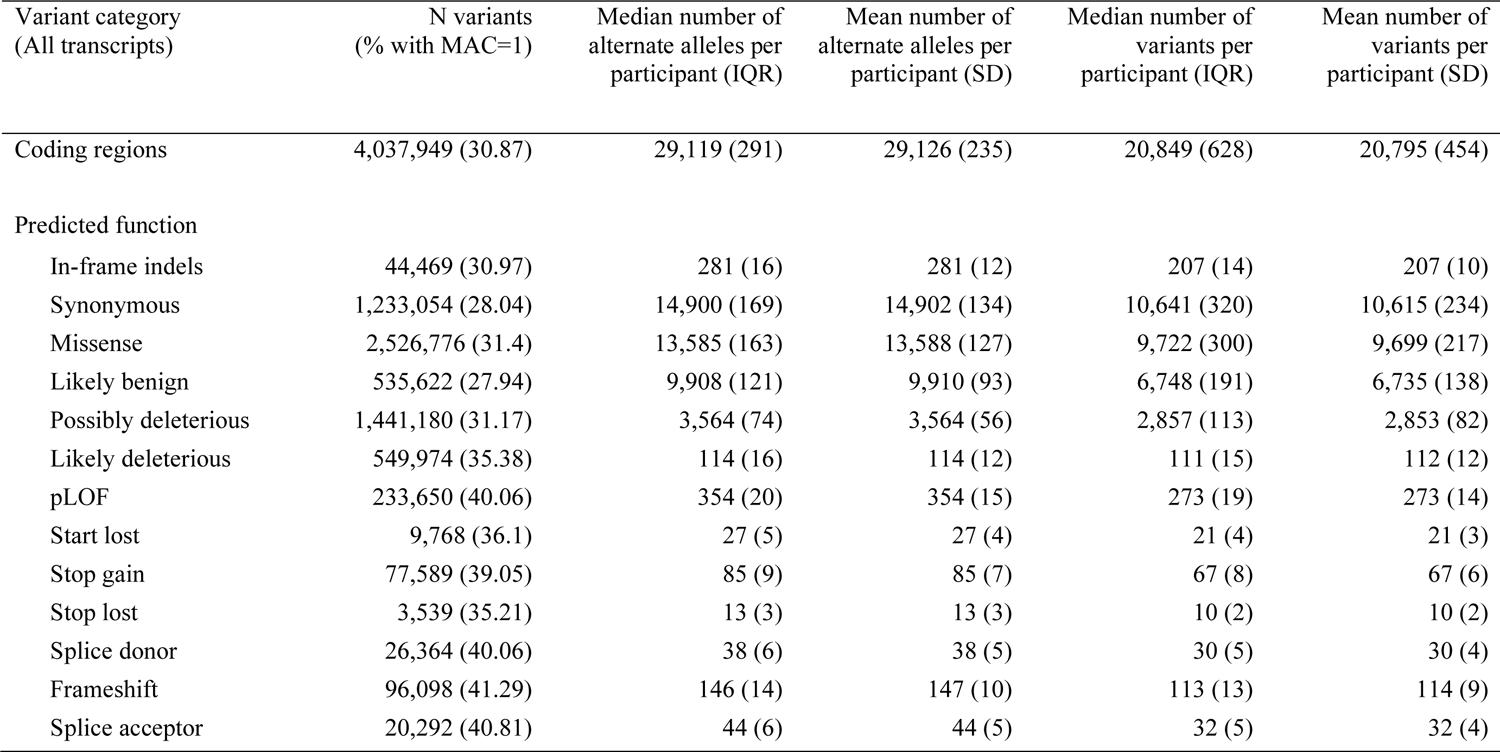
Number of coding variants discovered in exome sequencing of 141,046 MCPS participants. Variants were annotated using VEP. Predicted function for each variant was defined as the most deleterious consequence spanning all protein-coding transcripts in Ensembl v100. MAC = Minor Allele Count, IQR = Inter Quartile Range, SD = Standard Deviation.

A subset of 9,950 MCPS individuals were also whole genome sequenced, with mean depth of 38.5X. After filtering we identified 131.9 million variants in total, of which 1.5 million were coding variants (**Supplementary Table 5-6, Methods**). 96.2% of the variants were rare variants with MAF < 1%. There were 31.5 million unique WGS variants when compared to variants discovered by the TOPMed ^16^ and gnomAD ^21^ WGS datasets (**Supplementary Table 7**).

We compared the WGS and WES in the overlapping set of 9,950 individuals to examine the amount of coding variation called. Both datasets utilized the same calling and annotation framework but used dataset specific machine learning models and hard filters to QC variants. We found that the WGS dataset led to a 2.3% absolute increase in the amount of coding variation when using the canonical gene transcript to annotate variants **(Table 2**), with 93.2%, 4.5% and 2.3% of the union set of sites being called in both datasets, WGS-only and WES-only respectively (**Supplementary Table 8**). When variants were annotated by the most deleterious consequence across all transcripts of a gene, then WGS had 4.6% more coding variants (**Supplementary Table 9**), with 91.1%, 6.6% and 2.3% of the union set of sites being called in both datasets, WGS-only and WES-only respectively (**Supplementary Table 10**). When restricted to exome sequencing capture regions only, the differences between WGS and WES were much smaller (**Supplementary Tables 11-14**). **Supplementary Tables 15-18** compare WGS and WES for variants with alternative allele frequency <1%. The variant sets unique to WGS and WES have similar overlap to TOPMed and gnomAD site lists (**Supplementary Tables 19-22**). Concordance of genotype calls between WGS and WES datasets in 9,950 overlapping samples was very high with a mean biallelic SNP discordance of 0.0064% (**Supplementary Table 23**).

**Table 2:**
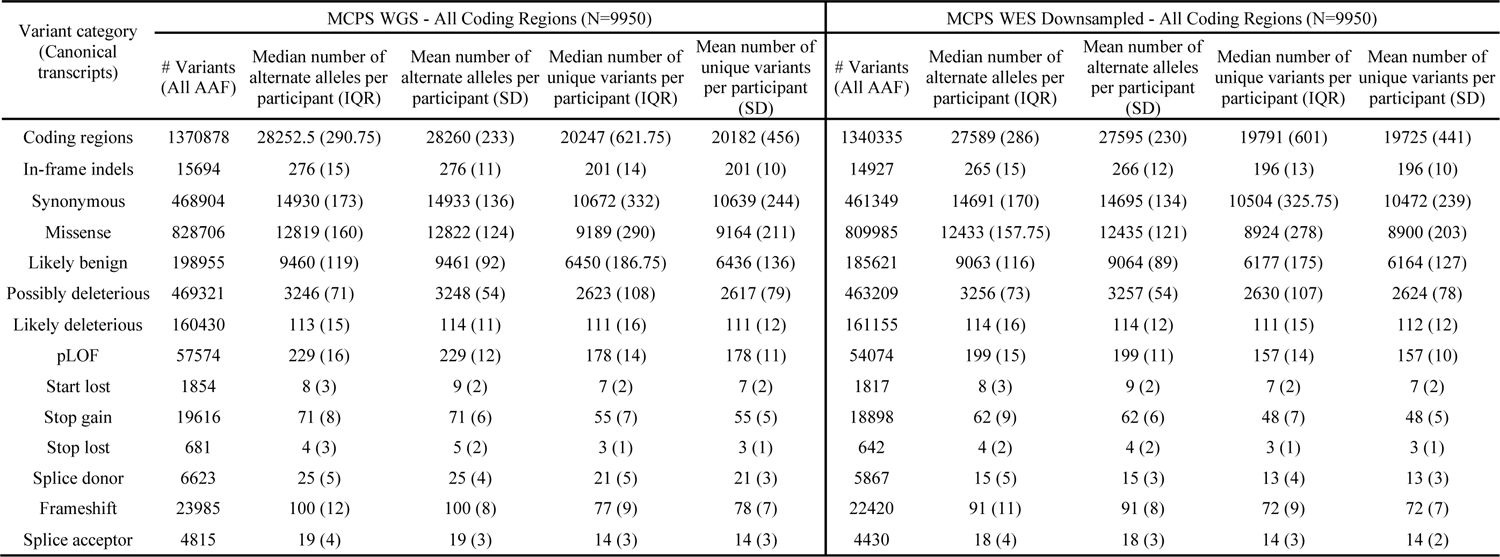
Comparison of WES and WGS datasets in coding genes. Variants were annotated with VEP. Predicted function is defined by canonical transcript consequence in Ensembl v100. Counts are restricted to the same set of 9,950 individuals with both WGS and WGS available. All variants passed QC for the respective platform. AAF = Alternate Allele Frequency, IQR = Inter Quartile Range, SD = Standard Deviation.

| Variant category<br>(Canonical transcripts) | MCPS WGS - All Coding Regions (N=9950) |  |  |  |  | MCPS WES Downsampled - All Coding Regions (N=9950) |  |  |  |  |
| --- | --- | --- | --- | --- | --- | --- | --- | --- | --- | --- |
|  | # Variants<br>(All AAF) | Median number of<br>alternate alleles per<br>participant (IQR) | Mean number of<br>alternate alleles per<br>participant (SD) | Median number of<br>unique variants per<br>participant (IQR) | Mean number of<br>unique variants<br>per participant<br>(SD) | # Variants<br>(All AAF) | Median number of<br>alternate alleles per<br>participant (IQR) | Mean number of<br>alternate alleles<br>per participant<br>(SD) | Median number of<br>unique variants per<br>participant (IQR) | Mean number of<br>unique variants per<br>participant (SD) |
| Coding regions | 1370878 | 28252.5 (290.75) | 28260 (233) | 20247 (621.75) | 20182 (456) | 1340335 | 27589 (286) | 27595 (230) | 19791 (601) | 19725 (441) |
| In-frame indels | 15694 | 276 (15) | 276 (11) | 201 (14) | 201 (10) | 14927 | 265 (15) | 266 (12) | 196 (13) | 196 (10) |
| Synonymous | 468904 | 14930 (173) | 14933 (136) | 10672 (332) | 10639 (244) | 461349 | 14691 (170) | 14695 (134) | 10504 (325.75) | 10472 (239) |
| Missense | 828706 | 12819 (160) | 12822 (124) | 9189 (290) | 9164 (211) | 809985 | 12433 (157.75) | 12435 (121) | 8924 (278) | 8900 (203) |
| Likely benign | 198955 | 9460 (119) | 9461 (92) | 6450 (186.75) | 6436 (136) | 185621 | 9063 (116) | 9064 (89) | 6177 (175) | 6164 (127) |
| Possibly deleterious | 469321 | 3246 (71) | 3248 (54) | 2623 (108) | 2617 (79) | 463209 | 3256 (73) | 3257 (54) | 2630 (107) | 2624 (78) |
| Likely deleterious | 160430 | 113 (15) | 114 (11) | 111 (16) | 111 (12) | 161155 | 114 (16) | 114 (12) | 111 (15) | 112 (12) |
| pLOF | 57574 | 229 (16) | 229 (12) | 178 (14) | 178 (11) | 54074 | 199 (15) | 199 (11) | 157 (14) | 157 (10) |
| Start lost | 1854 | 8 (3) | 9 (2) | 7 (2) | 7 (2) | 1817 | 8 (3) | 9 (2) | 7 (2) | 7 (2) |
| Stop gain | 19616 | 71 (8) | 71 (6) | 55 (7) | 55 (5) | 18898 | 62 (9) | 62 (6) | 48 (7) | 48 (5) |
| Stop lost | 681 | 4 (3) | 5 (2) | 3 (1) | 3 (1) | 642 | 4 (2) | 4 (2) | 3 (1) | 3 (1) |
| Splice donor | 6623 | 25 (5) | 25 (4) | 21 (5) | 21 (3) | 5867 | 15 (5) | 15 (3) | 13 (4) | 13 (3) |
| Frameshift | 23985 | 100 (12) | 100 (8) | 77 (9) | 78 (7) | 22420 | 91 (11) | 91 (8) | 72 (9) | 72 (7) |
| Splice acceptor | 4815 | 19 (4) | 19 (3) | 14 (3) | 14 (3) | 4430 | 18 (4) | 18 (3) | 14 (3) | 14 (2) |

A total of 138,511 MCPS individuals were genotyped on the Illumina GSA v2 beadchip and passed quality control (**Methods, Supplementary Table 24**). Array genotypes were highly concordant with WGS and WES genotypes in overlapping samples (mean biallelic SNP discordance of 0.03% for both datasets) (**Supplementary Table 23**).

### Relatedness

The genetic data allowed us to investigate familial relatedness within the cohort which was expected to be high due to the household recruitment strategy. Accounting for relatedness is essential for validity of GWAS^22^ and epidemiological studies^23^ and can be leveraged in heritability estimation to reduce bias of shared environmental effects^24^. We characterized familial relatedness using the quality control filtered genotyping array dataset (**Methods**). We used shared identical-by-descent (IBD) segments to infer relatedness to avoid estimation biases in samples from admixed populations that can occur when using methods based on population allele frequency estimates ^25^. We applied KING software^26^ to unphased data, and the hap-IBD^27^ and IBDkin^28^ methods to a phased array dataset (**Methods**). Both unphased and phased approaches produced comparable results (**Supplementary Figure 1**).

**Figure 1a** and **Supplementary Figure 2** illustrate the extensive relatedness identified in MCPS. There are 31,597 parent-offspring, 29,482 full sibling, 47,080 second-degree relative, and 120,180 third-degree relative pairs. A small proportion (0.05%) of parent-offspring pairs had genotypes at a small number of loci that were inconsistent with this type of relationship, resulting in elevated estimates of sharing 0 alleles IBD. We determined genotyping error to be the most likely cause of this phenomenon as opposed to uniparental disomy. Close to 71% (97,953 individuals) in MCPS have at least one relative in the study that is third-degree or closer and many of the MCPS participants have multiple close relatives (**Figure 1b)**. The largest connected component in a graph of individuals with third-degree relationships or closer involves 22% of the cohort (30,682 individuals) (**Supplementary Figure 3**). These levels of relatedness are much higher than those observed in the UK Biobank^1^, but are comparable to the Geisinger Health Study^29^ (both MCPS and the Geisinger Health Study recruited participants from regions with families living in close proximity) **(Supplementary Table 25)**. We used PRIMUS^30^ to reconstruct 22,766 first-degree family networks containing a total of 65,777 individuals with a median size of 2.9, up to a maximum size of 48 people, including 3,595 nuclear families (**Supplementary Figure 4, Supplementary Table 26**). A graph of 14,428 individuals with second-degree family networks of size greater than four highlights the complexity of the patterns of relatedness, as well as partial clustering of relationships within districts of Coyoacán and Iztapalapa (**Supplementary Figure 5**). The largest connected component in this graph contains 9,180 individuals. We also investigated relationships within and across the two districts (see **Supplementary Table 27**). With reconstruction of pedigree networks in MCPS, we were able to investigate the proportion of relatives who cross boundaries and have residences in different districts. Among the first-degree relatives, we find that only 3% of parent-child pairs and 7% of full sibling pairs lived in different districts. The percentages of second- and third-degree relative pairs with residences in different districts was 13% and 17%, respectively, which is much lower than would be expected if there was random mixing of individuals from the contiguous districts. Interestingly, although there was a marked 10% to 15% decrease in the percentages of second- or third-degree relative pairs who both had a residence in the Coyoacán district compared with first-degree relationship types, the percentages of relative pairs who had a residence in the Iztapalapa district remained fairly consistent across relationship types (**Supplementary Table 27**). These results provide some insight into patterns of migration (or lack thereof) within families between the Coyoacán and Iztapalapa districts.

**Figure 1:**
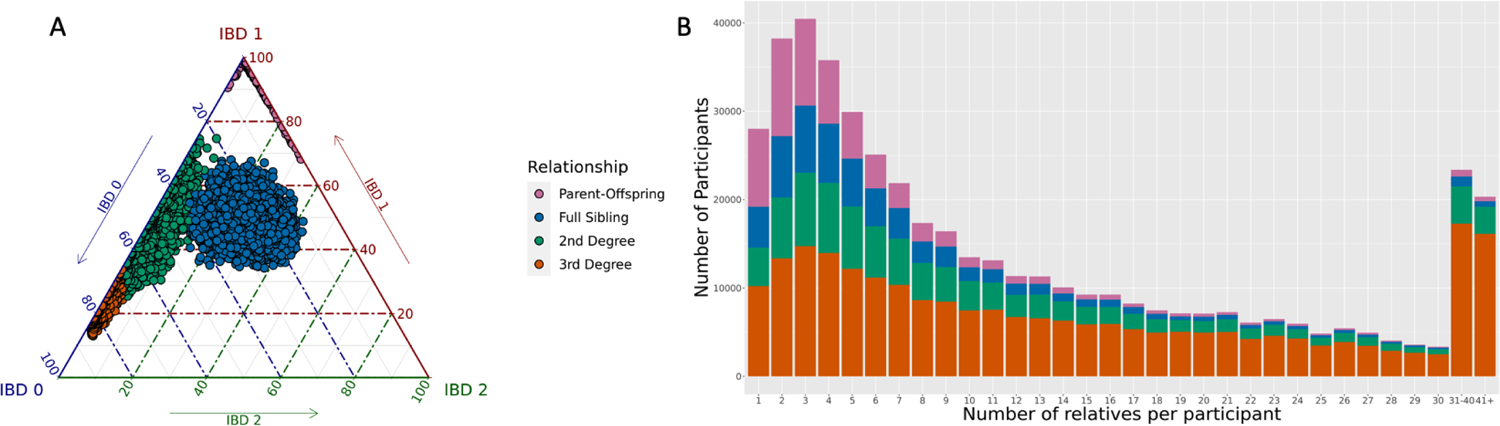
Familial relatedness. (a) Percentage of the genome estimated to have 0, 1 or 2 alleles identical-by-descent (IBD) (b) Distribution of the number of relatives that participants have in the MCPS cohort. The height of each bar shows the count of participants with the stated number of relatives. The colors indicate the proportions of each relatedness class within each bar.

### Population structure

The genetic dataset allowed us to characterize the ancestry composition and heterogeneity of MCPS individuals relative to pre-Columbian population structure in Mexico. Accounting for genetic ancestry and admixture is crucial in GWAS^31^ and can be used to boost power^32^ and for explorations of polygenic risk scores portability^33^. We used a variety of complementary analysis approaches to investigate the fine-scale population structure in the MCPS dataset, with a specific emphasis on elucidating the Native American component of genetic ancestry. Firstly, we applied PCA to a reference dataset of 108 African (Yoruba) and 107 European (Iberian) samples from the 1000 Genomes (1000G) dataset^34^, and 591 unrelated samples from 60 Native Mexican groups corresponding to Central, Southern, South Eastern, Northern and North Western regions of Mexico from the Metabolic Analysis of an Indigenous Sample (MAIS)^2^ (see **Methods, Figure 2, Supplementary Figure 6**). We included a representative set of unrelated MCPS samples (n=500) in the PCA model fitting procedure and projected the remaining 138,011 MCPS samples onto the inferred PC axes. **Figure 2a** shows that PC1 and PC2 separate Native Mexican, African and European samples, and that MCPS samples lie on the axis between Native Mexican and European samples. **Figure 2b** shows that PC3 differentiates Native Mexican geographic sub-groups and suggests that the majority of MCPS samples have ancestry from Central, Southern and South Eastern Mexico.

**Figure 2:**
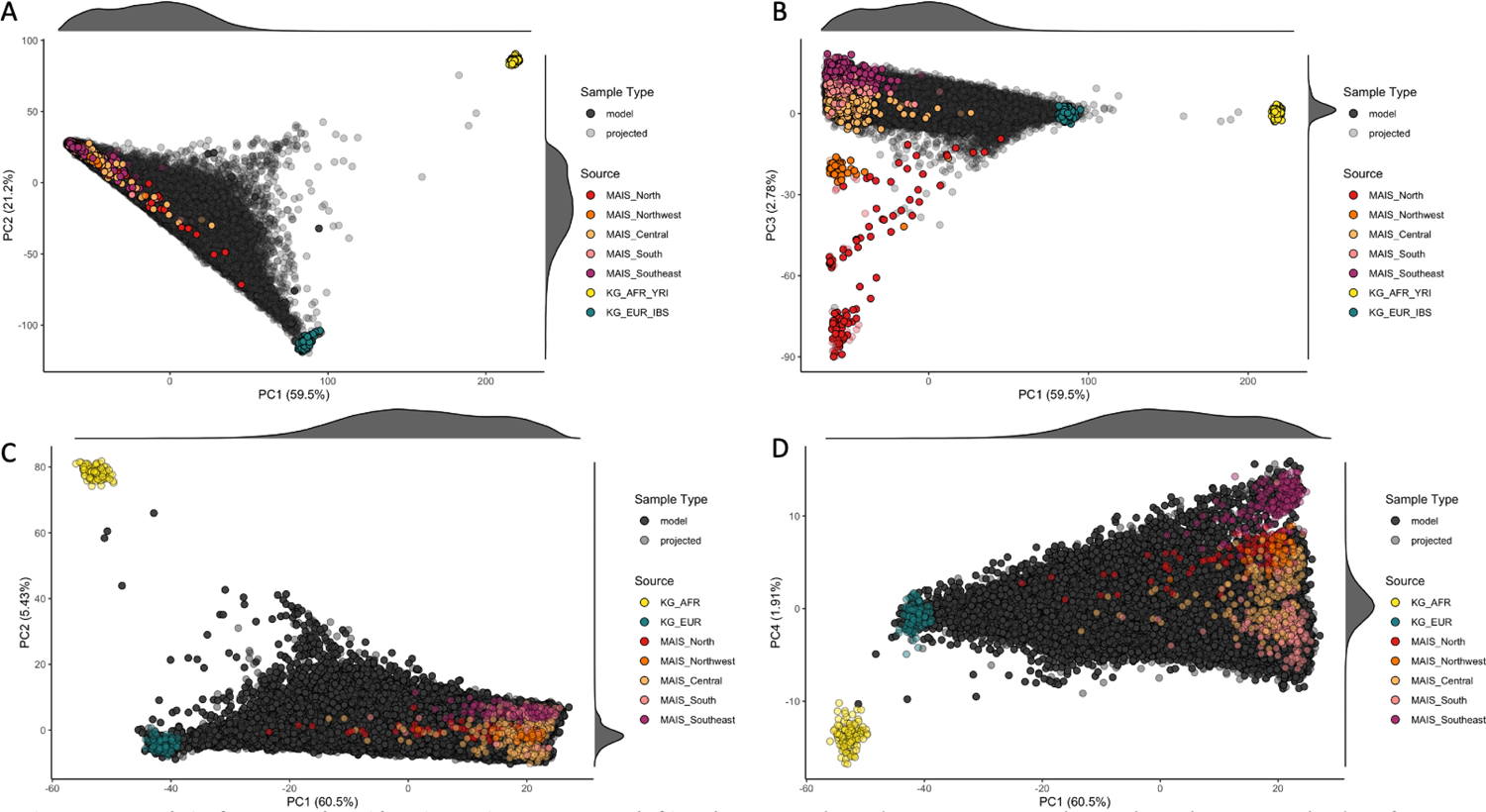
PCA analysis of MCPS together with Native Mexican, European and African datasets. Panels A and B use 500 MCPS samples together with 108 African Yoruba (KG_AFR_YRI) and 107 European Iberian (KG_EUR_IBS) samples from the 1000 Genomes Project dataset, and 591 unrelated samples from 60 Native Mexican groups corresponding to Central, Southern, South Eastern, Northern and North Western regions of Mexico from the Metabolic Analysis of an Indigenous Sample (MAIS). Panels C and D use an unrelated set of 58,051 samples together with the 1000 Genomes and MAIS samples. All other MCPS samples are projected onto the axes.

To provide more focus on the genetic variation within the MCPS dataset we also applied PCA to a filtered array dataset of 58,051 unrelated MCPS samples, with all other MCPS samples and 1000G, HGDP and MAIS samples projected onto the inferred PC axes (**Figure 2c,d, Supplementary Figure 7**). This analysis further highlighted that Mesoamerican ancestry from indigenous groups in Central, Southern and South Eastern Mexico predominates, whereas ancestry from indigenous groups in the Northern and more arid regions of the country is sparsely represented in MCPS.

Examination of the SNP loadings from this PCA analysis highlighted that many PCs exhibited local effects attributable to long-range LD consistent with recent admixture. More stringent LD filtering reduces this phenomenon and suggests that analysis of large scale admixed datasets requires careful selection of PCs used in GWAS (**Supplementary Figures 8-10**). Parametric admixture estimation also corroborated significant ancestry proportions from Mesoamerican ancestry groups among MCPS participants (**Supplementary Figure 11, Methods**).

While PCA aims to uncover population structure in a dataset using a set of mostly unlinked markers, haplotype-based approaches that can utilize linkage disequilibrium (LD) between SNPs have been shown to uncover much finer scale population structure ^35, 36^. We applied two different methods to measure the similarity between pairs of individuals using phased array haplotypes from a set of unrelated MCPS individuals. The first approach used identical-by-descent (IBD) segments^27^, and the second approach measured the extent of haplotype sharing using a scalable implementation of a haplotype-copying hidden Markov model^37^ (**Methods**). Both of these approaches produced low-dimensional representations with noticeably more ‘star-like’ structure than PCA (**Supplementary Figures 12-13**). In combination with ancestry proportions from the local ancestry inference (see next section), this highlighted the ability of these approaches to delineate the contributions of Mesoamerican and European ancestry more clearly.

### Local ancestry estimation

We carried out a supervised population structure analysis by applying local ancestry inference (LAI) with RFMix^38^ using a reference panel of haplotypes from Africa, Europe and America (**Methods**). **Supplementary Figure 14** shows local ancestry at segments genome-wide for 12 representative MCPS individuals estimated from the LAI results and **Figure 3** shows population distributions of LAI-based ancestry proportion estimates, including five indigenous sub-groups within Mexico. Overall, we estimate that 66.0% of autosomal ancestry was attributable to Native Mexican groups with the majority coming from Central Mexico (35.6%). Southern Mexico and South Eastern Mexico accounted for 15.9% and 11.8% respectively, with much smaller amounts of ancestry attributable to Northern Mexico (1.6%) and North Western Mexico (1.1%). In addition, 2.9% and 31.1% of ancestry was attributable to African and European groups respectively. We observed that MCPS individuals with the most Native Mexican ancestry seem to have a greater relative contribution from indigenous groups from Southern Mexico (i.e. from the states of Oaxaca and Veracruz) (**Supplementary Figure 15)**. We also find lower amounts of Native Mexican ancestry and higher amounts of European ancestry in Coyoacán than in Iztapalapa, consistent with socio-demographic characteristics of these districts (**Supplementary Text**).

**Figure 3:**
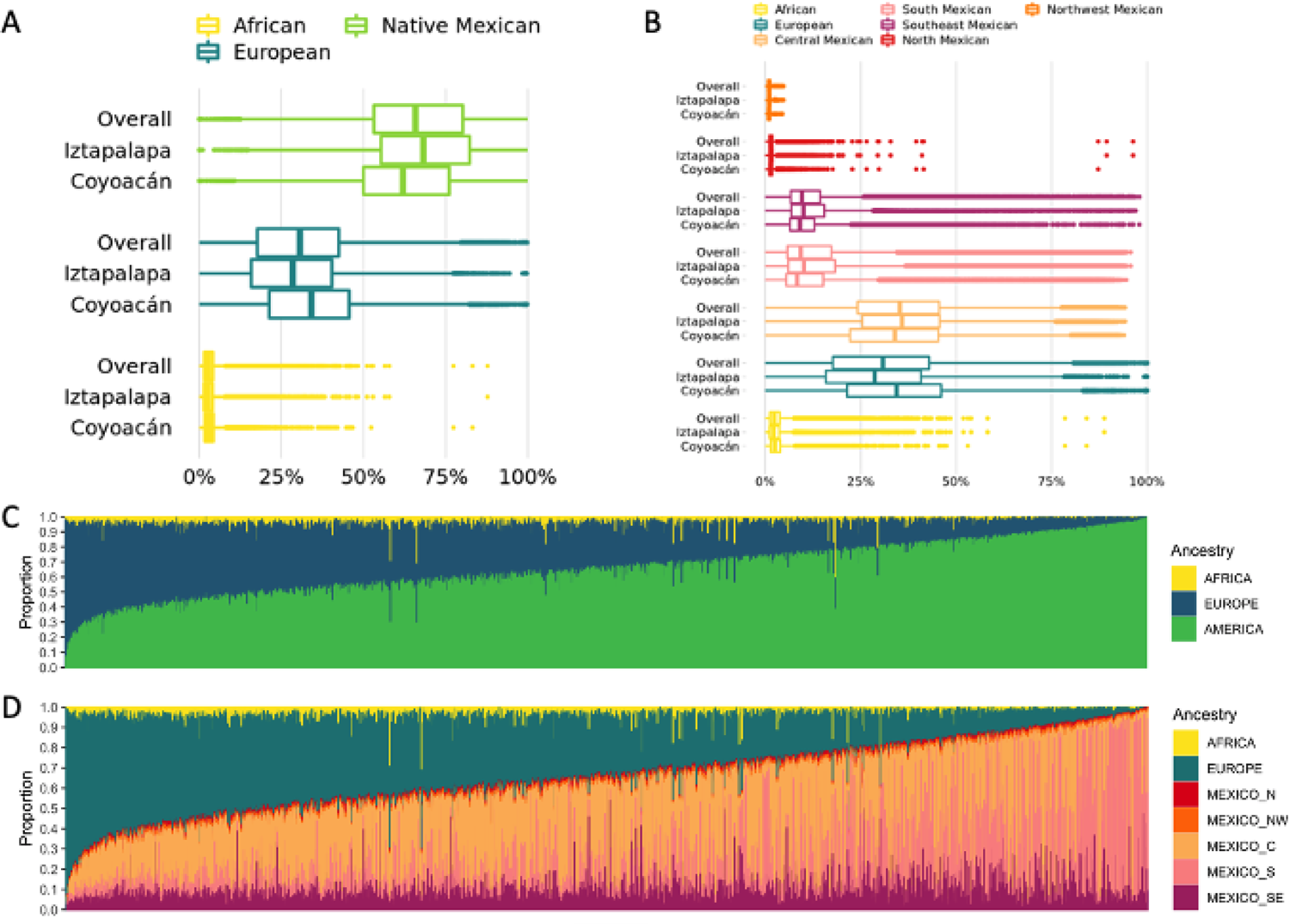
Global ancestry proportions estimated from local ancestry inference (LAI). Distributions of LAI-based global ancestry proportions from a 7-way analysis (panel B) and reduced to 3 continental groups (panel A). Stacked bar plots of 3-way (panel C) and 7-way (panel D) local ancestry proportions for 138,511 MCPS individuals.

**Figure 4:**
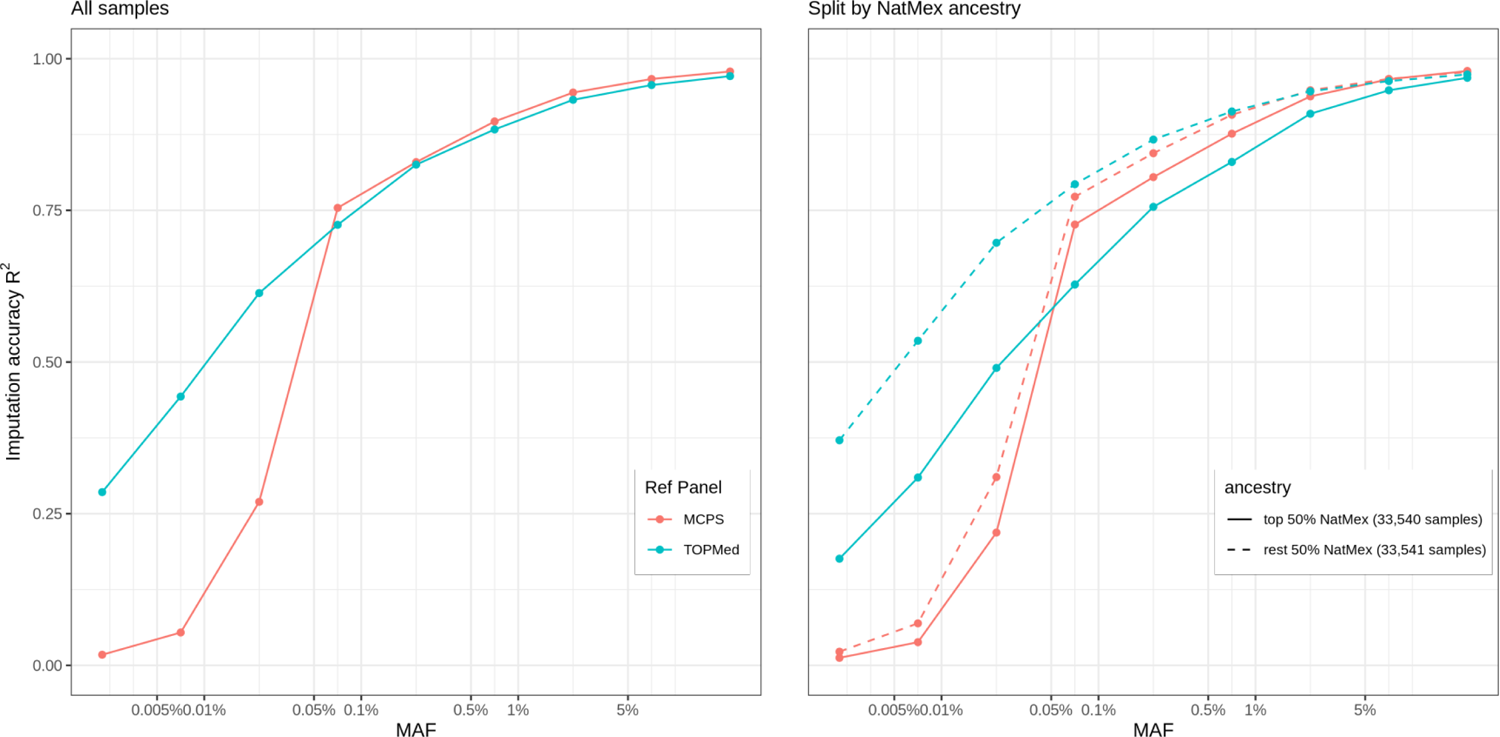
Imputation accuracy using the MCPS10k and TOPMed imputation panels. Accuracy is measured using the R^2^ between the imputed variants and 125,639 variants measured using exome sequencing on chromosome 2 in 67,079 MCPS samples not in (or related to) the MCPS reference panel samples. Results are stratified by allele frequency (x-axis on log10 scale), reference panel (red = MCPS, blue = TOPMed) and into two groups (top and bottom 50% of Native Mexican ancestry shown by solid and dashed lines). The left-hand plot shows results at on all samples. The right-hand plot shows the results stratified by the amount of Native Mexican estimated in each sample.

**Figure 5:**
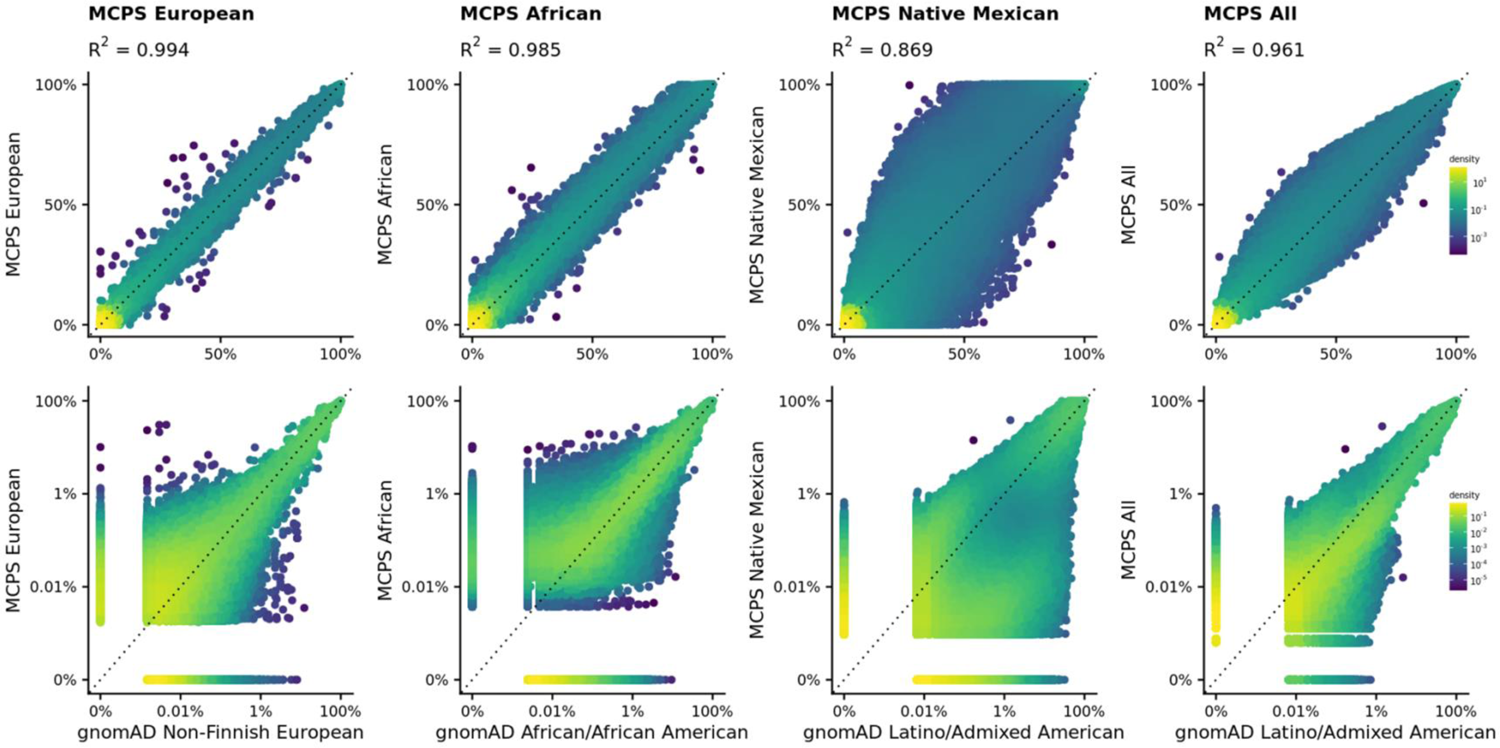
Allele frequency comparison between MCPS WES and gnomAD. Allele frequencies on linear (top) and log (bottom) scale. The comparisons from left to right are MCPS European vs gnomAD Non-Finnish European, MCPS African vs gnomAD African, MCPS Native American vs gnomAD Latino/Admixed American and All MCPS vs gnomAD Latino/Admixed American.

Using 3,595 parent couples inferred from the genetic relatedness analysis we observed significant correlation in ancestry between partner pairs (**Supplementary Figure 16**) as has been observed in other admixed studies^39–41^. We used a linear model to predict ancestry of each partner using the ancestry of their spouse, education level (4 categories) and district (Coyoacán and Iztapalapa) of both partners. We found that education and district explained between 0.5-5% of the variation in ancestry, whereas spousal ancestry explained between 15-26% of the variation in ancestry. This suggests that genomic ancestry is a much better predictor of partners’ ancestry than these sociodemographic factors.

**Supplementary Figure 17** shows the proportion of ancestry across each chromosome from a 3-way LAI analysis (**Methods**). This highlighted an excess of African ancestry in and around the MHC on chromosome 6 (African 17.3%, P-value = 2.9e-14; **Supplementary Figure 18)** consistent with previous observations^42^. We also observed ancestry proportions on chromosome X that exhibited elevated levels of Native Mexican ancestry compared to autosomes (African 3.2%, Native Mexican 73.8%, European 22.7%), consistent with an imbalance of male and female contributions to admixture. Using a simplified population mixture event model^43, 44^ that best fits the observed chromosome X ancestry proportions we estimate that the proportion of Native Mexican ancestry explained by female contribution was 71.3%, while for Europeans the female contribution accounted for 7.5%. (**Supplementary Table 28**).

### Homozygosity

The relatedness analysis highlighted a subset of parent-offspring pairs with elevated levels of sharing two alleles IBD (**Figure 1a**), which can be caused by extensive homozygosity within a population. The exome variant survey highlighted an increased amount of homozygous pLoF variants compared to the UK Biobank exome dataset (**Supplementary Table 4**). Homozygosity, particularly at pLoF variants, can be especially useful in understanding gene function, drug discovery, and for call back studies^45^. Population bottlenecks and consanguinity can increase homozygosity, whereas admixture can decrease homozygosity within a population. We estimated levels of homozygosity in each MCPS individual by estimating runs of homozygosity (ROH) from the phased array dataset using hap-IBD^27^ (**Methods**). There were 60,722 MCPS participants (43.9%) who had at least one ROH segment of length 4 centimorgans (cM) or longer. The mean homozygosity across the whole dataset was 0.34%, and 0.78% among the 60,722 individuals with at least one ROH segment greater than or equal to 4 cM (**Supplementary Tables 29, Supplementary Figure 19**). As a comparison, we ran the same analysis on the UK Biobank phased array genotypes and found the mean homozygosity was 0.07%, and 0.59% in the 55,206 (11.3%) of the participants with at least one ROH segment.

We observed that the total amount and number of ROH segments was positively correlated with the proportion of ancestry that is native to Mexico (**Supplementary Figure 20**). Combining ROH segments with local ancestry estimates (**Methods**) we found that 79.0% of ROH segments can be assigned to Native Mexican ancestry, clearly exceeding the 66.3% average amount of Native Mexican ancestry in the sample. Similarly, we observed a depleted proportion of ROH in European and African segments (19.10% and 1.9% respectively) compared to the average amount of European and African ancestry in the sample (30.2% and 3.5% respectively). We observed that 68.4% of ROH segments are homozygous for a single ancestry across their whole length. These results are consistent with previous reports that Native American populations tend to have higher levels of homozygosity than European and African populations ^45^.

The mean number of rare homozygous pLOFs (rhLOF), and the proportion of rhLOFs in ROH was correlated with the proportion of the genome in ROH segments (**Supplementary Figure 21**). Overall, for LOF variants with allele frequency <0.1% we identified 3,763 rhLOF genotypes at 2,646 variants in 2,169 different protein-coding genes in 3,519 individuals, and 52.2% of these were found within ROH segments (recall that, overall, <0.5% of these genomes lies in ROH segments). Given the rate of rhLOF variants in MCPS (**Supplementary Table 4**), we investigated the local ancestry assignment for each observed rhLOF within ROH and observed that segments of Native Mexican ancestry account for 62.6% of rhLOFs (**Supplementary Table 30)**.

### An MCPS imputation reference panel

We created a phased haplotype reference panel (MCPS10k) for the purposes of genotype imputation that is being made available via the Michigan Imputation Server (see **URLs**). The phasing process utilized phase information from sequencing reads and pedigrees, and WGS variants were phased onto an array haplotype scaffold to facilitate ancestry specific allele frequency estimation (**Methods**). Using the WGS trios we estimate that haplotypes were phased with a switch error rate of 0.0024 (**Methods, Supplementary Figure 22**) and we observed that switch error rate depended upon ancestry proportion (**Supplementary Figure 23**).

We assessed the utility of the MCPS10k reference panel for genotype imputation by imputing chromosome 2 using the phased array dataset of 67,079 MCPS individuals not included in the reference panel and pruned for relationships up to the first degree. For comparison, we also imputed the MCPS dataset using the diverse TOPMed reference panel that includes 47,159 European, 24,267 African, and 17,085 admixed American genomes (**Methods**).

MCPS10k and TOPMed imputation produced, respectively, a set of 9,801,290 and 9,437,266 autosomal variants with imputation info score >0.3. However, the information scores (a well calibrated measure of accuracy) for an overlapping set of 6,473,872 variants were generally higher using MCPS10k than TOPMed for MAF bins greater than 0.01% (**Supplementary Figure 24**).

We compared the MCPS10k and TOPMed imputed genotypes to the exome sequencing data at 128,745 sites on chromosome 2. **Figure 4** shows the results of the imputation accuracy stratified by allele frequency, reference panel and degree of Native Mexican ancestry (defined as two groupings with individuals split above and below the median proportion of Native Mexican ancestry). The results show that MCPS10k outperformed TOPMed for MAF > 0.1%, but below that threshold the TOPMed panel had better performance. Furthermore, we find that the MCPS10k panel provided the greatest imputation benefits for those samples with the highest proportions of Native Mexican ancestry.

Finally, we assessed the imputation performance in 1000 Genomes individuals with Mexican ancestry from Los Angeles (MXL) and found that TOPMed provided improved imputation performance compared to MCPS10k (**Supplementary Figure 25-26**). Our ADMIXTURE analysis of the MXL samples suggests that they have substantially higher European ancestry than MCPS samples (median 44% versus 28%). In addition, the MXL samples have less ancestry from Central, South and South East Mexico, and more from North and North West Mexico than MCPS (**Supplementary Figure 27)**. Previous studies^46^ have suggested that Mexican-Americans from California tend to have increased Native American ancestry from Northwest Mexico as compared to individuals from Mexico City. The limited ancestry from North and North Western Mexico in MCPS and the large number of European reference samples in TOPMed likely explains why the MCPS10k panel does not provide the best imputation accuracy in the MXL samples. Similarly, the TOPMed panel provided the best performance in 1000 Genomes individuals with Peruvian ancestry from Lima (PEL), Colombian ancestry from Medellin (CLM) and Puerto Rican ancestry from Puerto Rico (PUR) compared to MCPS10k (**Supplementary Figure 25-26**). These results emphasize the value of closely matching the ancestry of imputation reference panels to the samples being studied. While our panel provides improved imputation for individuals of Mesoamerican Mexican ancestry, additional panels may be required to provide similar benefits for other Latin American populations with admixture from different Native American ancestral populations.

### Ancestry specific allele frequency estimation

We combined the LAI results with the phased WES and WGS datasets to estimate Native Mexican, African and European allele frequencies at 141,802,412 genetic variants, increasing by 10-fold the number of LAI-resolved frequencies currently available in the gnomAD browser (see schematic in **Supplementary Figure 28**). These frequencies are available in a public browser (see **URLs**). Median sample sizes for estimation of Native Mexican, African and European ancestry were 91,856, 4,312 and 42,009 respectively for WES variants, and 6,549, 341 and 3,058 for WGS variants. For comparison, gnomAD v3.1 median sample sizes are 7,639, 20,719 and 34,014 for Latino/Admixed American, African and Non-Finnish European ancestries. **Figure 5** compares WES allele frequency estimates using our deconvolution approach in MCPS to the more direct approach used in gnomAD v3.1. European allele frequencies showed excellent agreement (*r^2^* = 0.994) and African allele frequencies only showed slightly less agreement (*r^2^* = 0.987), despite greater heterogeneity in African ancestry populations and the lower median African sample size in the MCPS cohort. **Supplementary Figure 29** compares MCPS WGS and gnomAD allele frequencies.

**Table 3** shows the allele frequencies at 46 loci previously reported to show trait associations in contemporary Mexican or other Latin American populations. For example, we found that the top SNP associated with type 2 diabetes at the *SLC16A11* locus^12^ - rs75493593 - has an overall frequency of 36% but population-specific allele frequencies of 0.1%, 0.7% and 53% in African, European and Native Mexican populations, respectively. This is in agreement with previous estimates reported by the SIGMA Type 2 Diabetes Consortium. Another notable example occurs at the *IGF2* locus where the pLOF splice acceptor variant rs149483638 that confers protection against type 2 diabetes^47^ and has an overall frequency of 23% but population-specific allele frequencies of 0.06%, 0.05% and 35% in African, European and Native Mexican populations, respectively. Moreover, the rare *MC4R* missense variant rs79783591 associated with obesity^48^ is absent from the gnomAD browser but has an overall frequency of 1.1% in MCPS with an inferred Native Mexican frequency of 1.6%, and African and European frequencies less than 0.05%.

**Table 3:**
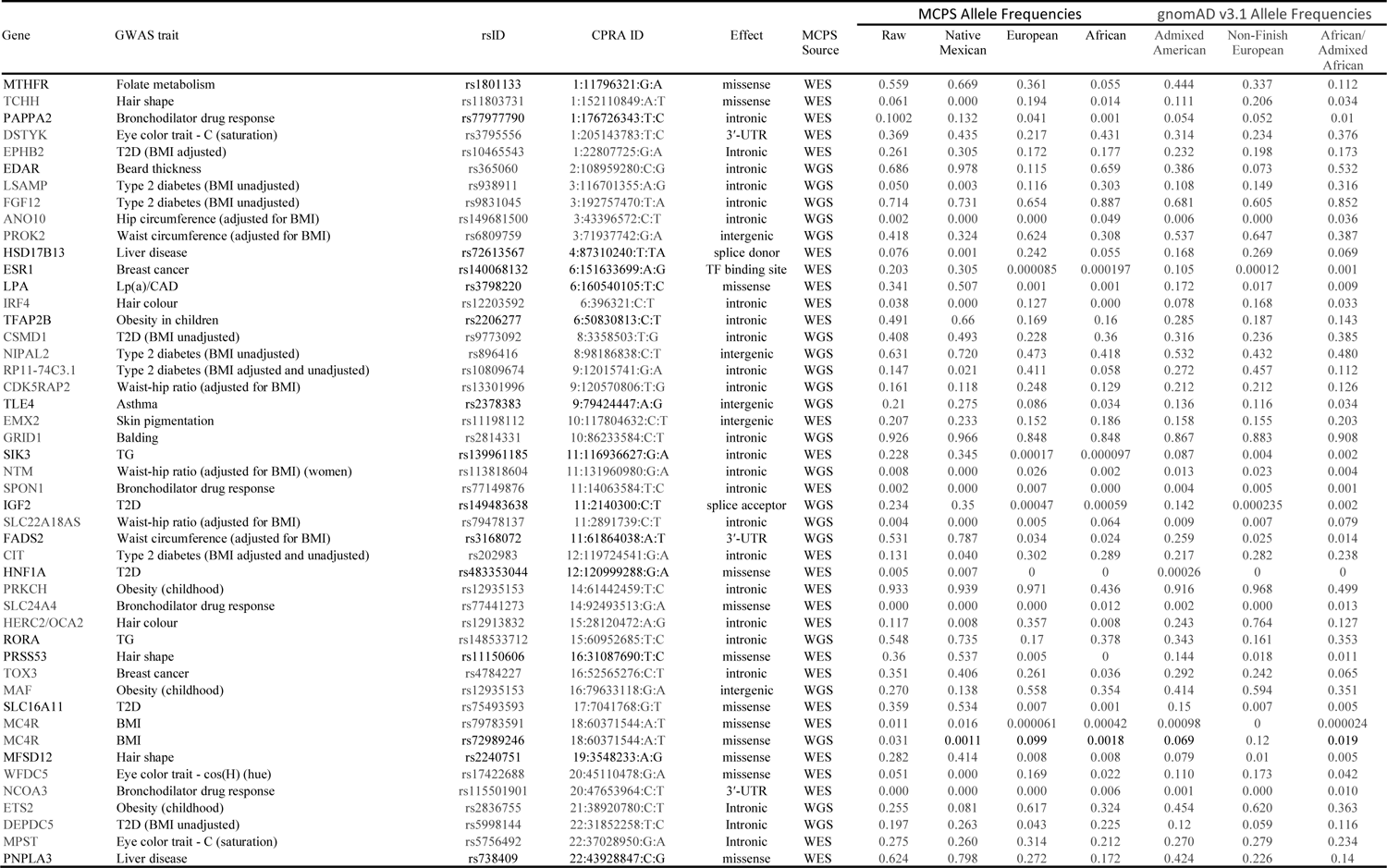
Ancestry specific allele frequencies at GWAS loci previously reported in studies of Mexican and Latin/Central American populations. MCPS Native Mexican, European and African allele frequencies, estimated in MCPS WES/WGS data using our deconvolution approach, are reported together MCPS Raw allele frequencies calculated directly on raw MCPS data. Allele frequencies for three relevant population groups available in gnomAD 3.1 are added for comparison.

We used the 3-way LAI segments to further decompose the annotated variants into three continental groups and found that across all variant classes the highest levels of variation were found in African segments and lower levels in Native Mexican and European segments, consistent with the demographic history of these populations (**Supplementary Table 31**). For example, we estimate that the mean number of pLOF variants in Native Mexican, European and African genomes to be 347, 361 and 427 respectively, although rare homozygous pLOF were more frequent among longer ROHs of Native American ancestry as shown above.

## Discussion

The MCPS genetic data resources described in this study represent the largest in Mexico to date, the most extensive sequencing study in individuals of non-European ancestry, and a major contribution towards the goal of increasing the diversity of genetic collections. Through scalable genotype and haplotype-based approaches to characterize fine-scale population structure and admixture, we traced the Native American component of ancestry within MCPS individuals to predominantly Mesoamerican indigenous groups from Central, Southern and South Eastern Mexico. Many indigenous groups within Southern Mexico belong to the Oto-mangue linguistic family (e.g. Mixteco, Zapoteco, Ixcateco) whereas most indigenous groups from South Eastern Mexico belong to the Maya linguistic family (Maya, Chuj, Ixil, Awakateco). Genetic analyses of indigenous groups in Mexico have previously shown that indigenous groups in these regions share extensive genetic similarly that closely aligns with linguistic family membership^2^. On the other hand, indigenous groups in the Central region of Mexico (e.g. Otomi, Nahuatl) show pronounced genetic similarity (i.e. low measures of pairwise Fst) despite spanning distinct linguistic families (e.g. Oto-mangue, Yuto-nahua, Totonaco-tepehua). Our analyses revealed that Mesoamerican ancestry from these three regions was prevalent within the MCPS cohort, with particularly elevated relative proportions of South Eastern ancestry among individuals with the most Native American admixture in-keeping with the more restrictive mating patterns seen in South Eastern Mexican peoples previously ^49^. In contrast, ancestry from Aridoamerican indigenous groups in the Northernmost regions of the country and from Mesoamerican groups in the Northwest state of Nayarit (Cora, Tepehuano, Mexicanero, and Huichol) was underrepresented in MCPS. Moreover, as seen in previous studies in Mexico ^2,46^, we found evidence of sex imbalance on the X chromosome. The higher proportion of Mesoamerican ancestry on chromosome X is consistent with sex-biased gene flow resulting from predominantly-male European colonization of the Americas^50^ and may have implications for health disparities between men and women in light of the longer runs of homozygosity, and more rare pLOF variants, that tracked with Mesoamerican ancestry. Such health disparities may also be compounded by the assortative mating observed in MCPS, which has been well-documented elsewhere^51, 52^. Furthermore, IBD-based analyses revealed extensive and complex patterns of relatedness between participants within Coyoacán and Iztapalapa, largely reflecting the household-based recruitment strategy of the study. Together our analyses have uncovered an exceptionally complex and unique combination of admixture and relatedness within MCPS.

We developed a novel approach for estimating population-specific allele frequencies that leverages local ancestry information and interpolated ancestry at called variants in the MCPS WES and WGS datasets. This dramatically increased (by10-fold) both the number of variants with ancestry-specific allele frequencies and the Native Mexican effective sample size used for estimating allele frequencies from WES data. Without a suitable reference dataset of population specific allele frequencies, efforts to diagnose and interpret genomic variants in the context of rare disorders are greatly encumbered as it is difficult to distinguish previously unreported or undersampled population specific variants from potentially pathogenic variants. Our study expands the availability of such allelic information, which is made accessible to the genomics research community via the MCPS Variant Browser to facilitate future discoveries.

The MCPS WES and WGS datasets substantially add to the global survey of characterized genomic variants by over 31 million variants. Additionally, we uncover elevated levels of homozygosity and homozygous pLOF variants attributable to Native Mexican ancestry, suggesting a role for future studies of admixed Mexicans as a previously untapped resource for the study of homozygous loss of function alleles in humans. Comparing WGS and WES datasets in the same set of 9,950 samples we found that the WGS dataset led to a 2.3% absolute increase in the amount of coding variation when using the canonical gene transcript to annotate variants. Further quantitative comparisons in larger datasets such as UK Biobank will be needed to examine the overall utility of WGS over WES and imputation for novel causal variant discovery.

The MCPS10k imputation reference panel is being made available via the Michigan Imputation Server for use in other studies. From our investigations we found that imputation accuracy with MCPS10k was superior to the TOPMed reference panel at genetic variants with MAF>0.1%, while TOPMed outperformed MCPS10k for the imputation of extremely rare variants in MCPS. We also found that MCPS10K provided the highest imputation accuracy for those individuals with high proportions of Mesoamerican ancestry. In theory, a combination of the MCPS10k and TOPMed reference panels should result in superior imputation performance than using either reference panel alone. There are, however, significant challenges in bringing together large WGS datasets across studies for imputation, motivating the need for novel approaches that can combine imputation results from different panels. The results from our study highlight the need for large diverse WGS datasets from many different populations and the potential for a single world-wide-reference panel to increase representation and parity in imputation accuracy across ancestries.

The publicly-available MCPS genetic resources, particularly the allele frequency and imputation databases, will contribute to future studies and serve as a major resource for understanding the genetic basis of diseases across Mexico as well as in the United States where there is a large population of individuals of Mexican descent. In addition, our study can serve as a blueprint for obtaining novel insight into the complex genetic architecture of other diverse populations. The utility of the MCPS genetic resource has recently been demonstrated through its contribution to the discovery of loss of function variation in *GPR75* that is protective against obesity, which was bolstered by the inclusion of the MCPS cohort in the exome-wide association meta-analysis ^48^. Moreover, the analysis of MCPS exomes was instrumental in estimating that *MC4R* heterozygous deficiency is more than seven times greater in Mexico than in the UK^48^. Future studies will link genetic variation to other disease traits via cross-cohort meta-analysis, increase resolution of fine-mapping, explore the construction and portability of polygenic risk scores in the Mexican population, leverage admixture, relatedness, and household information to potentially boost power of discovery in association studies and utilize Mendelian randomization to uncover causal relationships between modifiable exposures and disease.

## Methods

### Blood sample collection, processing and storage, and DNA extraction

At recruitment, a 10-ml venous EDTA blood sample was obtained from each participant and transferred to a central laboratory using a transport box chilled (4 to 10°C) with ice packs. Samples were refrigerated overnight at 4°C, and then centrifuged (2100g at 4°C for 15 min) and separated the next morning. Plasma and buffy-coat samples were stored locally at −80°C, then transported on dry ice to Oxford (United Kingdom) for long-term storage over liquid nitrogen. DNA was extracted from buffy coat at the UK Biocentre using Perkin Elmer Chemagic 360 systems and suspended in TE buffer. UV-VIS spectroscopy using Trinean DropSense96 was used to determine yield and quality and samples were normalised to provide 2μg DNA at 20ng/μL concentration (2% of samples provided a minimum 1.5ug DNA at 10ng/uL concentration) with 260:280nm ratio of >1.8 and a 260:230nm ratio of 2.0-2.2.

### Exome sample preparation and sequencing and quality control

Genomic DNA samples were transferred to the Regeneron Genetics Center from the UK Biocentre and stored in an automated sample biobank at −80°C prior to sample preparation. DNA libraries were created by enzymatically shearing DNA to a mean fragment size of 200 base pairs, and a common Y-shaped adapter was ligated to all DNA libraries. Unique, asymmetric 10 base pair barcodes were added to the DNA fragment during library amplification to facilitate multiplexed exome capture and sequencing. Equal amounts of sample were pooled prior to overnight exome capture, with a slightly modified version of IDT’s xGenv1 probe library; all samples were captured on the same lot of oligos. The captured DNA was PCR amplified and quantified by qPCR. The multiplexed samples were pooled and then sequenced using 75 base pair paired-end reads with two 10 base pair index reads on the Illumina NovaSeq 6000 platform or S4 flow cells. A total of 146,068 samples were made available for processing. We were unable to process 2,628 samples, most of which failed QC during processing due to low or no DNA being present. A total of 143,440 samples were sequenced. The average 20X coverage is 96.5%, and 98.7% of the samples are above 90%.

Samples showing disagreement between genetically-determined and reported sex (n=1,027), high rates of heterozygosity/contamination (VBID > 5%) (n=249), low sequence coverage (less than 80% of targeted bases achieving 20X coverage) (n=29), or genetically-identified sample duplicates (n=1,062 total samples), and WES variants discordant with genotyping chip (n=8) were excluded. In addition, n=1,339 samples were flagged by MCPS for exclusion. In total, 2,394 unique samples did not pass one or more of our QC metrics. The remaining 141,046 samples were then used to compile a project-level VCF (PVCF) for downstream analysis, using the GLnexus joint genotyping tool. This final dataset contained 9,950,580 variants.

### Whole genome sample preparation and sequencing and quality control

Approximately 250ng of total DNA was enzymatically sheared to a mean fragment size of 350 base pairs. Following ligation of a Y-shaped adapter unique, asymmetric 10 base pair barcodes were added to the DNA fragments with three cycles of PCR. Libraries were quantified by qPCR, pooled, and then sequenced using 150 base pair paired-end reads with two 10 base pair index reads on the Illumina NovaSeq 6000 platform on S4 flow cells. A total of 10,008 samples were sequenced. This included 200 mother-father-child trios and three more extended pedigrees. The rest of the samples were chosen to be unrelated to third-degree or closer, and enriched for parents of nuclear families. The average mean coverage was 38.5X and 99% of samples have mean coverages > 30X, and all samples are above 27X.

Samples showing disagreement between genetically-determined and reported sex (n=16), high rates of heterozygosity/contamination (VBID > 5%) (n=10), or genetically-identified sample duplicates (n=19 total samples) were excluded. In addition, n=34 samples were flagged by MCPS for exclusion. In total, 58 unique samples did not pass one or more of our QC metrics. The remaining 9,950 samples were then used to compile a project-level VCF (PVCF) for downstream analysis, using the GLnexus joint genotyping tool. This final dataset contained 158,464,363 variants.

### Variant calling

The MCPS WES and WGS data were reference-aligned with the OQFE protocol^53^ which employs BWA MEM to map all reads to the GRCh38 reference in an alt-aware manner, marks read duplicates, and adds additional per-read tags. The OQFE protocol retains all reads and original quality scores such that the original FASTQ is completely recoverable from the resulting CRAM file. Single-sample variants are called with DeepVariant v0.10.0 ^54^ using default WGS parameters or custom exome parameters^53^, generating a gVCF for each input OQFE CRAM file. These gVCFs are aggregated and joint-genotyped with GLnexus v1.3.1. All constituent steps of this protocol are executed with open-source software.

### Identification of low-quality variants from sequencing using machine learning

Briefly, we defined a set of positive control and negative control variants based on: (i) concordance in genotype calls between array and exome sequencing data; (ii) transmitted singletons; (iii) an external set of likely “high quality” sites; and (iv) an external set of likely “low quality” sites. To define the external “high quality” set, we first generated the intersection of variants that pass QC in both TOPMED Freeze 8 and gnomAD v3.1 genomes. This set was additionally restricted to 1000 genomes phase 1 high-confidence SNPs ^34^ and Mills and 1000 genomes gold-standard INDELs^55^, both available via GATK resource bundle (**URLs**). To define the external “low quality” set, we intersected gnomAD v3.1 fail variants with TOPMED Freeze 8 mendelian or duplicate discordant variants. Prior to model training, the control set of variants were binned by allele frequency, and then randomly sampled such that an equal number of variants were retained in the positive and negative labels across each frequency bin. The model was then trained on up to 33 available site quality metrics, including, for example, the median value for allele balance in heterozygote calls and whether a variant was split from a multi-allelic site. We split the data into training (80%) and test (20%) sets. We performed a grid search with 5-fold cross-validation on the training set to identify the hyperparameters that return the highest accuracy during cross-validation, which are then applied to the test set to confirm accuracy. This approach identified as low-quality a total of 616,027 WES and 22,784,296 WGS variants (of which 161,707 and 104,452 were coding variants respectively). We further applied a set of hard filters to exclude monomorphs, unresolved duplicates, variants with >10% missingness, ≥3 mendel errors (WGS only), or failed HWE with excess heterozgosity (HWE p-value < 1×10^-30^ and observed heterozygote count > 1.5x expected heterozygote count), resulting in a dataset of 9,325,897 WES and 131,851,586 WGS variants (of which 4,037,949 and 1,460,499 were coding variants respectively).

### Variant annotation

Variants were annotated as previously described. Briefly, variants were annotated using Ensembl Variant Effect Predictor (VEP)^56^, with the most severe consequence for each variant chosen across all protein coding transcripts. In addition, we also derived canonical transcript annotations based on a combination of MANE, APPRIS and Ensembl canonical tags. MANE annotation is given the highest priority, followed by APPRIS. When neither MANE nor APPRIS annotation tags are available for a gene, the canonical transcript definition of Ensembl is used. Gene regions were defined using Ensembl Release 100. Variants annotated as stop gained, start lost, splice donor, splice acceptor, stop lost or frameshift, for which the allele of interest is not the ancestral allele, are considered predicted LOF variants. Five annotation resources were utilized to assign deleteriousness to missense variants: SIFT^57^; PolyPhen2 HDIV and PolyPhen2 HVAR^58^; LRT ^59^; and MutationTaster ^60^. Missense variants were considered “likely deleterious” if predicted deleterious by all five algorithms, “possibly deleterious” if predicted deleterious by at least one algorithm, and “likely benign” if not predicted deleterious by any algorithm.

### Genotyping

Samples were genotyped on the Illumina Global Screening Array (GSA) v2 beadchip according to the manufacturer’s recommendations. A total of 146,068 samples were made available for processing, of which 145,266 (99.5%) were successfully processed. The average genotype call rate per sample was 98.4% and 98.4% of samples had a call rate above 90%. Samples showing disagreement between genetically-determined and reported sex (n=1,827), low quality samples (call rates below 90%) (n=2,276), genotyping chip variants discordant with exome data (n=44), genetically-identified sample duplicates (n=1,063 total samples) were excluded. In addition, n=1,122 samples were flagged by MCPS for exclusion and n=223 samples were failed for “other” reasons. In total, 4,435 unique samples did not pass one or more of our QC metrics. The remaining 140,831 samples were then used to compile a project-level VCF (PVCF) for downstream analysis. This dataset contained 650,380 poly-morphic variants.

### Genotyping QC

The input array data from the RGC Sequencing Lab consisted of 140,831 samples and 650,380 variants and were passed through multiple quality control steps: checks for consistency of genotypes in sex chromosomes (steps 1-4); sample- and variant-level missingness filters (steps 5-6); the Hardy-Weinberg equilibrium exact test applied to a set of 81,747 3rd-degree unrelated samples, identified from the initial relatedness analysis by PLINK and PRIMUS (step 7); setting genotypes with Mendel errors in nuclear families to missing (step 8); and the second round of steps 5-7 (step 9). PLINK commands associated with each step are displayed in column 2 (**Supplementary Table 9**). The final post-QC array data consisted of 138,511 and 559,923 variants.

### Array phasing

We used SHAPEIT v4.1.3 ^61^ to phase the array dataset of 138,511 samples and 539,315 autosomal variants that passed the array QC procedure. To improve the phasing quality, we leveraged the inferred family information by building a partial haplotype scaffold on unphased genotypes at 1,266 trios from 3,475 inferred nuclear families identified (randomly selecting one offspring per family when there were more than 1). We then ran SHAPEIT one chromosome at a time, passing the scaffold information with the --scaffold option.

### Exome and whole genome phasing

We separately phased the SVM filtered exome and whole genome sequencing datasets onto the array scaffold. For the WGS phasing we used WhatsHap^62^ to extract phase information in the sequence reads and from the subset of available trios and pedigrees, and this information was fed into SHAPEIT v4.2.2 via the --use-PS 0.0001 option. Phasing was carried out in chunks of 10,000 and 100,000 variants (WES and WGS respectively) and using 500 SNPs from the array data as a buffer at the beginning and end of each chunk. The use of the phased scaffold of array variants means that chunks of phased sequencing data can be concatenated together to produce whole chromosome files that preserve the chromosome-wide phasing of array variants. A consequence of this process is that when a variant appears in both the array and sequencing datasets, it is the data from the array dataset that is used.

To assess the performance of the WGS phasing process, we repeated the phasing of chromosome 2 by removing the children of the 200 mother-father-child trios. We then compared the phase of the trio parents to that in the phased dataset that included the children. We observed a mean switch error rate of 0.0024. Without using the WhatsHap to leverage phase information in sequencing reads increases the mean switch error rate to 0.0040 (**Supplementary Figure 22**).

### Relatedness, pedigree reconstruction and network visualization

The relatedness inference criteria and relationship assignments were based on kinship coefficients and probability of zero IBD sharing from the KING software^26^. We reconstructed all first-degree family networks using PRIMUSv1.9.0 ^30^ applied to the IBD-based KING estimates of relatedness along with the genetically derived sex and reported age of each individual. 99.3% of the first-degree family networks were reconstructed unambiguously. To visualize the relationship structure in MCPS we used the Graphviz software (see **URLs**) to construct networks such as **Supplementary Figure 5**. We used the sfdp layout engine which uses a “spring” model that relies on a force-directed approach to minimize edge length.

### Measuring IBD segments and homozygosity

To identify IBD segments and measure runs of homozygosity, we ran hap-ibd (v1.0) ^27^ using the phased array dataset of 138,511 samples and 538,614 sites from autosomal loci. Hap-ibd was run with the parameter min-seed=4, which looks for IBD segments that are at least 4 cM long. We filtered out IBD segments in regions of the genome with fourfold more or fourfold less than the median coverage along each chromosome following the procedure in IBDkin^28^, and filtered out segments overlapping regions with fourfold less than the median SNP marker density (**Supplementary Figure 30**). For the homozygosity analysis, we intersected the sample with the exome data to evaluate loss of function variants, resulting in a sample of 138,200. We further overlaid the ROH segments with local ancestry estimates, and assigned ancestry where the ancestries were concordant between haplotypes and posterior probability was >0.9, assigning ancestry to 99.8% of the ROH.

### Principal Components Analysis

We used the workflow described in Privé et al. 2020^63^ and implemented in the R package *bigsnpr*. In brief, pairwise kinship coefficients are estimated with *Plink* (v2.0) and samples were pruned for first and second-degree relatedness (kinship coefficient < 0.0884) to obtain a set of unrelated individuals. Linkage disequilibrium (LD) clumping was performed with a default LD *r^2^* threshold of 0.2 and regions with long-range LD were iteratively detected and removed using a procedure based on evaluating robust Mahalanobis distances of PC loadings. Sample outliers are detected using a procedure based on K nearest neighbours. Principal component (PC) scores and loadings for the first 20 PCs are efficiently estimated using *truncated* singular value decomposition (SVD) of the scaled genotype matrix. After removal of variant and sample outliers, a final iteration of truncated SVD was performed to obtain the PCA model. The PC scores and loadings from this model were then used to project withheld samples, including related individuals, into the PC space defined by the model using the Online Augmentation, Decomposition and Procustes (OADP) algorithm. For each PC analysis in this study, variants with minor allele frequency (MAF) < 0.01 were removed.

### ADMIXTURE analysis

ADMIXTURE^64^ version 1.3.0 (**URLs**) was used to estimate ancestry proportions in a set of 3,964 reference samples representing African, European, East Asian, and American ancestries from a dataset of merged genotypes. 765 samples of African ancestry were obtained from 1KG (n=661) and HGDP (n=104), 658 samples of European ancestry were obtained from 1KG (n=503) and HGDP (n=155), and 727 samples of East Asian ancestry were obtained from 1KG (n=504) and HGDP (n=223). American samples were limited 716 indigenous Mexican samples from the MAIS study, 64 admixed Mexican American samples from Los Angeles from 1KG (MXL), 21 Maya and 13 Pima samples from HGDP, and 1,000 unrelated Mexican samples from MCPS. Included SNPs were limited to variants present on the Illumina GSAv2 genotyping array for which TOPMED-imputed variants in the MAIS study had info *r^2^* ≥ 0.9 (m=199,247 SNPs). To select the optimum number of ancestry groups (K) to include in the admixture model, five-fold cross validation was performed for each K in the set 4 to 25 with the –cv flag. In order to obtain ancestry proportion estimates in the remaining set of 137,511 MCPS samples, the population allele frequencies (P) estimated from the analysis of reference samples were fixed as parameters so that the remaining samples could be projected into the admixture model. Projection was performed for the K=4 model and for the K=18 model that yielded the lowest cross validation error, and point estimation was attained with the block relaxation algorithm.

### External datasets used in genetic analyses

The Metabolic Analysis of an Indigenous Sample (MAIS) genotyping datasets were obtained from Professor Lorena Orozco from Insituto Nacional de Medicina Genómica (INMEGEN). For 644 samples, genotyping was performed using the Affymetrix Human 6.0 array (n=599,727 variants). An additional 72 samples (11 ancestry groups) were genotyped with the Illumina Omni 2.5 array (n=2,397,901 variants). The set of 716 indigenous samples represent 60 of the 68 recognised ethnic groups in Mexico^2^. Per chromosome variant call files (VCF) for each genotyping array were uploaded to the TOPMed Imputation Server (**URLs**) and imputed from a multi-ethnic reference panel of 97,256 whole genomes. Phasing and imputation were performed using the programs *eagle* and *MiniMac*, respectively. The observed coefficient of determination (*r^2^*) for the reference allele frequency between the reference panel and the genotyping array was 0.696 and 0.606 for the Affymetrix and Illumina arrays, respectively.

Physical positions of imputed variants were mapped from genome build GRCh37 to GRCh38 using the program *LiftOver* and only variant positions included on the Affymetrix Global Screening Array version 2 (GSAv2) were retained. After further filtering out variants with imputation info *r^2^* < 0.9, the following quality control steps were performed prior to merging of the MAIS Affymetrix and Illumina datasets: 1) removal of ambiguous variants (i.e. A/T and C/G polymorphisms); 2) removal of duplicate variants; 3) identifying and correcting allele flips; 4) removal of variants with position mismatches. Merging was performed with the -- bmerge command in *Plink* (v1.9).

We used publicly available genotypes from the Human Genome Diversity Panel^65^ HDPG; n=929) and 1000 Genomes Project ^66^ (KG; n=2,504). To obtain a combined global reference dataset for downstream analyses of population structure, admixture, and local ancestry, the HGDP and KG datasets were merged. The resulting merged public reference dataset was subsequently merged with the MAIS dataset and MCPS genotyping array dataset. Each merge was performed with the –bmerge function in Plink (v1.9) after removing ambiguous variants, removing duplicate variants, identifying and correcting allele flips, and removing variants with position mismatches. The combined global reference dataset comprised 199,247 variants and 142,660 samples.

### Local Ancestry Inference

Reference samples used in the LAI were selected from a TeraStructure ancestry analysis of the HGDP, 1000G and MAIS samples. A continental ancestry score threshold ≥ 0.90 was applied to exclude samples with extensive admixture from the reference set. The seven group analysis used 666 African samples, 659 European samples, 98 Mexico-North, 42 Mexico-Northwest, 185 Mexico-Central, 128 Mexico-South and 163 Mexico-Southeastern samples at 204,626 autosomal and chromosome X SNPs. The 3 group analysis used 666 African samples, 659 European samples and 163 Native Mexican samples at 505,834 autosomal and chromosome X SNPs. The reference samples were phased with *SHAPEIT* v4.1.2 using default settings. RFMix was applied to 138,511 MCPS samples in the phased array dataset. Selected RFMix v2 parameters (-e 5 --reanalyze-reference -n 5 -G 15) tuned the LAI inference algorithm to perform 5 rounds of Expectation-Maximization (EM) with an option to treat reference haplotypes if they were query haplotypes and update the set of reference haplotypes in each EM round; the average number of generations since expected admixture was set to 15; and the number of terminal nodes for the random forest classifier was set to 5.

### Fine-scale population structure based on IBD sharing

IBD segments from hap-ibd were summed across pairs of individuals to create a network of IBD sharing represented by weight matrix *W* ∈ ℝ_≥0_^n×n^ for *n* samples. Each entry *W*_ij_ ∈ *W* gives the total length in cM of the genome that individuals *i* and *j* share identical by descent. We sought to create a low-dimensional visualization of the IBD network. We took an approach similar to Han et al. 2017 ^35^, who use the eigenvectors of the normalized graph Laplacian as coordinates for a low-dimensional embedding of the IBD network. Let *D* be the degree matrix of the graph with *d*_ii_ = ∑_j_ *W*_ij_ and 0 elsewhere. The normalized (random walk) graph Laplacian is defined to be L = I − *D*^−1^*W*, where *I* is the identity matrix.

The matrix *L* is positive semi-definite, with eigenvalues 0 = λ_0_ ≤ λ_1_ ≤ ⋯ ≤ λ_*n*−1_. The multiplicity of eigenvalue 0 is determined by the number of connected components in the IBD network. If *L* is fully connected, the eigenvector associated with eigenvalue 0 is constant, while the remaining eigenvectors can be used to compute a low-dimensional representation of the IBD network. If *p* is the desired dimension, and *u*_1_, …, *u*_*p*_ the bottom 1 … *p* eigenvectors of *L* (indexed from 0), the matrix *U* ∈ ℝ^*n* ×*p*^ with columns *u*_1_, …, *u*_*p*_ define a low-dimensional representation of each individual in the IBD network ^67^.^67^. In practice, we solve the generalized eigenvalue problem to obtain *u*_1_, …, *u*_*p*_.

*Wu* = μ*Du*

If u is an eigenvector of *L* with eigenvalue λ, then *u* solves the generalized eigenvalue problem with eigenvalue 1 − λ.

To apply to the IBD network of the MCPS cohort, we first removed edges with weight >72 cM following Han et al. 2017. We did this to avoid the influence on extended families on the visualization. We next extracted the largest connected component from the IBD network, and computed the bottom *u*_1_, …, *u*_20_ eigenvectors of the normalized graph Laplacian.

### Fine-scale population structure based on haplotype sharing

To examine fine-scale population structure using haplotype sharing we calculated a haplotype copying matrix *L* using IMPUTE5 ^6868^ with entries *Lij* that are the length of sequence individual *i* copies from individual *j*. IMPUTE5 employs a scalable imputation method that can handle very large haplotype reference panels. At its core is an efficient HMM that can estimate the local haplotype sharing profile of a ‘target’ haplotype with respect to a ‘reference’ set of haplotypes. To avoid the costly computations of using all the reference haplotypes, an approach based on the PBWT data structure is used to identify a subset of reference haplotypes that leads to negligible loss of accuracy. We leveraged this methodology to calculate the copying matrix *L*, using array haplotypes from a set of 58,329 unrelated individuals as *both* target and reference datasets, and used the --ohapcopy –ban-repeated-sample-names flags to ban each target haplotype being able to copy itself. SVD on a scaled centred matrix was performed using the bigstatsr package ^63^ to generate 20 PCs. This is equivalent to an eigen-decomposition of the variance-covariance matrix of recipients’ shared segment lengths.

### Imputation experiments

We imputed the filtered array dataset using both the MCPS10k reference panel and the TOPMed imputation server. For TOPMed imputation we used Plink2 to convert this dataset from Plink1.9 format genotypes to unphased VCF genotypes. For compatibility with TOPMed imputation server restrictions, we split the samples in this dataset into six randomly assigned subsets of about 23,471 samples, and also into chromosome specific bgzipped VCF files. Using the NIH Biocatalyst API (see **URLs**) we submitted these six jobs to the TOPMed imputation server. Upon completion of all jobs, we used bcftools merge to join the resulting dosage VCFs spanning all samples. For the MCPS10k imputation we used IMPUTE 5 v1.1.5. Each chromosome was split into chunks using the imp5Chunker program with minimum window size 5Mb, and minimum buffer size 500Kb. Information scores were calculated using qctool (**URL**s).

The 1000 Genomes WGS dataset was downloaded (**URLs**) and filtered to remove sites that are multi-allelic sites, duplicated, have missingness >2%, Hardy-Weinberg p-value < 1e^-8^ in any sub-population, and MAF<0.1% in any sub-population. We used only those 490 American (AMR) samples in the MXL, CLM, PUR and PEL sub-populations. We constructed 2 subsets of genotypes on chromosome 2 from the Illumina HumanOmniExpressExome (8v1-2) and Illumina GSA (v2) arrays, and these used as input to the TOPMed and MCPS10k imputation pipelines.

We measured imputation accuracy by comparing the imputed dosage genotypes to the true (masked) genotypes at variants not on the arrays. Markers were binned according to the MAF of the marker in 490 AMR samples. In each bin, we report the squared correlation (*r*^2^) between the concatenated vector of all the true (masked) genotypes at markers and the vector of all imputed dosages at the same markers.

### Ancestry specific allele frequency estimation

The LAI results consist of segments of inferred ancestry across each haplotype of the phased array dataset. Since the WES and WGS alleles were phased onto the phased array scaffold we inferred the ancestry of each exome allele using interpolation from the ancestry of the flanking array sites. For each WES and WGS variant on each phased haplotype we determined the RFMix ancestry probability estimates at the two flanking array sites and used those to interpolate their ancestry probabilities. Ancestry specific frequencies are then calculated from the weighted allele counts and summed ancestry probabilities. Singleton sites can be hard to phase using existing methods. Family information and phase information in sequencing reads was used in the WGS phasing, and this will have helped to phase a proportion of the singleton sites. In the WES dataset we found that 46% of exome singletons occurred in stretches of heterozygous ancestry. For these variants we gave equal weight to the two ancestries when estimating allele frequencies.

To validate the MCPS allele frequencies we downloaded the gnomAD v3.1 reference dataset (see **URLs**) and retained only high-quality variants annotated as passed QC (FILTER=”PASS”), SNVs, outside low-complexity regions and with the number of called samples greater than 50% of the total sample size (N = 76,156). We additionally overlapped gnomAD variants with TOPMed Freeze 8 high-quality variants (FILTER=”PASS”) (see **URLs**). We further merged gnomAD variants and MCPS exome variants by the C:P:R:A (chromosome:position: reference allele:alternative allele) names and excluded MCPS singletons, which were heterozygous in ancestry. That resulted in 2,249,986 overlapping variants available for comparison with the MCPS WES data. Median sample sizes in gnomAD non-Finish Europeans, African/Admixed African and Admixed American population groups were N = 34,014, 20,719 and 7,639 respectively.

## URLs

MCPS Allele Frequency browser https://rgc-mcps.regeneron.com/

SHAPEIT https://odelaneau.github.io/shapeit4/

QCTOOL https://www.well.ox.ac.uk/~gav/qctool_v2/

MakeScaffold https://github.com/odelaneau/makeScaffold

Hap-IBD https://github.com/browning-lab/hap-ibd

IMPUTE5 https://jmarchini.org/software/#impute-5

MICHIGAN imputation server https://imputationserver.sph.umich.edu/

gnomAD https://gnomad.broadinstitute.org

TOPMed Freeze 8 BRAVO variant browser, https://bravo.sph.umich.edu/freeze8/hg38/

TOPMed imputation server https://imputation.biodatacatalyst.nhlbi.nih.gov

Million Veteran Program https://www.research.va.gov/mvp/

PRIMUS https://primus.gs.washington.edu/primusweb/

GRAPHVIZ https://graphviz.org/

GATK resource bundle: https://gatk.broadinstitute.org/hc/en-us/articles/360035890811-Resource-bundle

ADMIXTURE https://dalexander.github.io/admixture/

1000 Genomes WGS http://ftp.1000genomes.ebi.ac.uk/vol1/ftp/data_collections/1000G_2504_high_coverage/working/20201028_3202_phased/

## Data availability

The MCPS investigators welcome requests from researchers who wish to access data from the Mexico City Prospective Study. If you are interested in obtaining data from the study for research purposes, or in collaborating with MCPS investigators on a specific research proposal, please visit our study website [http://www.ctsu.ox.ac.uk/research/prospective-blood-based-study-of-150-000-individuals-in-mexico] where you can download the study’s Data and Sample Access Policy in English or Spanish. The MCPS10k imputation reference panel described in this manuscript can be used freely for imputation through the University of Michigan Imputation server (see **URLs**).

## Acknowledgements

The Mexico City Prospective Study has received funding from the Mexican Health Ministry, the National Council of Science and Technology for Mexico, the Wellcome Trust, Cancer Research UK, British Heart Foundation and the UK Medical Research Council. These funding sources had no role in the design, conduct or analysis of the study or the decision to submit the manuscript for publication. Genotyping, exome sequencing and whole genome sequencing was funded through an academic partnership between the National Autonomous University of Mexico, the University of Oxford, Regeneron, AstraZeneca and Abbvie. The computational aspects of this research were supported by the Wellcome Trust Core Award Grant Number 203141/Z/16/Z and the NIHR Oxford BRC. The views expressed are those of the authors and not necessarily those of the NHS, the NIHR or the UK Department of Health. The authors are grateful to all the MCPS participants, without whom this research would not be possible. The authors are grateful to Claudia Gonzaga-Jauregui, Yongtao Guan, Brian Browning, Ying Zhou and Kelsey Grinde for discussions and input on various aspects of this work.

## Rights retention statement

For the purpose of open access, the authors have applied a Creative Commons Attribution (CC BY) licence to any Author Accepted Manuscript version arising.

## Ethics approval

Approval for the study was given by the Mexican Ministry of Health, the Mexican National Council of Science and Technology (0595 P-M) and the Central Oxford Research Ethics Committee (C99.260) and the Ethics and Research commissions from the Medicine Faculty at the National Autonomous University of Mexico (UNAM) (FMED/CI/SPLR/067/2015). All study participants provided written informed consent.

## AUTHOR CONTRIBUTIONS

Conceptualization: J.Marchini, A.B, G.R.A, J.R.E, J.A-D, P.K-M, R.T-C

Data curation: R.W

Data generation: M.H, X.B, S.B, W.S, J.D.O, J.G.R, L.O-O, H.G-O

Formal Analysis: A.Z, J.T, J.D.B, J.Mbatchou, M.T. T.T, S.G, T.J, Y.Z, D.L, R.W, J.S, R.P, A.P, K.S, A.N

Funding acquisition: J.A.D, A.B, R.C, J.R.E, S.P, P.K-M, R.T-C

Methodology: A.Z, J.T, J.D.B, T.T, J.Marchini

Project administration: M.J Resources: L.H, R.L, E.M, S.Z

Software: A.Z, J.T, M.T, J.Mbatchou, S.G, T.J, J.S, Y.Z, J.Marchini

Supervision: J.Marchini, J.R.E, T.T, G.A, W.S, J.G.R, E.J, J.A-D, J.B, S.P

Visualization: A.Z, J.T, J.Mbatchou, S.G, J.D.B, T.J, M.T, Y.Z, J.S

Writing first draft: J.Marchini, J.T, J.R.E, T.T

Revision of manuscript: All authors

## Supplementary Note

### History and socio-demographics of Mexico City

The difference in genetic ancestry identified between the inhabitants of Coyoacán and Iztapalapa has a historical correlation. The Mexico City districts of Coyoacán and Iztapalapa have existed since the pre-Hispanic times when they were relatively close (particularly Coyoacán) to the great city of Tenochtitlan. Although the indigenous populations settled in those places were the initial settlements, the population dynamics changed substantially over time, starting with the arrival of the Spaniards. Many Spaniards, including the conqueror Hernán Cortés, resided in Coyoacán while the capital of New Spain was being built (currently the historic center of the CDMX) over the ruins of Tenochtitlan. However, the modern populations of Coyoacán and Iztapalapa derive largely from the development of urban settlements and migrations that occurred from the 1950s to the 1970s. During this period of the twentieth century both districts, but particularly Iztapalapa, received large numbers of indigenous migrants from the Central (Nahuas, Otomies, Purepechas), South (Mixtecos, Zapotecos, Mazatecos), and Southeast (Chinantecos, Totonacas and Mayas) of the country. Today, Coyoacán houses a wide range of cultural and educational spaces and includes many middle and upper-class neighborhoods where those with more significant purchasing power, including many foreigners and Mexican mestizos with more European ancestry, have settled. Iztapalapa, further from the city center and with fewer cultural areas, is more affordable and remains popular among indigenous populations and those who migrate to Mexico City from rural parts of Mexico.

