## Supplementary Figures for "Genotyping, sequencing and analysis of 140,000 adults from the Mexico City Prospective Study"

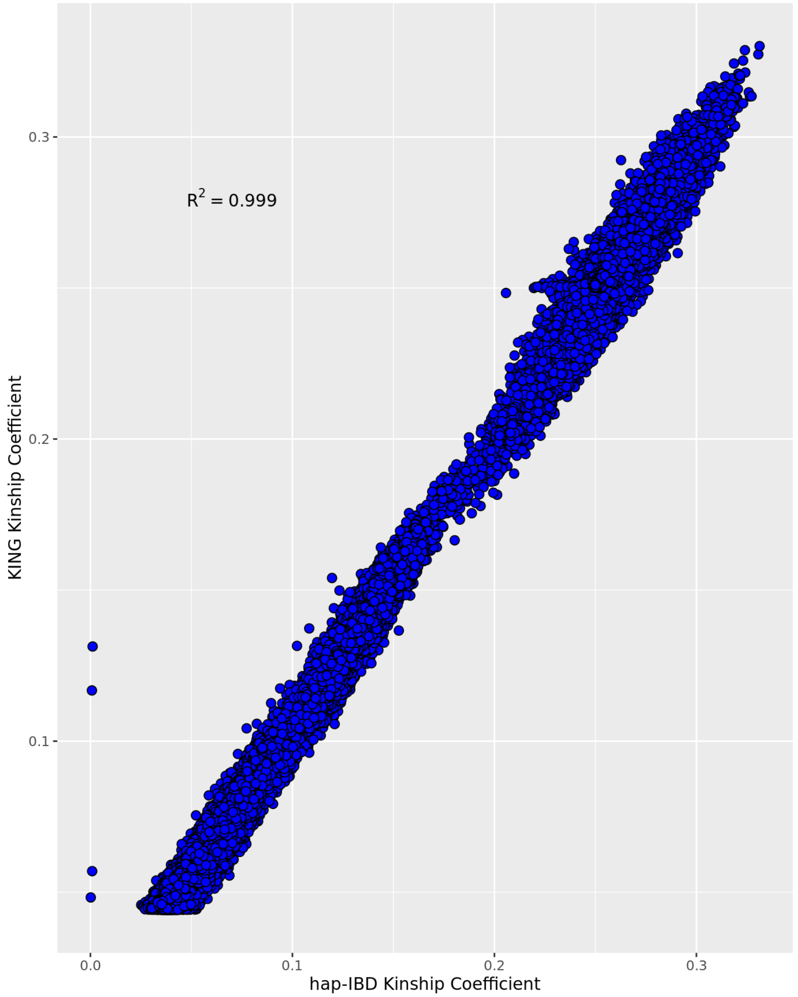


**Supplementary Figure 1:** Scatterplot of pairwise kinship estimates from IBD segments from the KING software and hap-IBD.

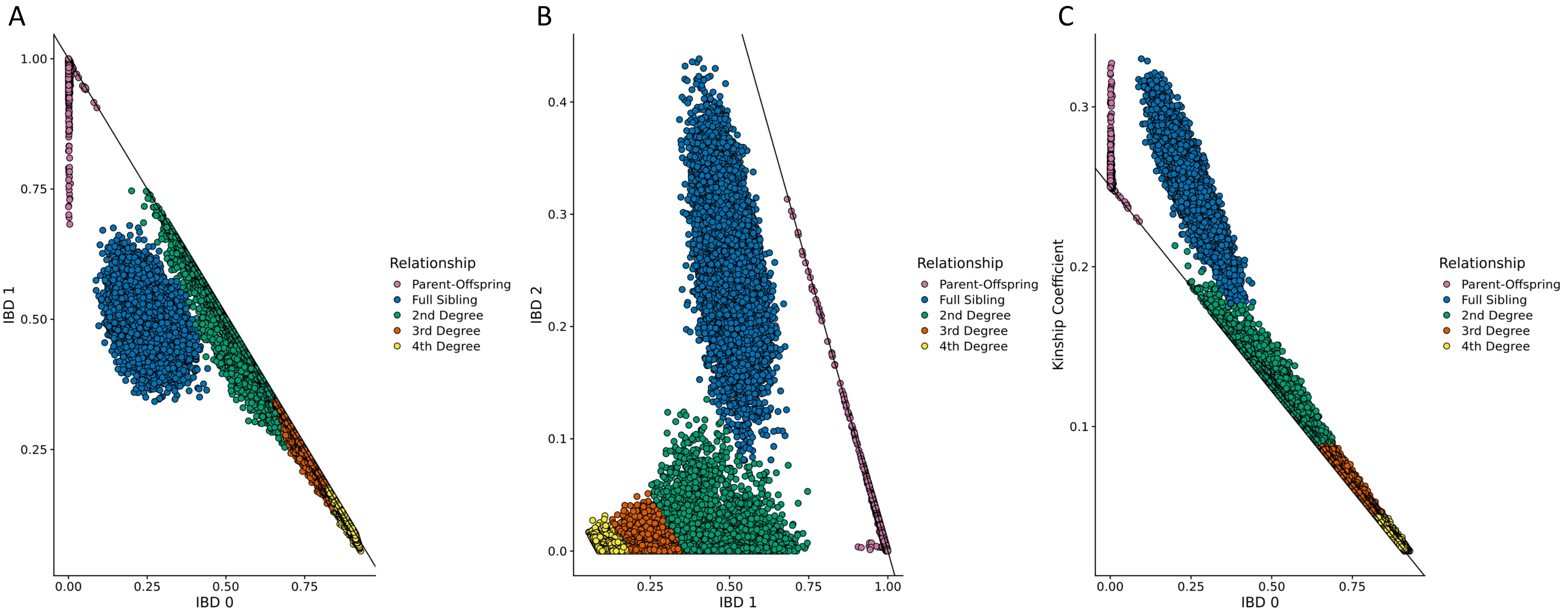


**Supplementary Figure 2: Pairwise measures of relatedness**. (A) IBD0 vs IBD1, (B) IBD1 vs IBD2, and (C) IBD0 vs Kinship coefficient.

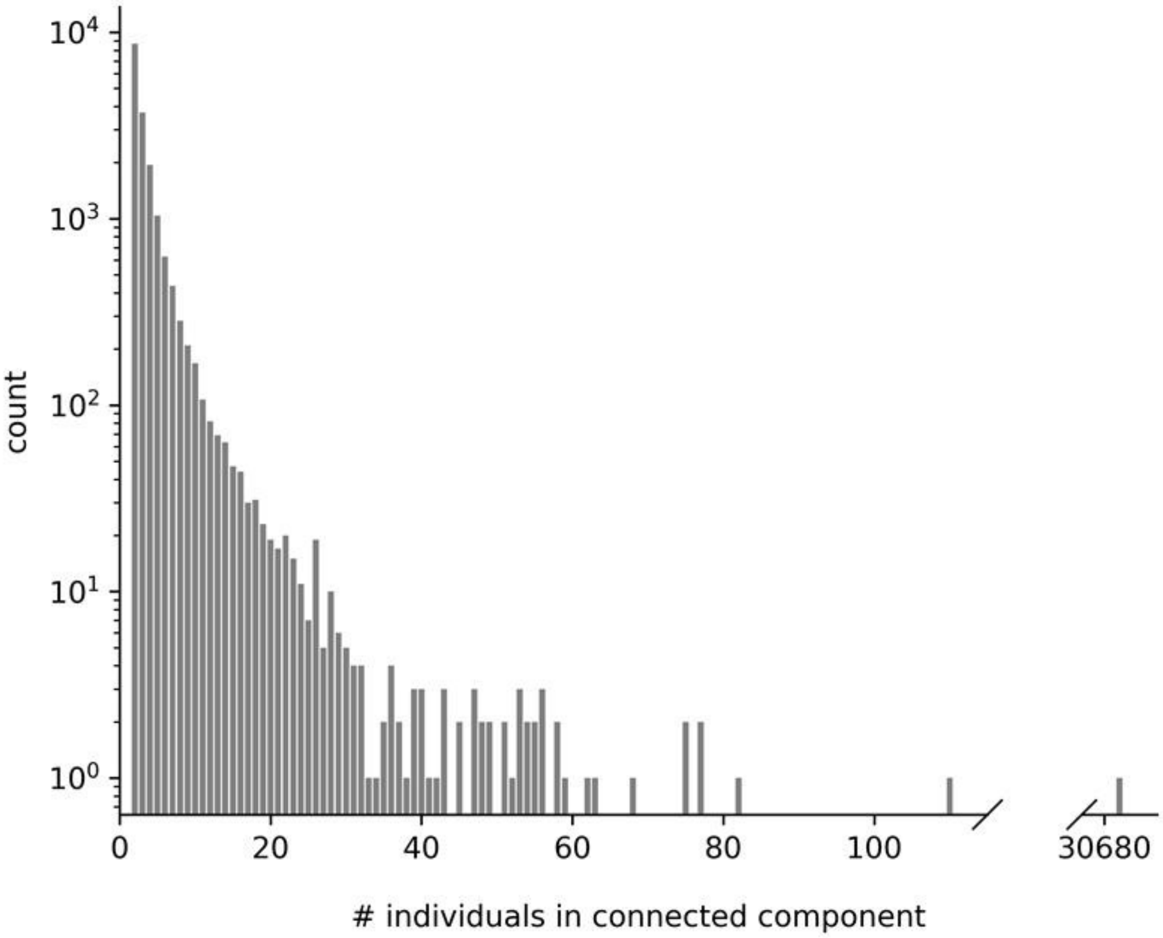


**Supplementary Figure 3**: **Histogram of size of connected components in the 3rd degree relatedness graph**. The largest connected component has size 30,680 and is shown on the histogram using a split axis.

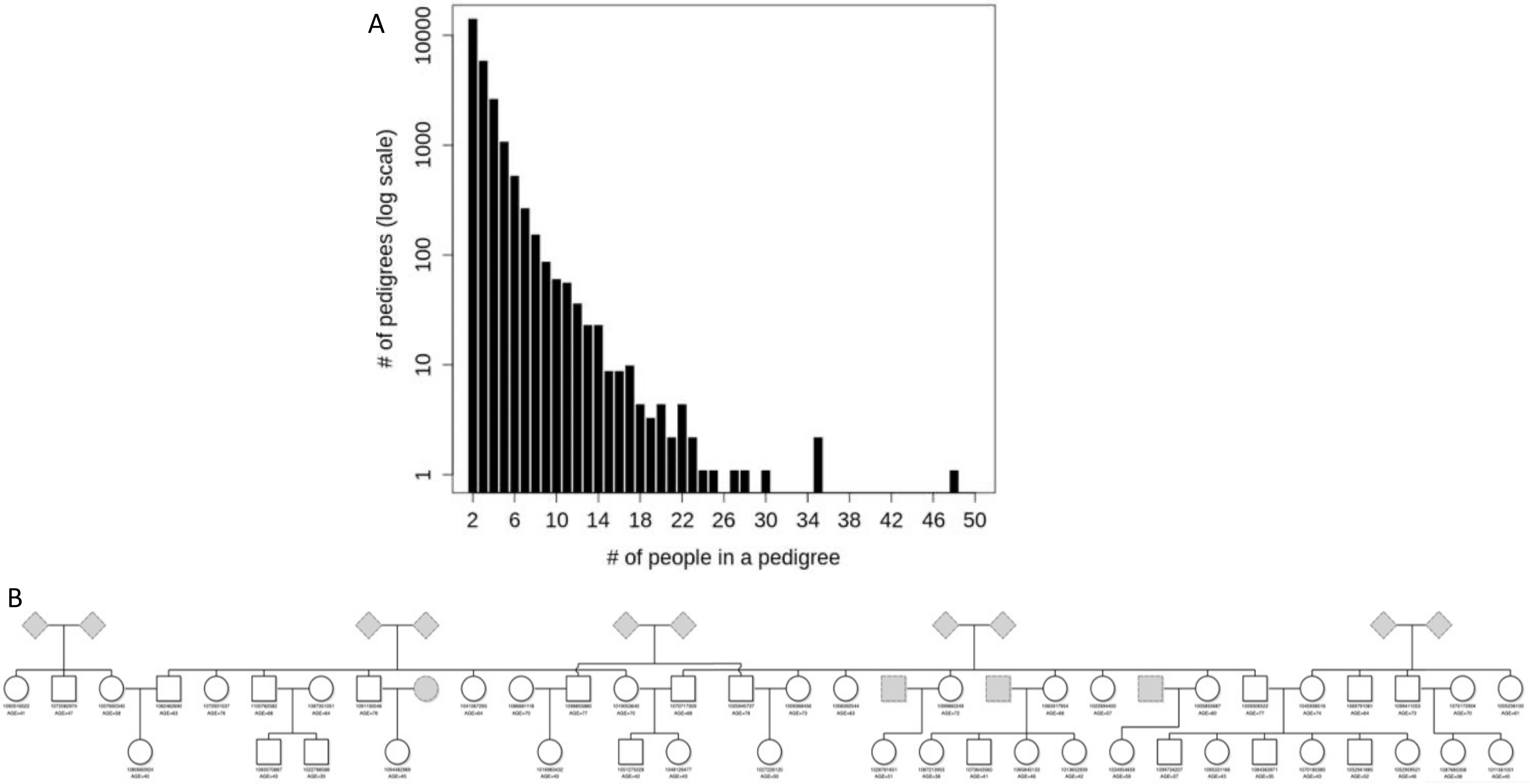


**Supplementary Figure 4: Summary of first-degree family networks.** (A) distribution of family network sizes and (B) largest first-degree family network of 48 individuals.

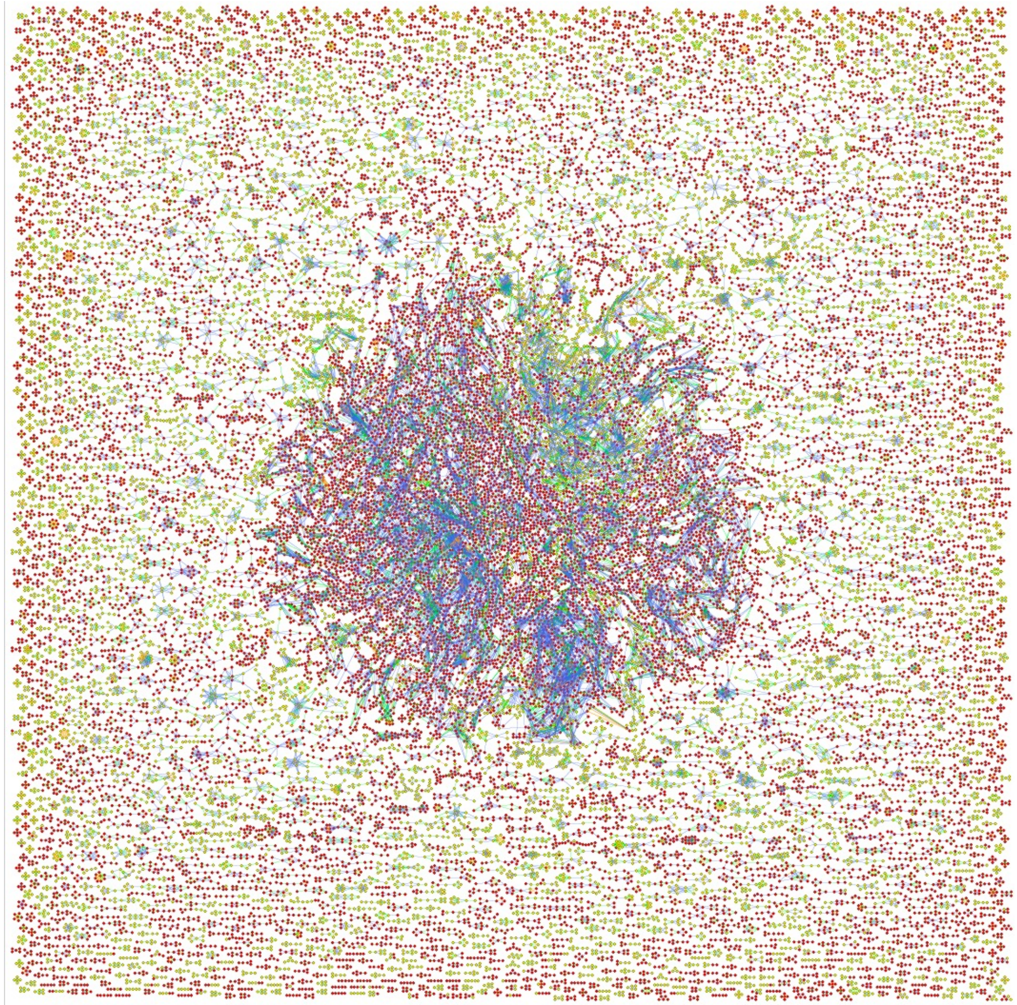

Nodes are individuals
Red = Iztapalapa
Yellow = Coyoacan

Edges are relationships
Green = Parent/child
Orange = Full-sibling
Blue = 2nd degree

**Supplementary Figure 5: Graph of second-degree family networks of size four or greater**. Plot created using used the Graphviz software with the sfdp layout engine which uses a “spring” model that relies on a force-directed approach to minimize edge length.

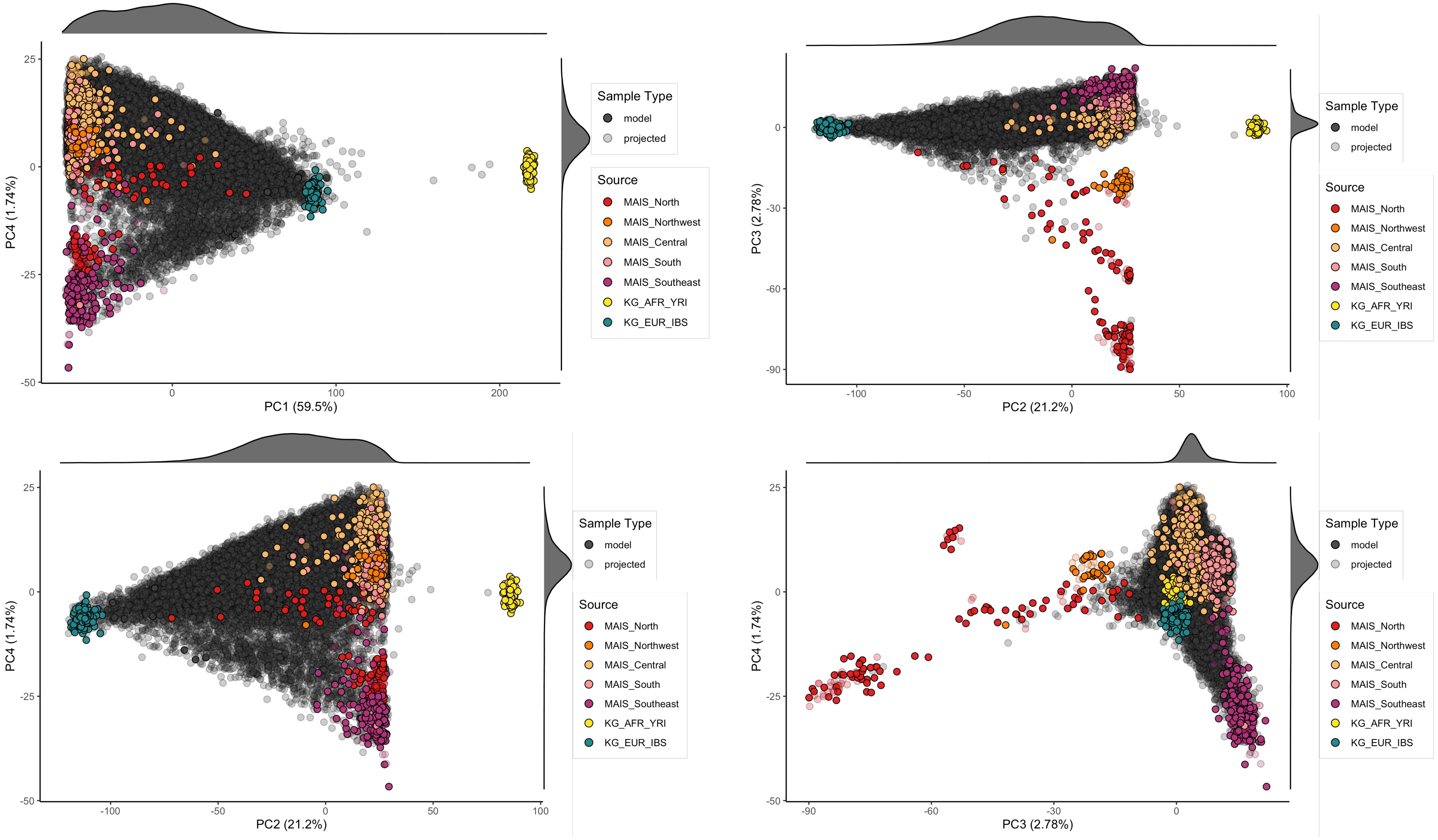


**Supplementary Figure 6: Selected PC scatterplots from a PCA of 500 MCPS samples**. Remaining MCPS samples and reference samples from the 1000 Genomes Project (Yoruba – YRI; Iberian – IBS) and MAIS (indigenous samples from Mexico) projected onto the PC axes. Main Figure 2 shows PC1 vs PC2 and PC1 vs PC3.

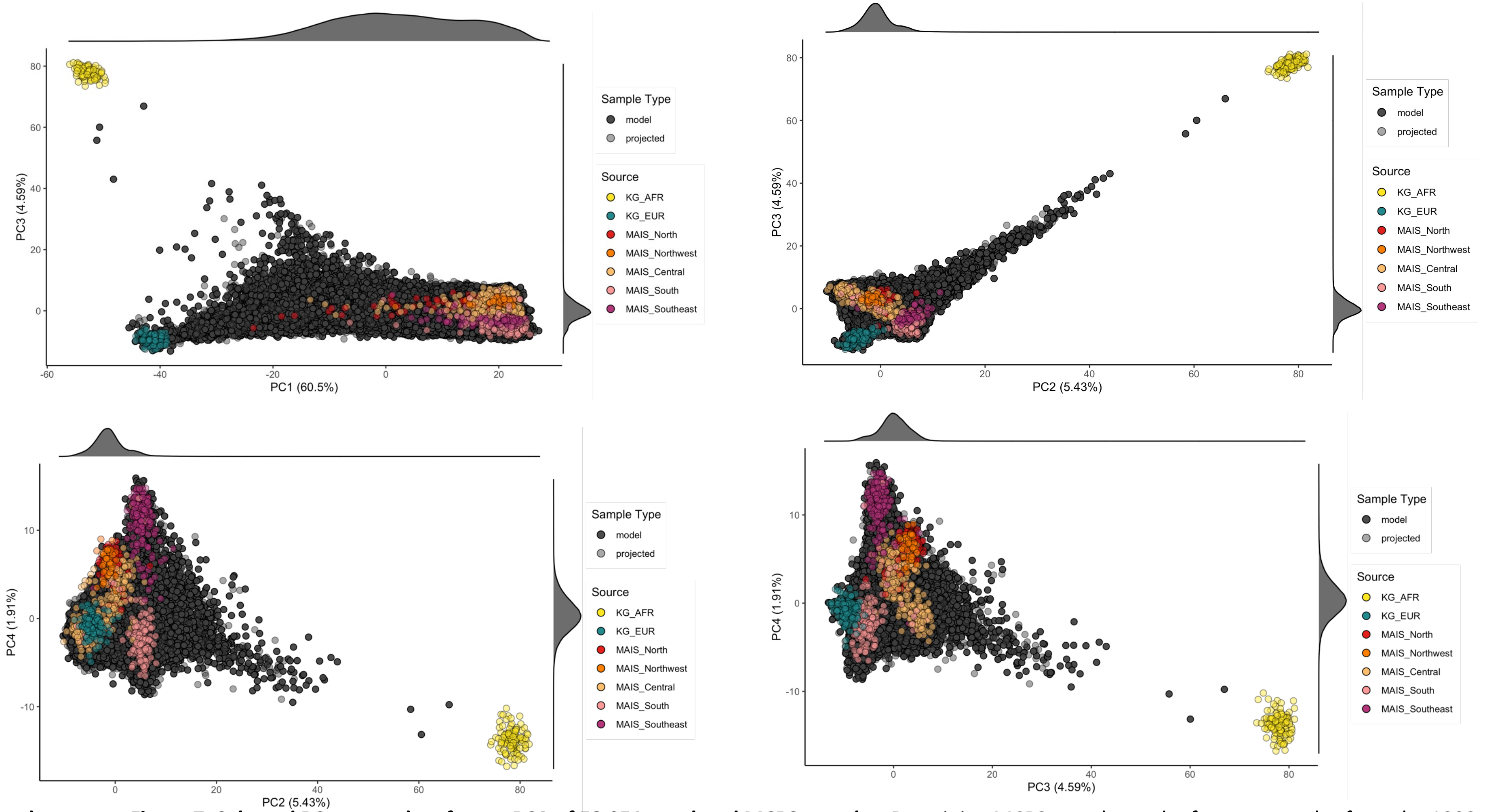


**Supplementary Figure 7: Selected PC scatterplots from a PCA of 58,051 unrelated MCPS samples**. Remaining MCPS samples and reference samples from the 1000 Genomes Project (Yoruba – YRI; Iberian – IBS) and MAIS (indigenous samples from Mexico) projected onto the PC axes. Main Figure 2 shows PC1 vs PC2 and PC1 vs PC4.

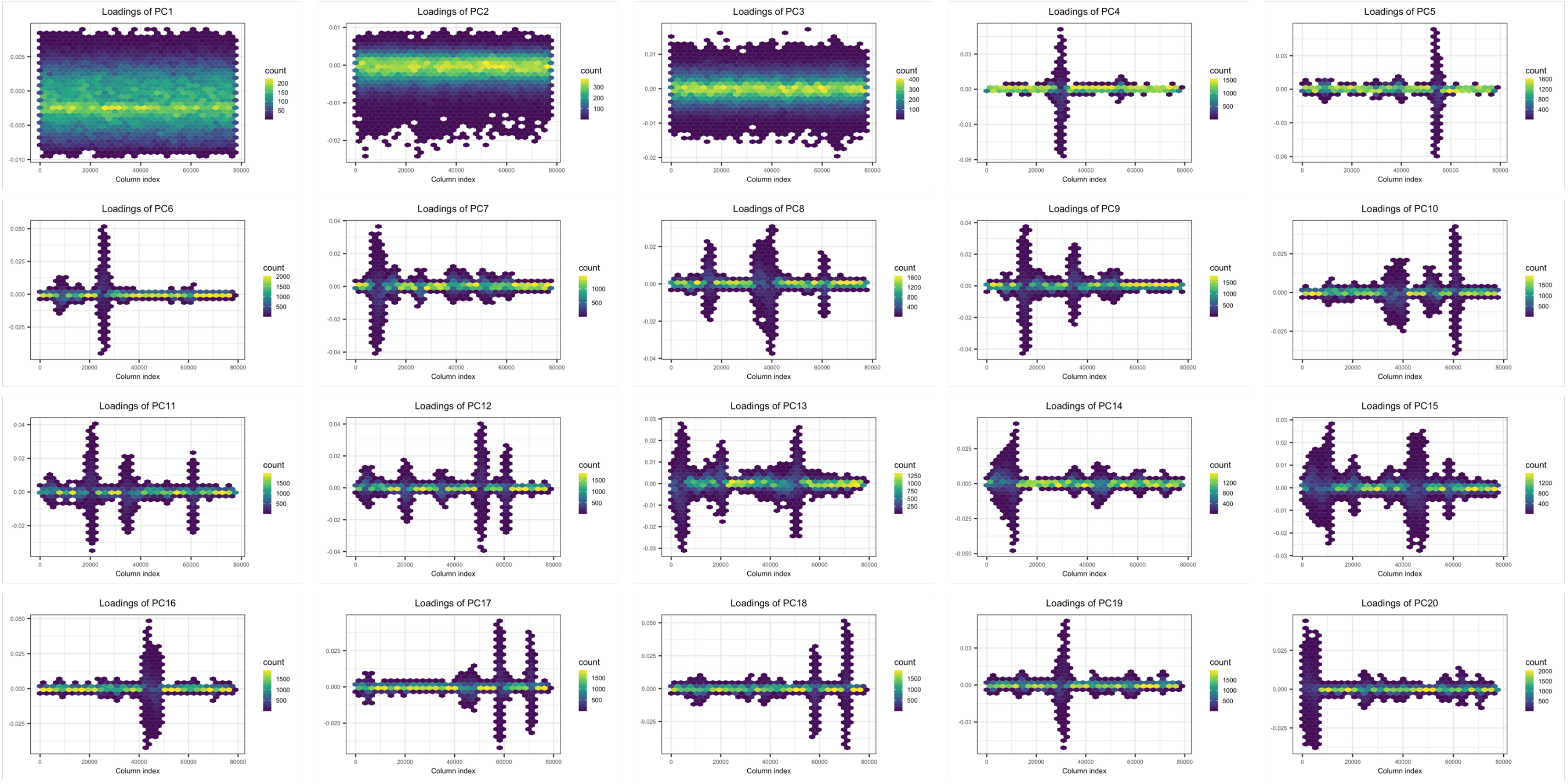


**Supplementary Figure 8**: PC SNP loadings from a PCA of 58,051 unrelated MCPS samples, using a LD $r^2$ threshold of 0.2 for SNP clumping.

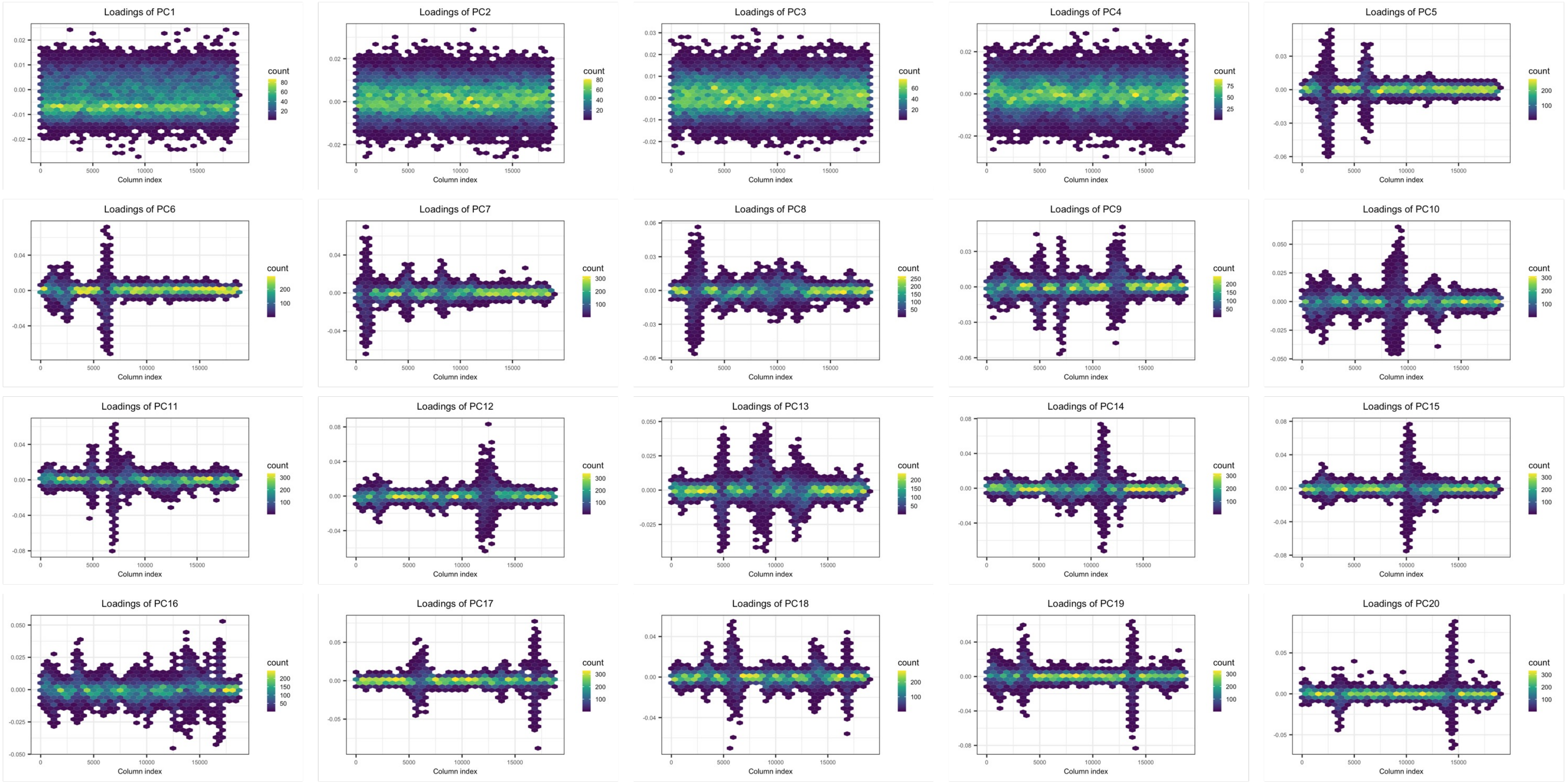


**Supplementary Figure 9**: PC SNP loadings from a PCA of 58,051 unrelated MCPS samples, using a LD $r^2$ threshold of 0.01 for SNP clumping.

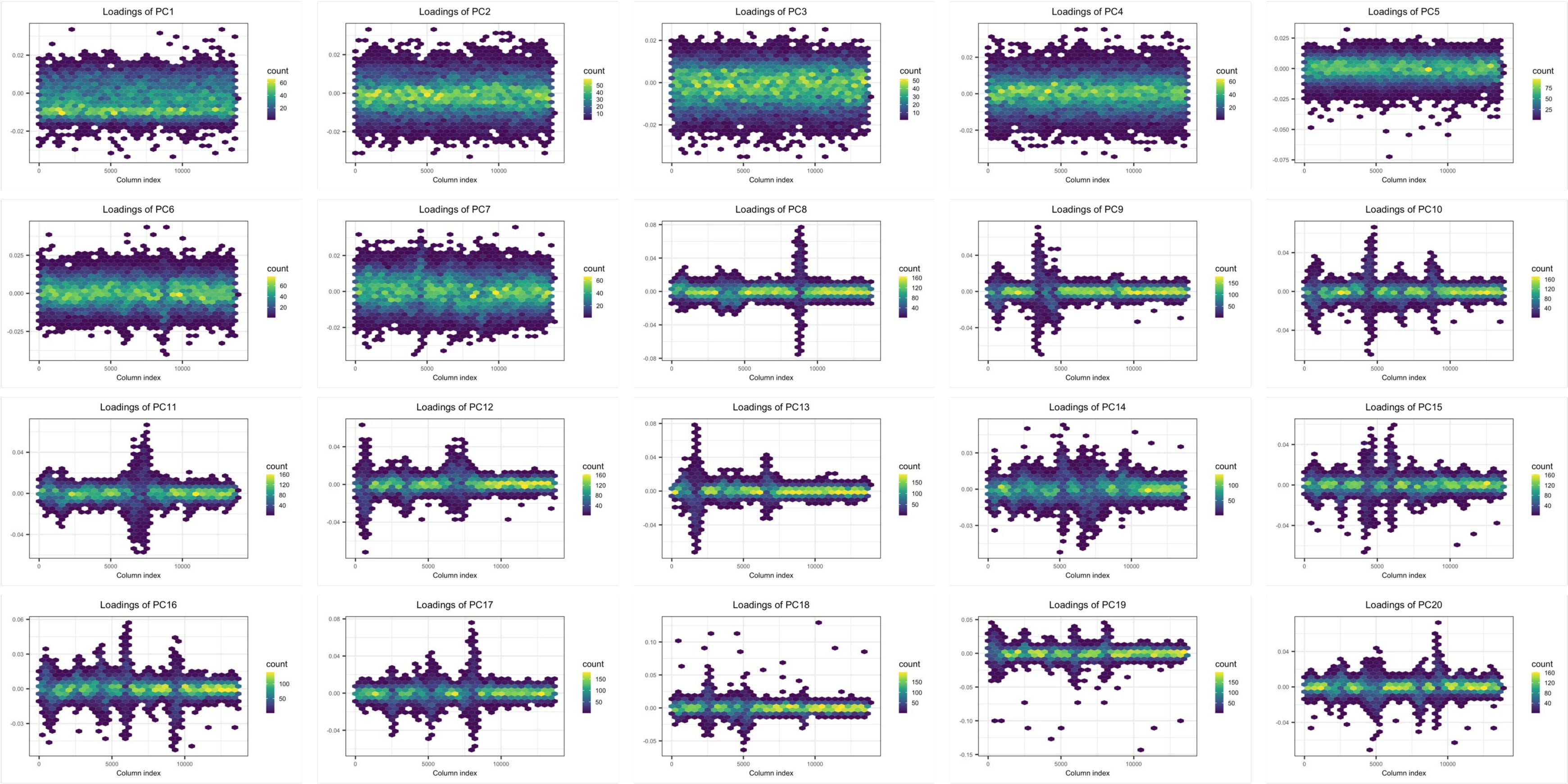


**Supplementary Figure 10**: PC SNP loadings from a PCA of 58,051 unrelated MCPS samples , using a LD $r^2$ threshold of 0.005 for SNP clumping.

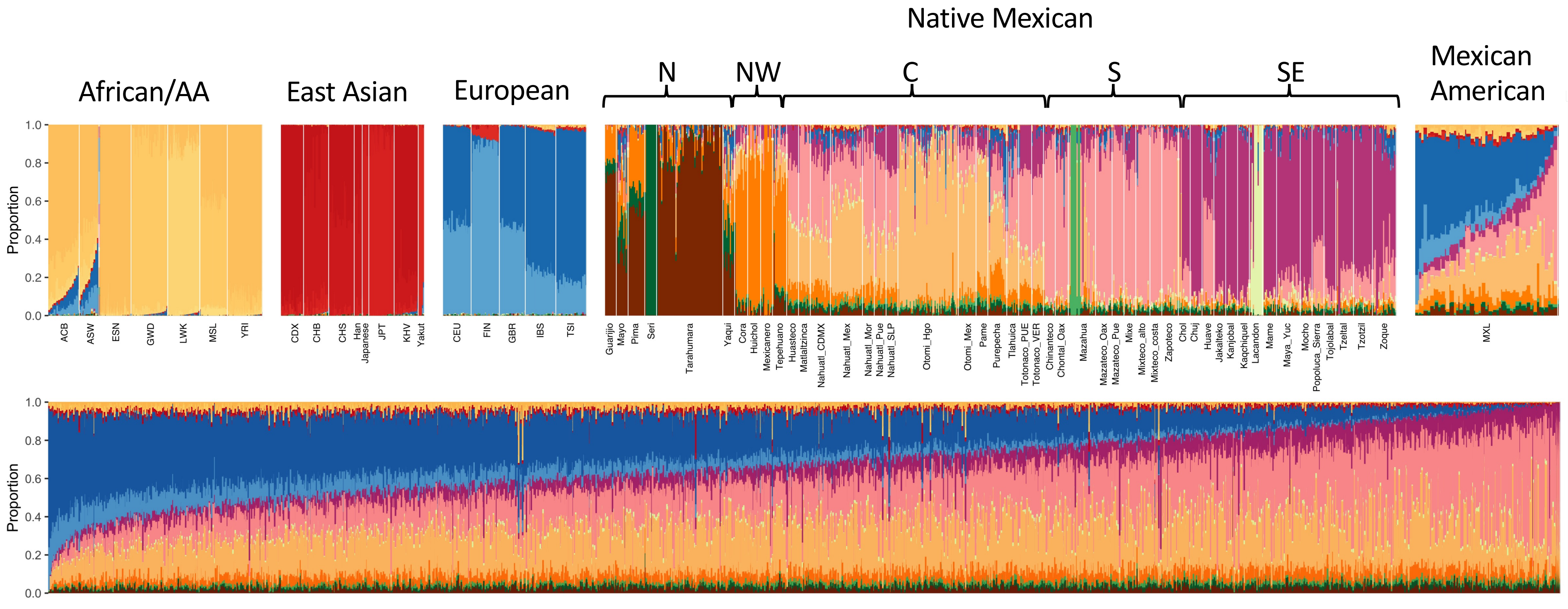


**Supplementary Figure 11: ADMIXTURE ancestry proportion estimates.** The program ADMIXTURE was used to estimate per-individual ancestry proportions and population-specific allele frequencies in a panel of 3,964 reference samples, including 1,000 MCPS samples. The remaining set of 137,511 MCPS samples were projected into the admixture model using parameter estimates from the reference sample. Results are shown for the K=18 model that attained the lowest cross-validation error. Ancestry proportion estimates for reference samples of African, European, and American ancestry from the 1KG, HGDP, and MAIS datasets are shown in the top row and estimates for MCPS participants are shown in the bottom row. AA=African American.

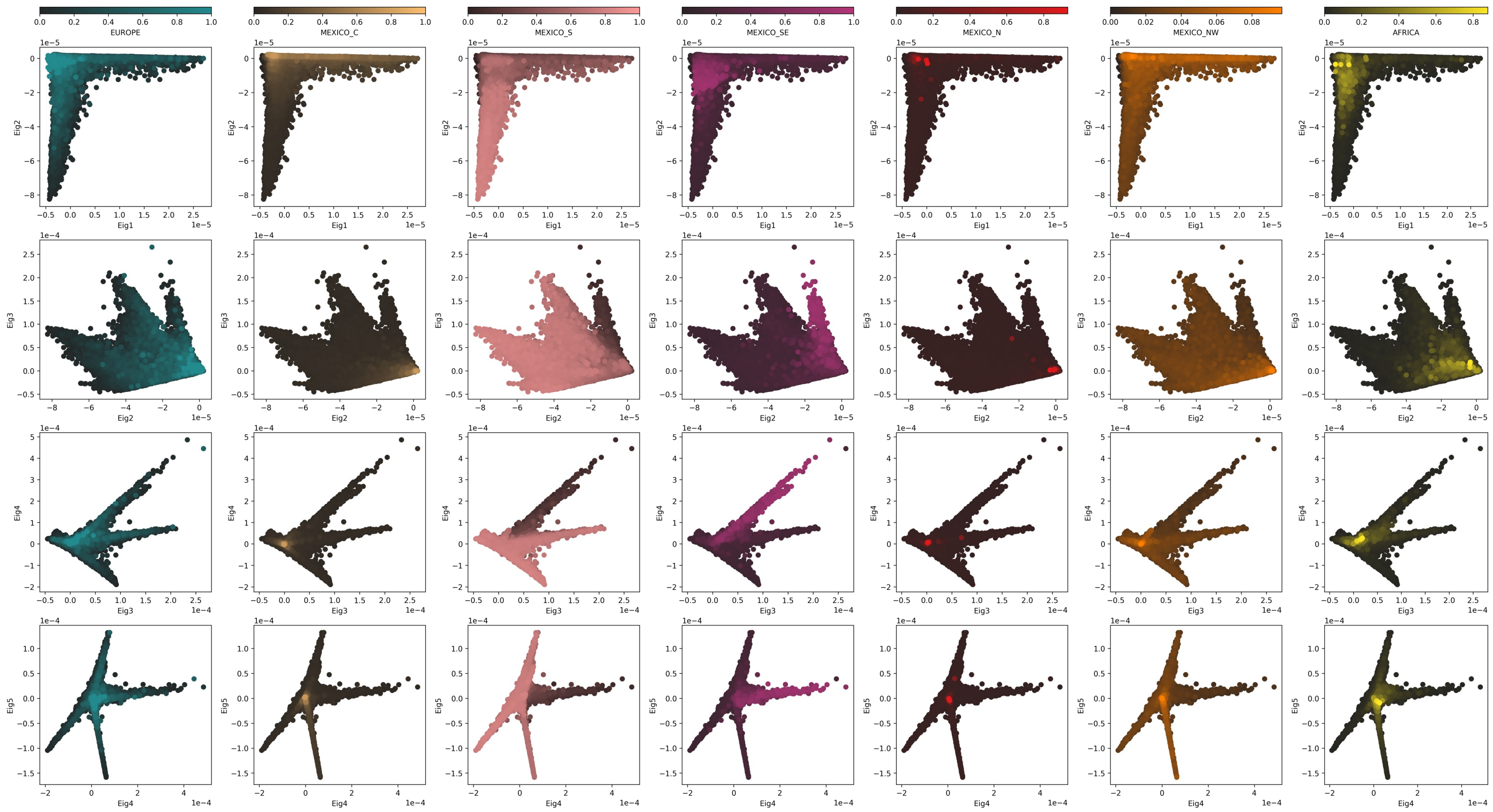


**Supplementary Figure 12 : Low-dimension visualization of IBD sharing.** Rows are different pairs of eigenvector scores. Columns show points coloured by seven different ancestry proportions obtained from the results of local ancestry inference with RFMix.

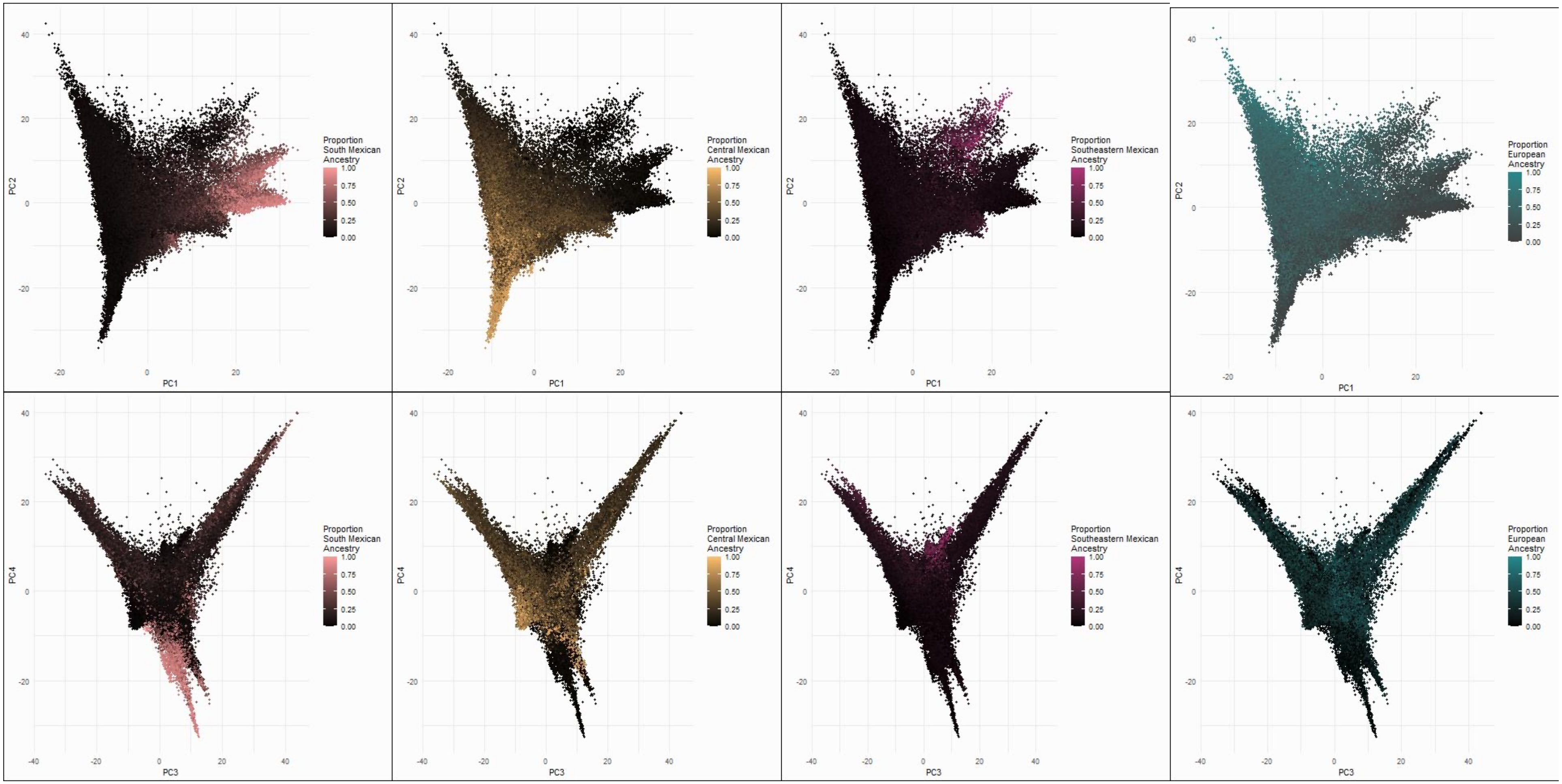


**Supplementary Figure 13 : Low-dimension visualization of haplotype sharing.** PC1 vs PC2 (Row 1) and PC3 vs PC4 (Row 2) . Columns are coloured using different ancestry proportions from the RFMix analysis.

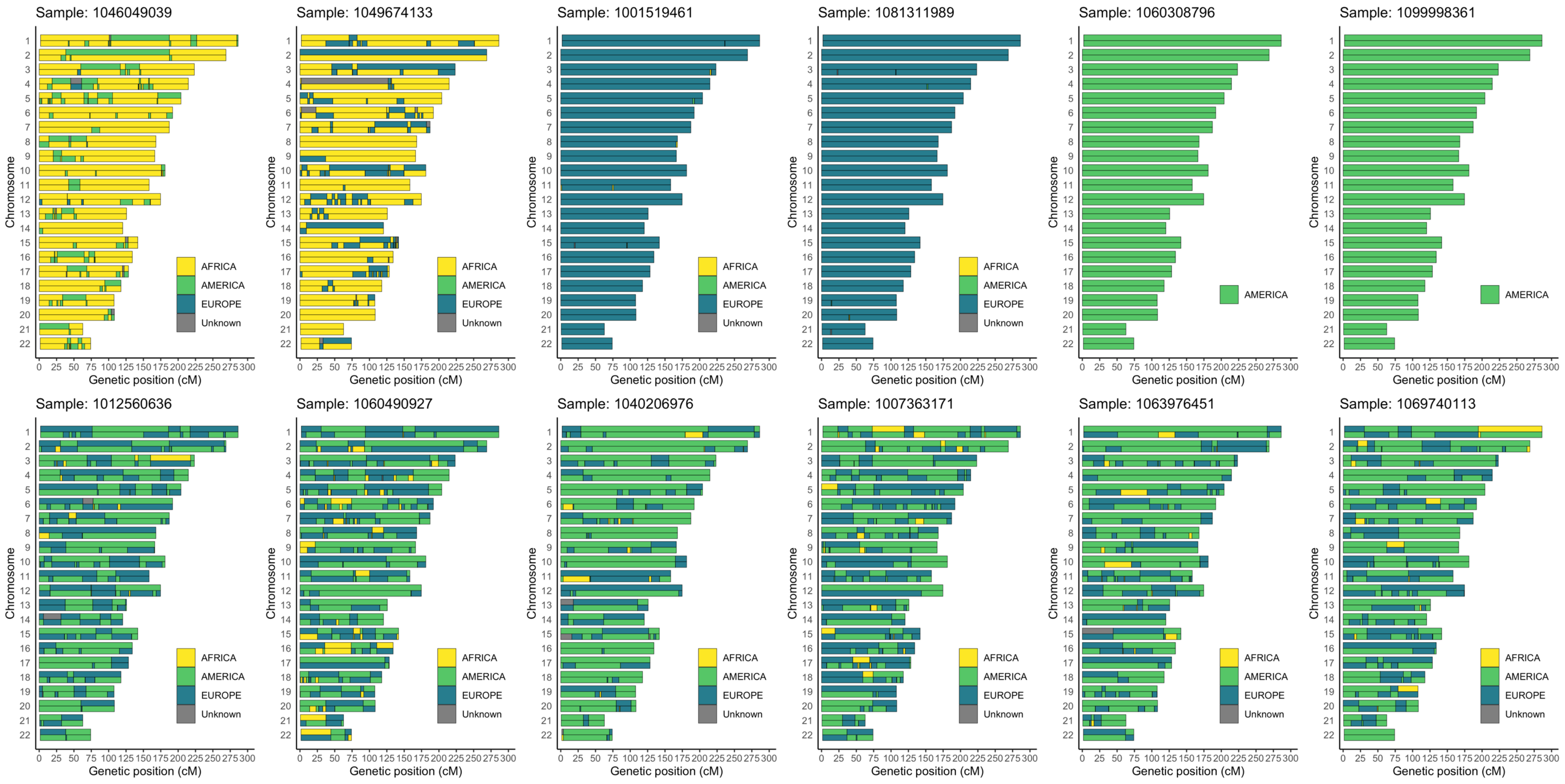


**Supplemental Figure 14: Karyograms showing genome segments from local ancestry inference.** LAI of a three-way admixture model was performed using a random-forest based method implemented in *RFMix*. Haplotypes with inferred genome segments derived from ancestral African (yellow), European (blue), and American (green) populations are shown for 12 MCPS samples. The top row shows samples with high proportions of African, European and Native Mexican ancestry. The bottom row shows more typical admixed samples.

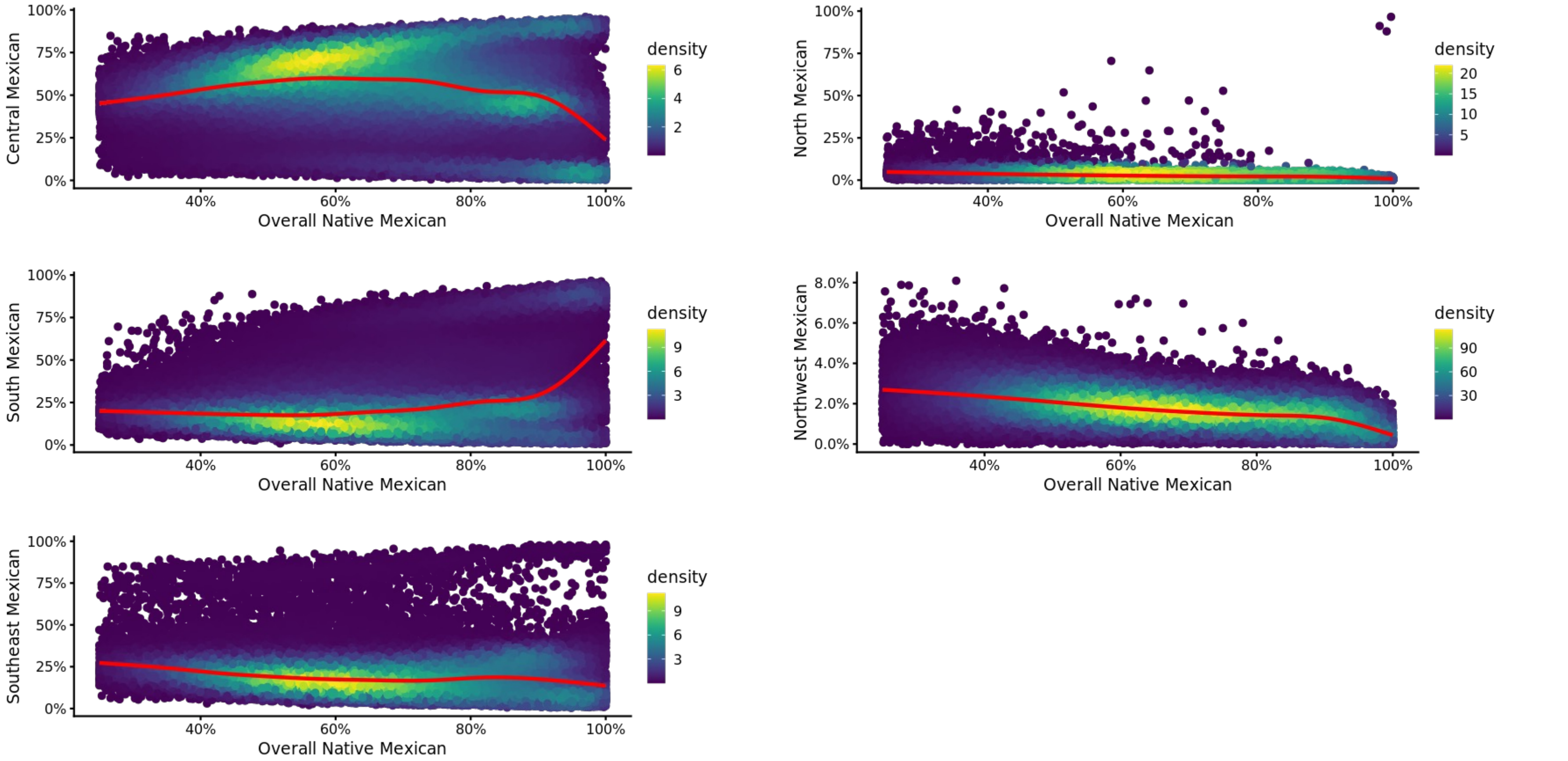


**Supplementary Figure 15 : Five Native Mexican ancestry components are compared to overall Native Mexican global ancestry proportions.** Each scatterplot shows individual-level ancestry proportions for 136,935 out of 138,511 MCPS samples with overall Native Mexican proportion > 25%. The point color underlines the estimated 2D density, and the red line is cubic-spline regression fit.

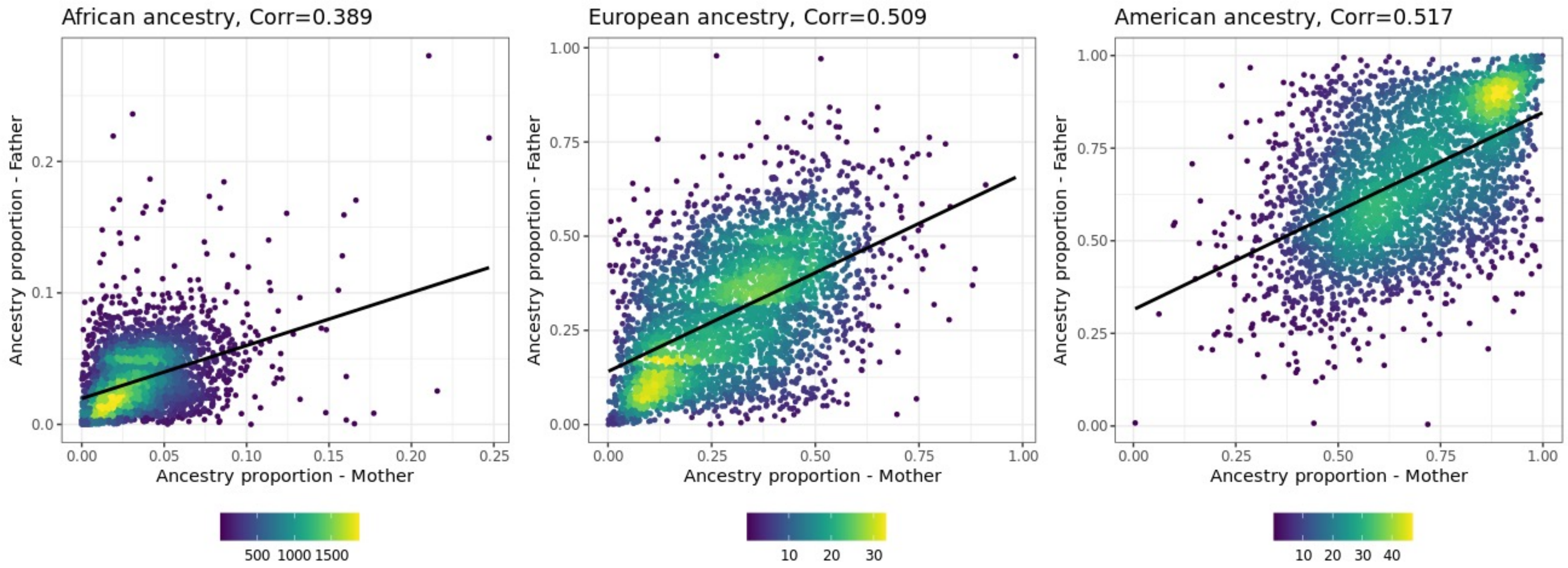


**Supplementary Figure 16 : Correlation in ancestry between spouses. The plots show the proportion of ancestry of the mother and father of** 3,595 parent couples inferred from the genetic relatedness analysis. Ancestry from the 3-way RFMix analysis was used to determine the proportions of African (left), European (middle) and American (right) ancestry.

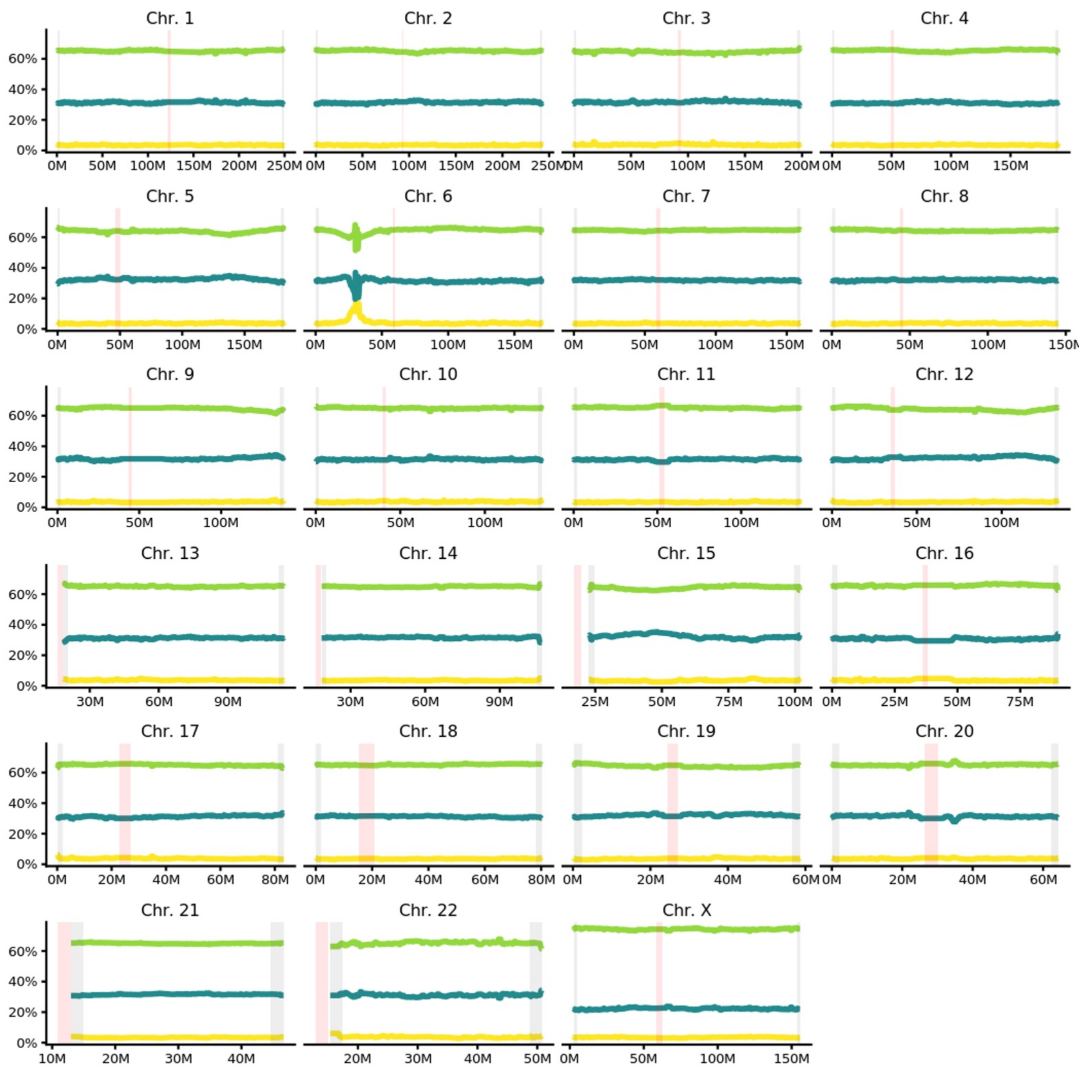


**Supplementary Figure 17 : Genome-wide distribution of local ancestry proportions.** The ancestry dosages inferred by RFMix are averaged across 78,833 unrelated MCPS samples and plotted along the genome. For each panel (or Chromosome) two gray rectangles denote terminal 2Mbp-length regions (of analyzed sites) at the beginning and end of Chromosome, while the red rectangle denotes the centromere region.

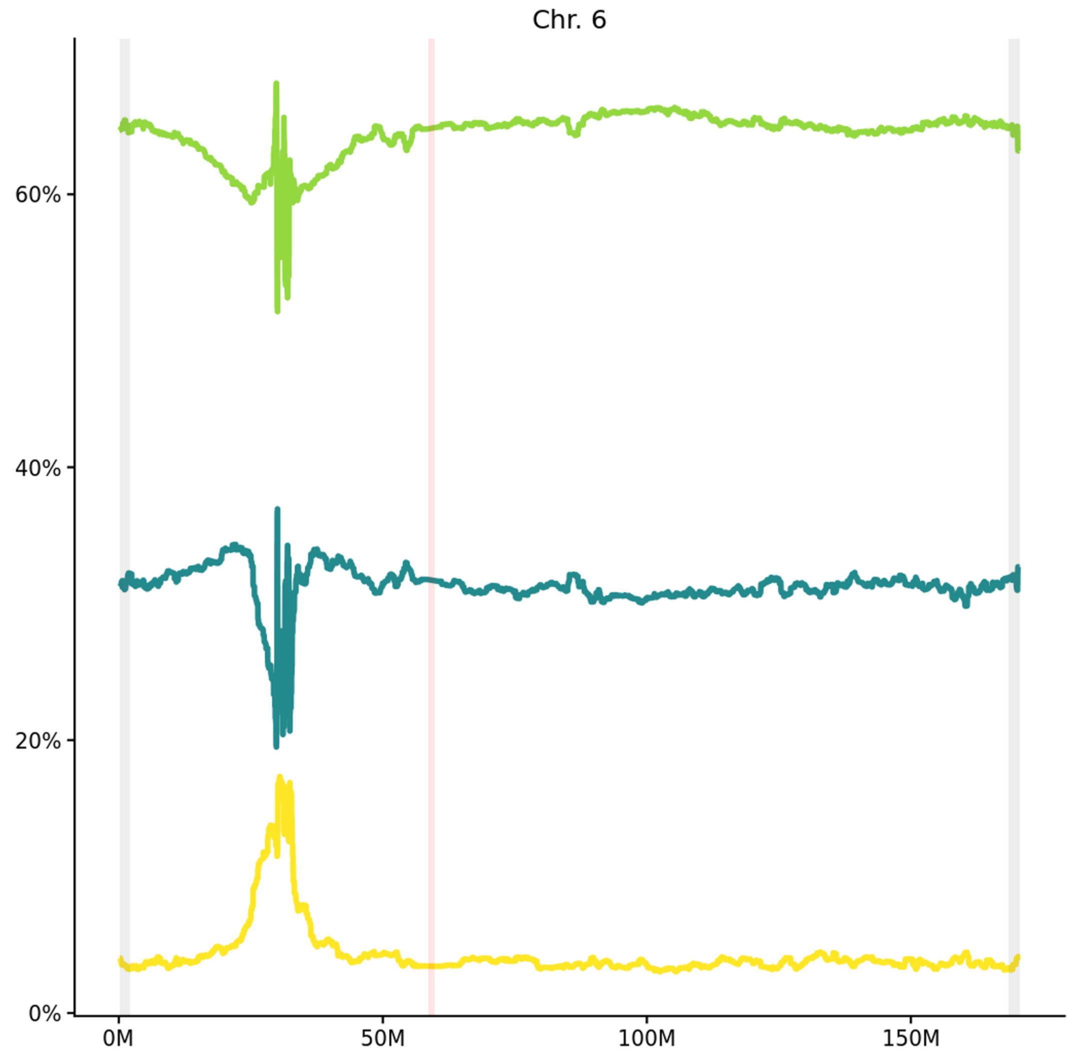


**Supplementary Figure 18 : Chromosome 6 distribution of local ancestry proportions.** The ancestry dosages inferred by RFMix are averaged across 78,833 unrelated MCPS samples and plotted along chromosome 6. The two gray bars denote terminal 2Mbp-length regions (of analyzed sites) at the beginning and end of the chromosome, while the red bar denotes the centromere region.

The peak of 17.3% African ancestry encompasses a region on chromosome 6 between 2.82 and 3.29 Mb. Under assumption of binomial sampling and normal approximation for sample mean, we obtain a p-value 2.9e-14 for African ancestry to exceed 17%. These results replicate findings previously reported in Guan et al 2014.

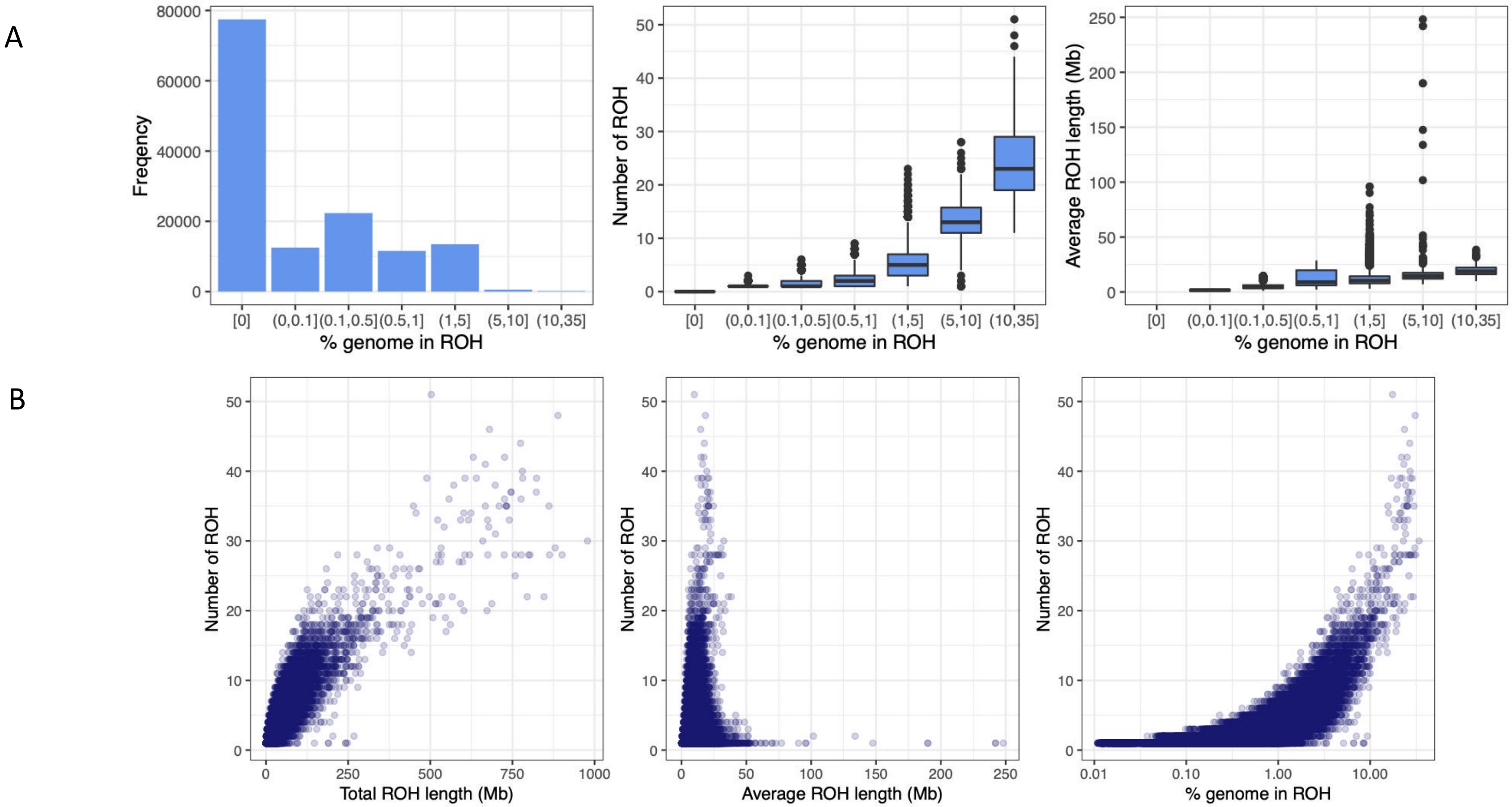


**Supplementary Figure 19 : Distribution of ROH (a)** Histogram of the sample counts, distribution of the per-sample number of ROH segments, and distribution of per-sample average ROH segment length are given by fraction of genome in ROH. **(b)** For each individual, the total length, average length, and fraction of genome in ROH is given by number of ROH.

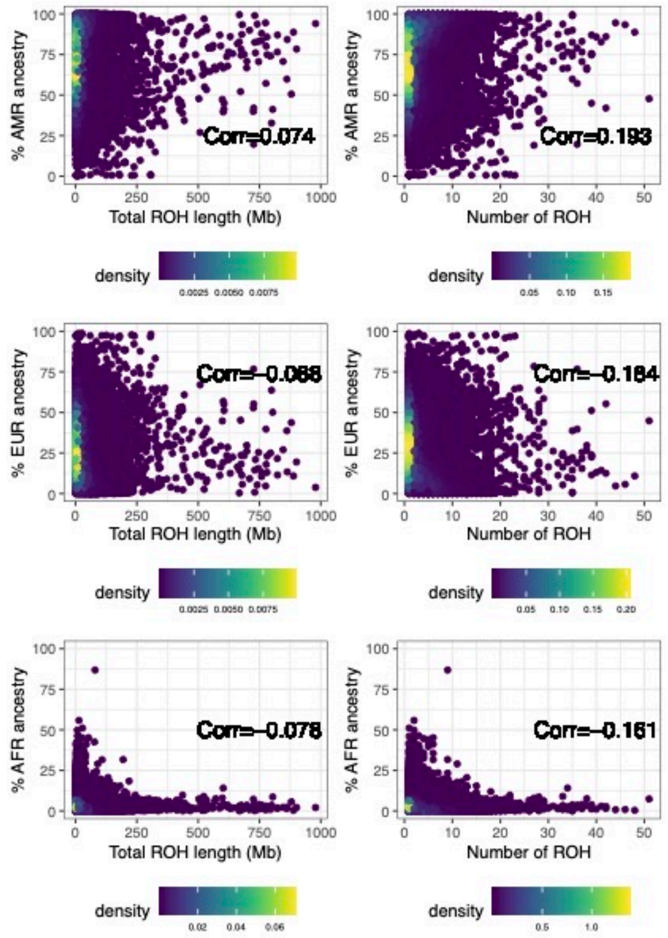


**Supplementary Figure 20 : ROH segments by ancestry** The fraction of ancestry attributed as Native Mexican, African, and European for each individual is given by total ROH length and number of ROH segments.

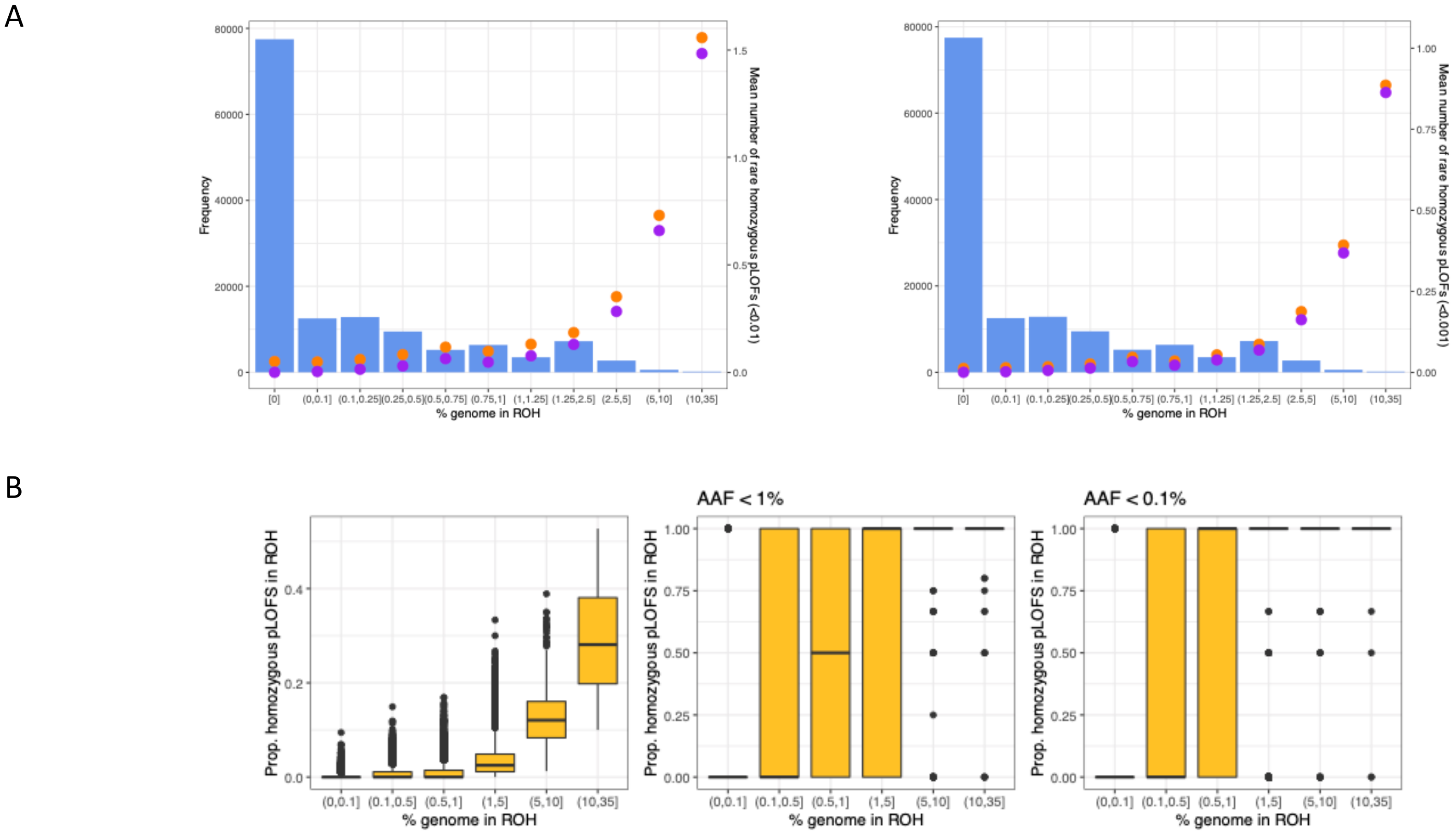


**Supplementary Figure 21 : Loss of function variants by ROH (A)** The number of individuals and average number of rhLOFs within ROH (purple) and overall (orange) are given by the fraction of the genome in ROH. **(B)** The per-sample proportion of homozygous pLOFs falling within ROH are given by fraction of the genome in ROH, for all frequencies, AAF<1%, and AAF<0.1%.

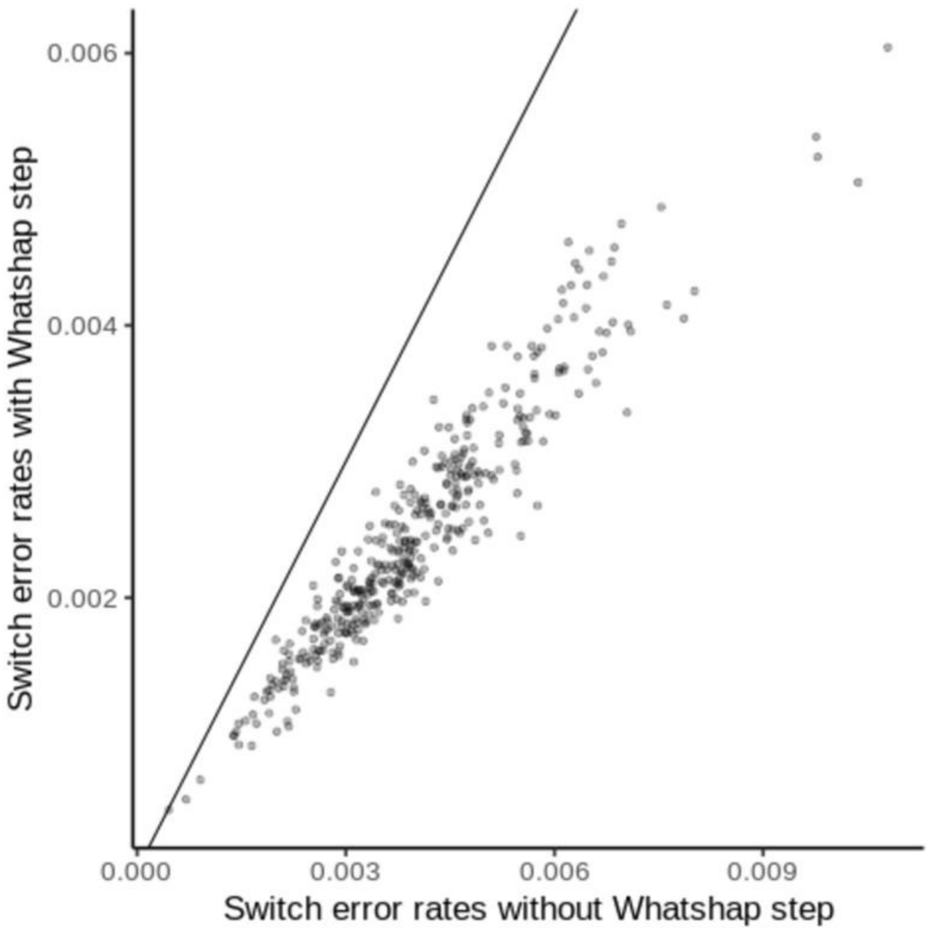


**Supplementary Figure 22 : Phasing accuracy of WGS dataset**. The phasing switch error for each of 392 individuals phased with (y-axis) and without (x-axis) using the the WhatsHap method to leverage phase information in sequencing reads. The 392 individuals are parents of mother-father-child trios that were phased without including the children.

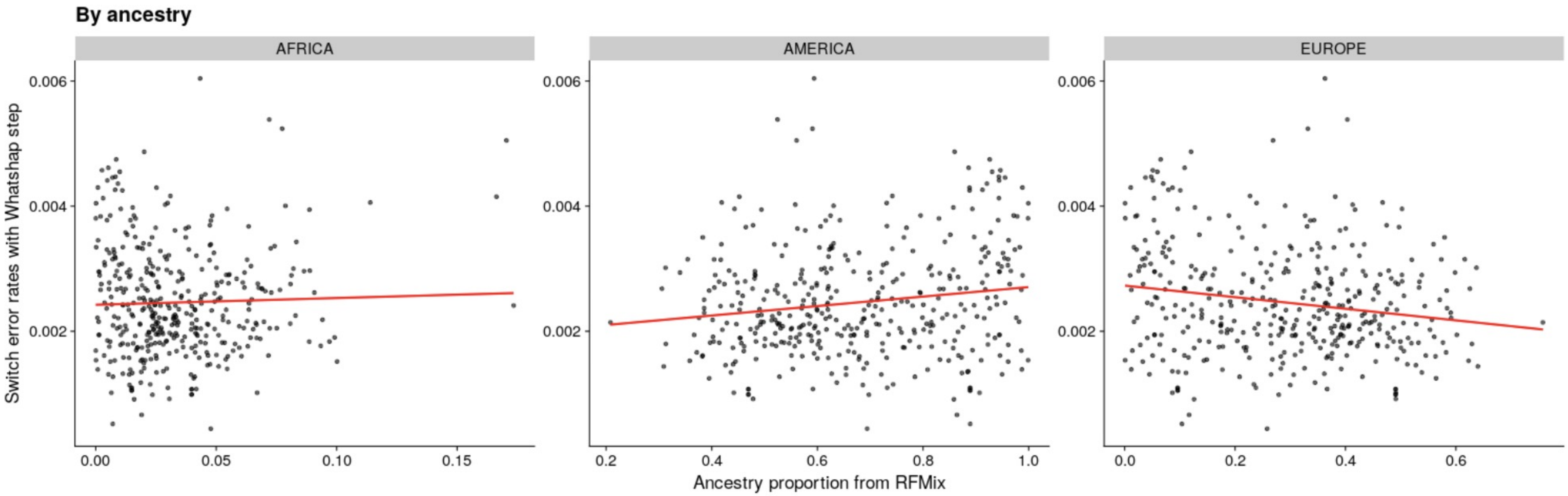


**Supplementary Figure 23 : Phasing accuracy of WGS dataset stratifies by ancestry**. The phasing switch error (y-axis) for each of 392 individuals plotted against estimated proportion (x-axis) of African ancestry (left), American ancestry (middle) and European ancestry (right). The p-values for the test that the regression line being different from 0 was 0.0016, 0.0004 and 0.53 for proportion of American, European and African ancestry respectively. The 392 individuals are parents of mother-father-child trios that were phased without including the children.

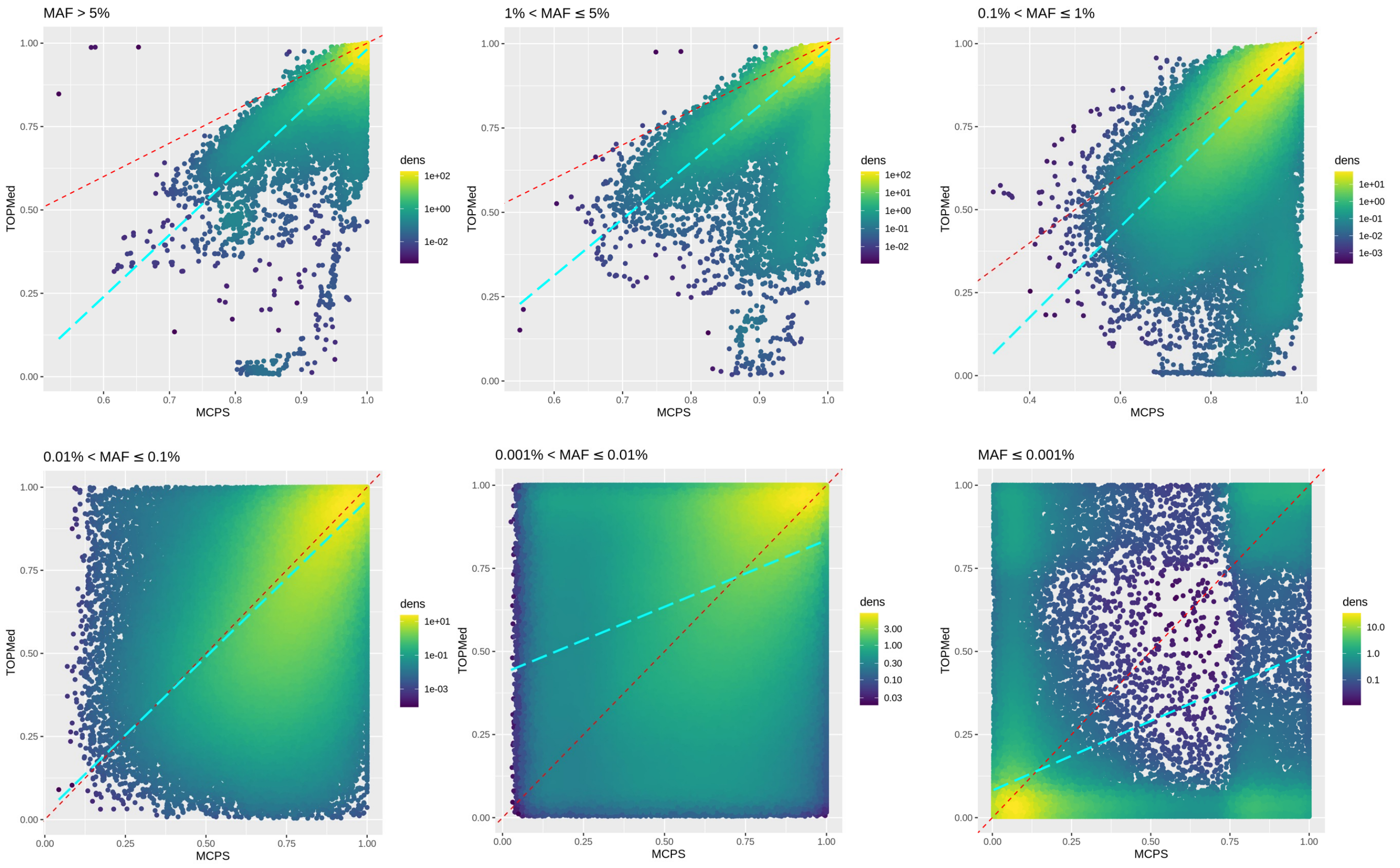


**Supplementary Figure 24 : Comparison of MCPS10k and TOPMed imputation.** Plots show imputation info scores from MCS10k and TOPMed imputed variants in 67,079 MCPS samples at 6,473,872 variants on chromosome 2. Each plot uses a different MAF bin. The red line is Y=X. The blue dashed line is the regression line.

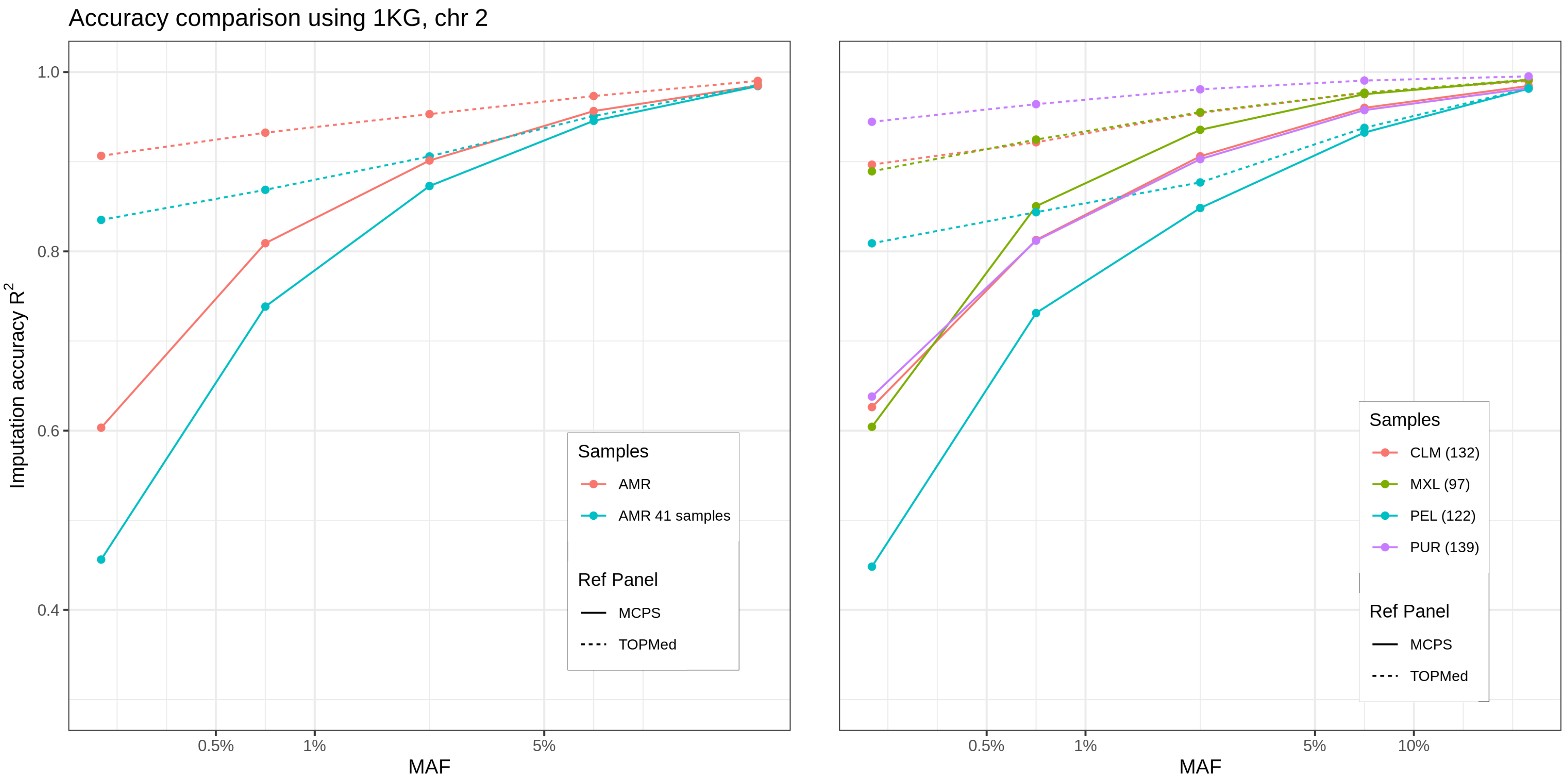


**Supplementary Figure 25 : Imputation accuracy using the MCPS10k and TOPMed imputation panels applied to 1000 Genomes samples.** Accuracy is measured using the $R^2$ between the imputed variants and 1,610,370 variants from the 1000 Genomes WGS dataset. Imputation was based on genotypes at SNPs on the Illumina HumanOmniExpressExome-8v1-2_A array. Results are stratified by allele frequency in the 490(x-axis on log10 scale), reference panel (solid = MCPS, dotted = TOPMed) and by populations. In the left hand plot shows results at on all samples with American ancestry (AMR) and on the 41 samples with >90% American ancestry. The right hand plot shows the results stratified by the four groups : Mexican ancestry from Los Angeles (MXL), Peruvian ancestry from Lima (PEL), Colombian ancestry from Medellin (CLM) and Puerto Rican ancestry from Puerto Rico (PUR) .

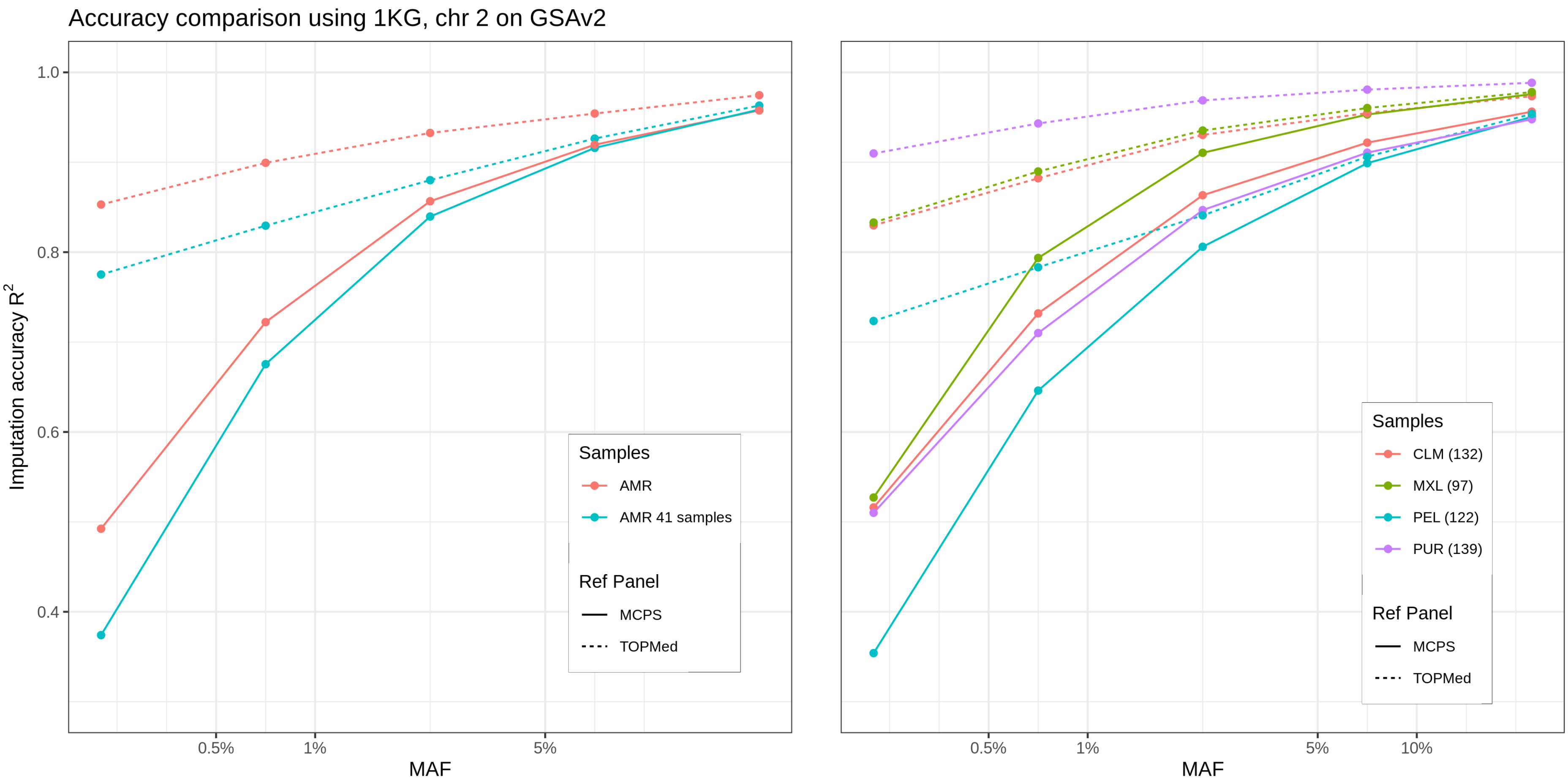


**Supplementary Figure 26 : Imputation accuracy using the MCPS10k and TOPMed imputation panels applied to 1000 Genomes samples.** Accuracy is measured using the $R^2$ between the imputed variants and 1,610,370 variants from the 1000 Genomes WGS dataset. Imputation was based on genotypes at SNPs on the Illumina Global Screening Array (GSA v2). Results are stratified by allele frequency in the 490(x-axis on log10 scale), reference panel (solid = MCPS, dotted = TOPMed) and by populations. In the left hand plot shows results at on all samples with American ancestry (AMR) and on the 41 samples with >90% American ancestry. The right hand plot shows the results stratified by the four groups : Mexican ancestry from Los Angeles (MXL), Peruvian ancestry from Lima (PEL), Colombian ancestry from Medellin (CLM) and Puerto Rican ancestry from Puerto Rico (PUR) .

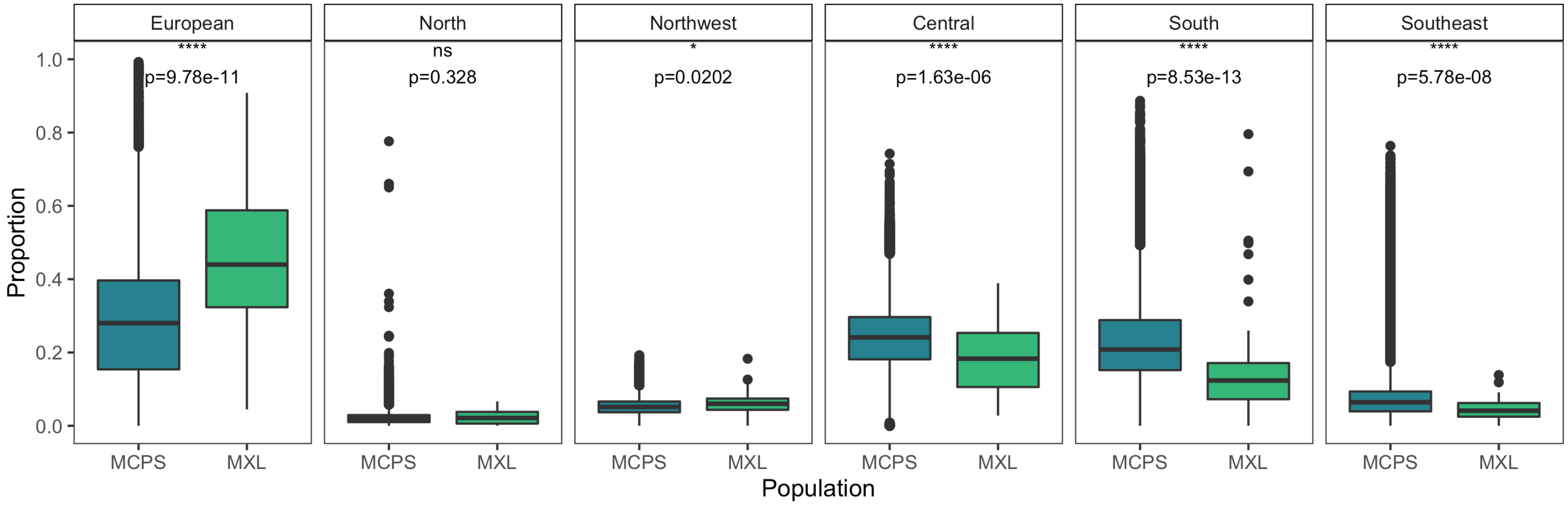


**Supplementary Figure 27 : Comparison of ancestry between MCPS and MXL samples**. We identified the groups from the ADMIXTURE analysis (Supplementary Figure 11) that had the highest average proportions in each of the five Mexico indigenous regions (i.e. North, Northwest, etc) and Europe and compared these ancestry proportions between the 1000 Genomes MXL and MCPS cohorts with Mann Whitney U tests.

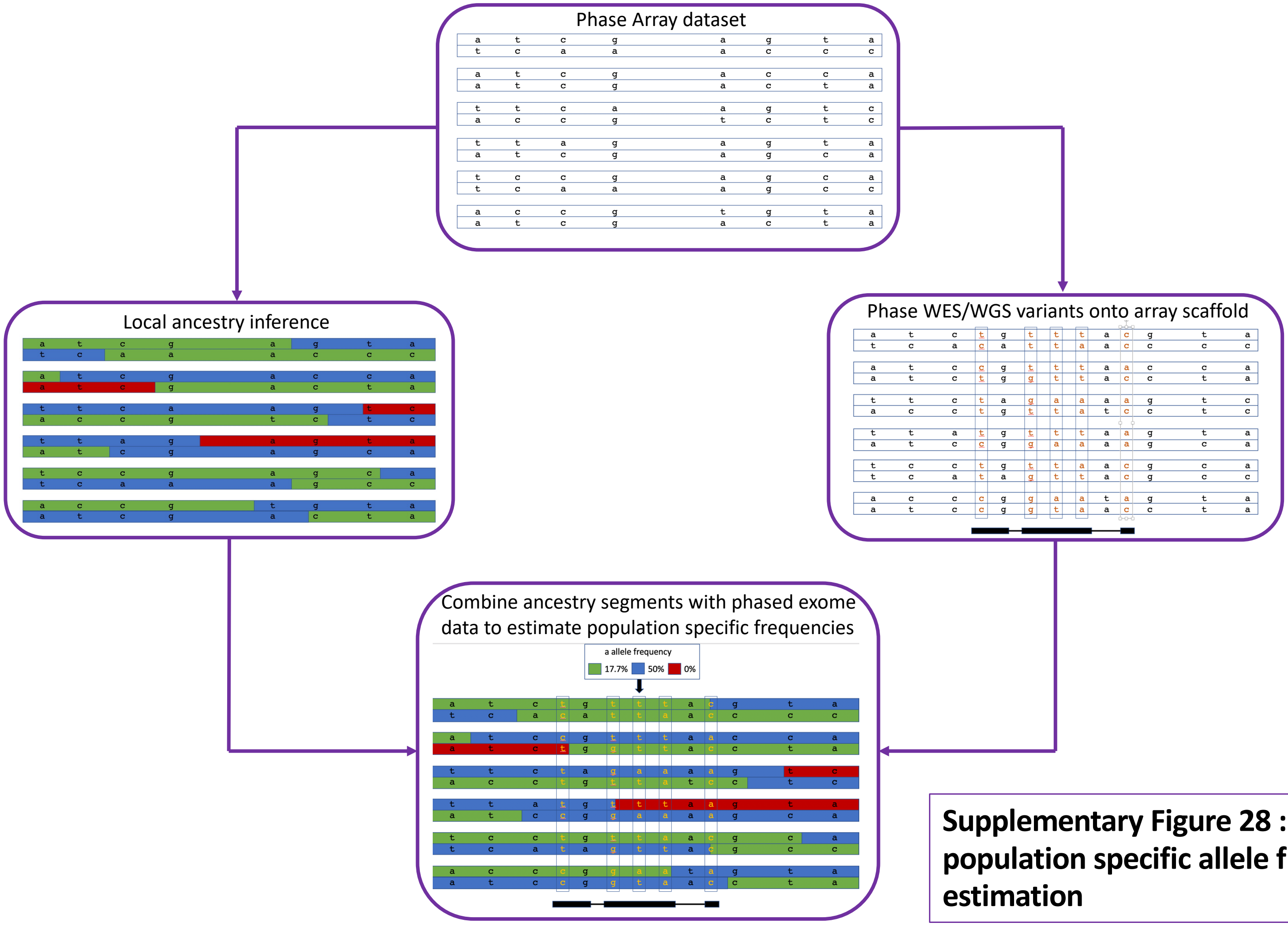


**Supplementary Figure 28 : Schematic of population specific allele frequency estimation**

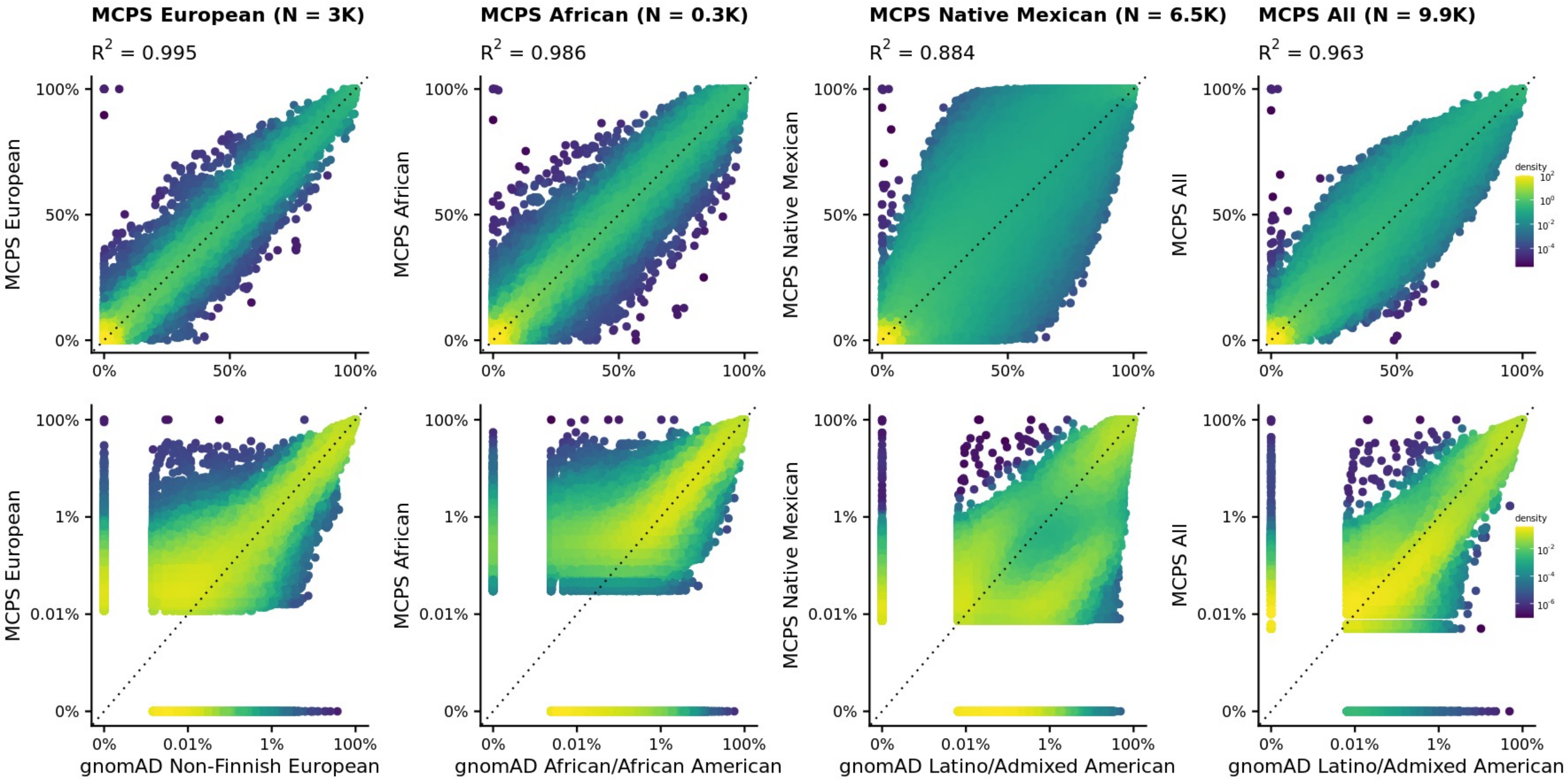


**Supplementary Figure 29 : Allele frequency comparison between MCPS WGS and gnomAD.** Allele frequencies on linear (top) and log (bottom) scale. The comparisons from left to right are MCPS European vs gnomAD Non-Finnish European, MCPS African vs gnomAD African, MCPS Native American vs gnomAD Latino/Admixed American amd Overall MCPS vs gnomAD Latino/Admixed American. The number N in each title represents the estimated effective sample size for the MCPS allele frequency estimation.

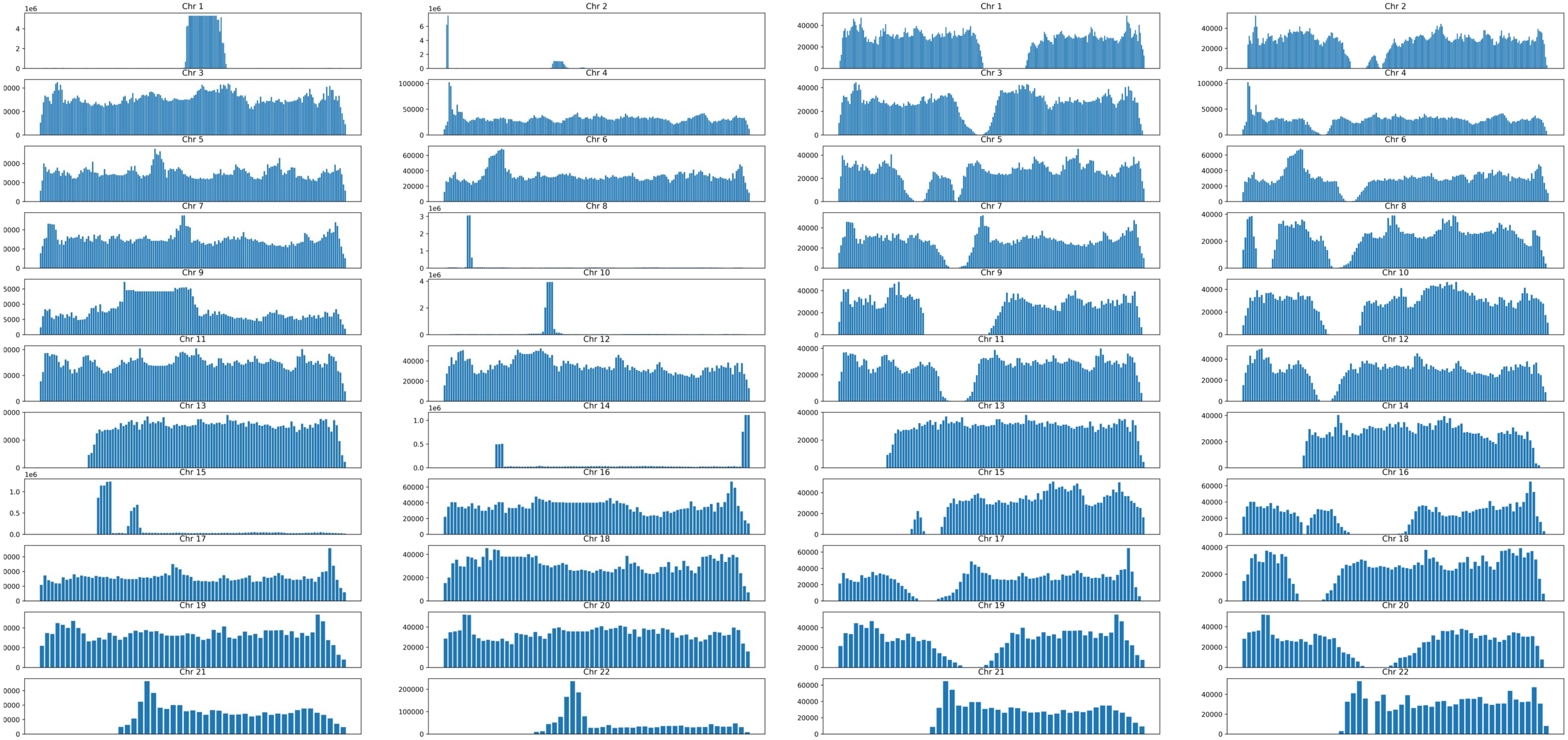


**Supplementary Figure 30 : IBD segment coverage from 10K MCPS samples before (left) and after (right) filtering.** IBD segments were filtered out if they intersected a 1Mb bin with either i) fourfold more than the median IBD coverage along a chromosome, or ii) fourfold fewer than the median number of SNP array markers.
