## Supplementary Tables for "Genotyping, sequencing and analysis of 140,000 adults from the Mexico City Prospective Study"

**Supplementary Table 1** : Cohort characteristics. Measures are reported for all 141,046 samples that were exome sequenced.

| Characteristic | Men (n=46 069) | Women (n=94 977) | All (n=141 046) |
|---|---|---|---|
| Age, years | 54 (13) | 53 (13) | 53 (13) |
| **Socioeconomic and lifestyle characteristics** | | | |
| Resident of Coyoacán | 18 805 (41%) | 34 684 (37%) | 53 489 (38%) |
| University/college educated | 10 561 (23%) | 10 517 (11%) | 21 078 (15%) |
| Current smoker | 22 326 (48%) | 21 292 (22%) | 43 618 (31%) |
| Current drinker | 34 566 (75%) | 57 934 (61%) | 92 500 (66%) |
| Any regular leisure-time physical activity | 13 565 (29%) | 17 495 (18%) | 31 060 (22%) |
| **Physical measurements** | | | |
| Height, cm | 164 (7) | 151 (7) | 156 (9) |
| Weight, kg | 76 (13) | 68 (13) | 70 (13) |
| BMI, kg/m² | 27.9 (4.4) | 29.6 (5.3) | 29.0 (5.1) |
| Waist circumference, cm | 96 (11) | 94 (12) | 94 (12) |
| Hip circumference, cm | 101 (8) | 106 (11) | 105 (11) |
| Waist-hip ratio | 0.95 (0.07) | 0.88 (0.07) | 0.90 (0.08) |
| SBP, mmHg | 129 (16) | 127 (17) | 128 (17) |
| DBP, mmHg | 84 (10) | 83 (10) | 83 (10) |
| Glycated hemoglobin, % | 5.5 (5.3-6.0) | 5.5 (5.3-6.0) | 5.5 (5.3-6.0) |
| **Self-reported medication** | | | |
| Anti-diabetic | 5045 (11%) | 10 697 (11%) | 15 742 (11%) |
| Anti-hypertensive | 5600 (12%) | 17 255 (18%) | 22 855 (16%) |
| Anti-thrombotic | 1404 (3%) | 2932 (3%) | 4336 (3%) |
| Lipid lowering | 292 (1%) | 485 (1%) | 777 (1%) |
| **Self-reported prior diseases** | | | |
| Diabetes* | 6476 (14%) | 13 219 (14%) | 19 695 (14%) |
| Coronary heart disease | 972 (2%) | 1204(1%) | 2176 (2%) |
| Stroke | 577 (1%) | 1061 (1%) | 1638 (1%) |
| Cancer | 317 (1%) | 1415 (1%) | 1732 (1%) |
| Liver cirrhosis | 133 (<0.5%) | 77 (<0.5%) | 210 (<0.5%) |
| Emphysema | 224 (<0.5%) | 206 (<0.5%) | 430 (<0.5%) |
| Chronic kidney disease | 354 (1%) | 825 (1%) | 1179 (1%) |

Numbers shown are mean (SD), median (IQR) or n (%), and are calculated on all available data (missing data percentages are low: <0.1% for each of the socio-economic and lifestyle characteristics, between 0.1% and 1.2% for the physical measurements and 0.3% for glycated hemoglobin). BMI=Body mass index, DBP=Diastolic blood pressure, SBP=Systolic blood pressure. * Doctor diagnosed and/or use of an antidiabetic medication.

* Doctor diagnosed and/or use of an antidiabetic medication.

**Supplementary Table 2 : Number of canonical coding variants discovered in exome sequencing of 141,146 MCPS participants**. Variants were annotated using VEP. Predicted function for each variant was defined by consequence in canonical coding transcripts in Ensembl v100. MAC = Minor Allele Count, IQR = Inter Quartile Range, SD = Standard Deviation

| Variant category (Canonical transcript) | N variants (% with MAC=1) | Median number of alternate alleles per participant (IQR) | Mean number of alternate alleles per participant (SD) | Median number of variants per participant (IQR) | Mean number of variants per participant (SD) |
|---|---|---|---|---|---|
| Coding regions | 3883715 (30.93) | 27646 (281) | 27652 (226) | 19800 (592) | 19751 (429) |
| **Predicted function** | | | | | |
| In-frame indels | 42878 (31.03) | 266 (15) | 266 (12) | 196 (12) | 196 (10) |
| Synonymous | 1211674 (28.03) | 14708 (166) | 14711 (132) | 10504 (320) | 10479 (233) |
| Missense | 2418983 (31.51) | 12474 (155) | 12476 (120) | 8938 (273) | 8919 (197) |
| Likely benign | 488296 (27.9) | 9103 (115) | 9104 (89) | 6194 (170) | 6183 (124) |
| Possibly deleterious | 1381266 (31.24) | 3256 (71) | 3257 (53) | 2628 (105) | 2624 (77) |
| Likely deleterious | 549421 (35.4) | 114 (16) | 114 (12) | 111 (16) | 111 (12) |
| pLOF | 210180 (40.89) | 200 (15) | 200 (11) | 157 (14) | 157 (10) |
| Start lost | 6247 (36.79) | 8 (3) | 9 (2) | 7 (2) | 7 (2) |
| Stop gain | 71453 (39.66) | 62 (9) | 62 (6) | 48 (7) | 49 (5) |
| Stop lost | 2229 (37.64) | 4 (2) | 4 (2) | 3 (1) | 3 (1) |
| Splice donor | 23218 (41.05) | 15 (5) | 15 (3) | 13 (4) | 13 (3) |
| Frameshift | 89376 (41.98) | 91 (11) | 92 (8) | 72 (10) | 72 (7) |
| Splice acceptor | 17657 (42.04) | 18 (4) | 18 (3) | 14 (3) | 14 (2) |

**Supplementary Table 3.** Number of coding variants available in MCPS WES, UKB WES, gnomAD, and TOPMed. MCPS variants were annotated with VEP, with function defined as the most deleterious consequence based on overlap with any protein-coding transcript in Ensembl v100. Variants for UKB WES, TOPMED Freeze 8, and gnomAD v3.1.2 were annotated with SNPeff, with function defined as the most deleterious consequence based on overlap with any protein-coding transcript in Ensembl v85. LOF= loss of function.

| Variant Type | MAF | MCPS Freeze 150 WES All ancestries | | UKB WES All ancestries (N=454,787) | TOPMed Freeze 8[a] All ancestries (N=132,345) | gnomAD 3.1[a] All ancestries (N=76,156) |
|---|---|---|---|---|---|---|
| | | Total Variants | Unique to MCPS | | | |
| Synonymous | All | 1,233,054 | 361,248 | 3,457,173 | 2,396,982 | 2,062,223 |
| | Singleton | 345,730 | 163,115 | 1,490,793 | 996,855 | 981,579 |
| | Doubleton - 0.01% | 689,869 | 192,613 | 1,813,474 | 1,165,582 | 838,004 |
| | 0.01-0.1% | 137,122 | 5,485 | 110,632 | 143,488 | 146,861 |
| | 0.1-1% | 34,803 | 18 | 20,256 | 50,742 | 53,974 |
| | 1-5% | 8,027 | 3 | 7,583 | 19,111 | 19,495 |
| | >5% | 17,503 | 14 | 14,435 | 21,204 | 22,310 |
| Missense | All | 2,526,776 | 892,337 | 7,878,586 | 5,063,772 | 4,231,655 |
| | Singleton | 793,479 | 413,577 | 3,724,820 | 2,343,133 | 2,237,366 |
| | Doubleton - 0.01% | 1,435,164 | 466,750 | 3,919,257 | 2,388,372 | 1,646,461 |
| | 0.01-0.1% | 228,563 | 11,958 | 179,390 | 228,510 | 234,619 |
| | 0.1-1% | 44,798 | 25 | 32,783 | 65,209 | 70,941 |
| | 1-5% | 8,742 | 5 | 9,418 | 19,718 | 21,104 |
| | >5% | 16,030 | 22 | 12,918 | 18,830 | 21,164 |
| LOF | All | 233,650 | 125,344 | 915,289 | 507,022 | 426,399 |
| | Singleton | 93,590 | 64,553 | 529,763 | 289,368 | 268,983 |
| | Doubleton - 0.01% | 124,147 | 59,578 | 372,343 | 201,473 | 134,760 |
| | 0.01-0.1% | 13,152 | 1,208 | 10,694 | 12,067 | 15,843 |
| | 0.1-1% | 1,921 | 3 | 1,821 | 2,795 | 4,554 |
| | 1-5% | 314 | 0 | 317 | 730 | 1,192 |
| | >5% | 526 | 2 | 351 | 589 | 1,067 |
| All coding[b] | All | 3,993,480 | 1,378,929 | 12,251,048 | 7,967,776 | 6,720,277 |
| | Singleton | 1,232,799 | 641,245 | 5,745,376 | 3,629,356 | 3,487,928 |
| | Doubleton - 0.01% | 2,249,180 | 718,941 | 6,105,074 | 3,755,427 | 2,619,225 |
| | 0.01-0.1% | 378,837 | 18,651 | 300,716 | 384,065 | 397,323 |
| | 0.1-1% | 81,522 | 46 | 54,860 | 118,746 | 129,469 |
| | 1-5% | 17,083 | 8 | 17,318 | 39,559 | 41,791 |
| | >5% | 34,059 | 38 | 27,704 | 40,623 | 44,541 |

a Pass variants. No additional HWE or genotype missingess filters applied to gnomAD or TOPmed.
No restriction to same ""coding regions"" across datasets."
b Includes synonymous, missense and LOF variants only.

**Supplementary Table 4.** Number of autosomal genes with at least N carriers of rare LOFs (AAF <= 1%) in MCPS WES and UK Biobank WES**.** Variants in MCPS were annotated with VEP and Ensembl v100. Variants in UKB were annotated with SNPeff and Ensembl v85. LOFs were defined as stop gain, stop loss, start lost, splice donor, splice acceptor, or frameshift variants impacting any protein-coding transcript.

| Zygosity | Source | Number of carriers | | | | | | | |
|---|---|---|---|---|---|---|---|---|---|
| | | 1+ | 5+ | 10+ | 25+ | 50+ | 100+ | 500+ | 1000+ |
| Heterozygous | Observed in MCPS WES 150K[a] | 15,912 | 15,900 | 14,196 | 11,059 | 8,091 | 5,296 | 1,168 | 492 |
| | Observed in UKB WES 150K | 18,065 | 17,435 | 16,415 | 13,576 | 10,300 | 6,587 | 1,505 | 721 |
| | Observed in UKB WES 450K | 18,131 | 18,028 | 17,787 | 16,915 | 15,289 | 12,684 | 4,629 | 2,423 |
| Homozygous | Observed in MCPS WES 150K[a] | 2,482 | 380 | 145 | 12 | 0 | 0 | 0 | 0 |
| | Observed in UKB WES 150K | 2,091 | 348 | 112 | 6 | 0 | 0 | 0 | 0 |
| | Observed in UKB WES 450K | 3,782 | 1,001 | 513 | 158 | 32 | 2 | 0 | 0 |

[a]For MCPS, Per-gene LOF carrier counts are calculated are calculated after exclusion of variants that fail stringent QC. These include: variants that fail SVM, % missingness > 10%, Mendel errors ≥ 3, or HWE P < 1e-30 with excess heterozygosity (Obs Hets > 1.5x Exp Hets).

For UKB, the following criteria was applied prior to calculation of per-gene LOF carrier counts: i) Genotypes with Depth < 7 for SNPs or Depth < 10 for INDELs were set to missing, ii) variants were excluded if no heterozygote carriers had allele balance above 0.15 for SNPs and 0.20 for INDELs, iii) variants with > 10% missingness were excluded, iv) variants with HWE P < 1e-15 were excluded.

NOTE - the UKB QC prior to mask generation is described in supplement of the 50k paper (https://doi.org/10.1038/s41586-020-2853-0)

**Supplementary Table 5 : Number of coding variants discovered in whole genome sequencing of 9,950 MCPS participants.** Variants were annotated using VEP. Predicted function for each variant was defined as the most deleterious consequence spanning all protein-coding transcripts in Ensembl v100. MAC = Minor Allele Count, IQR = Inter Quartile Range, SD = Standard Deviation.

| Variant category (All transcripts) | N variants (% with MAC=1) | Median number of alternate alleles per participant (IQR) | Mean number of alternate alleles per participant (SD) | Median number of variants per participant (IQR) | Mean number of variants per participant (SD) |
|---|---|---|---|---|---|
| Coding regions | 1460499 (49.04) | 30674 (309.75) | 30683 (249) | 21990 (685) | 21919 (500) |
| **Predicted function** | | | | | |
| In-frame indels | 16695 (47.11) | 300 (16) | 300 (12) | 218 (14) | 218 (10) |
| Synonymous | 484938 (44.67) | 15370 (176.75) | 15373 (138) | 10989 (339) | 10957 (250) |
| Missense | 887628 (50.66) | 14496 (175) | 14500 (135) | 10386 (334) | 10354 (243) |
| Likely benign | 230281 (44.54) | 10707 (128) | 10709 (100) | 7326 (219) | 7308 (158) |
| Possibly deleterious | 496576 (51.2) | 3676 (78) | 3677 (58) | 2942 (121) | 2935 (88) |
| Likely deleterious | 160771 (57.77) | 114 (15) | 114 (11) | 111 (16) | 111 (12) |
| pLOF | 71238 (59.06) | 510 (23) | 510 (17) | 389 (22) | 389 (16) |
| Start lost | 3219 (55.27) | 31 (5) | 31 (4) | 23 (5) | 24 (3) |
| Stop gain | 22679 (59.83) | 109 (10) | 109 (8) | 86 (9) | 86 (7) |
| Stop lost | 1342 (52.76) | 18 (3) | 18 (3) | 13 (3) | 13 (2) |
| Splice donor | 9567 (57.17) | 84 (9) | 84 (7) | 64 (7) | 65 (5) |
| Frameshift | 27856 (59.96) | 203 (16) | 203 (12) | 155 (14) | 155 (10) |
| Splice acceptor | 6575 (58.49) | 65 (7) | 65 (6) | 47 (6) | 47 (4) |

**Supplementary Table 6 : Number of canonical coding variants discovered in whole genome sequencing of 9,950 MCPS participants.** Variants were annotated using VEP. Predicted function for each variant was defined by consequence in canonical coding transcripts in Ensembl v100. MAC = Minor Allele Count, IQR = Inter Quartile Range, SD = Standard Deviation.

| Variant category (All transcripts) | N variants (% with MAC=1) | Median number of alternate alleles per participant (IQR) | Mean number of alternate alleles per participant (SD) | Median number of variants per participant (IQR) | Mean number of variants per participant (SD) |
|---|---|---|---|---|---|
| Coding regions | 1370878 (49.2) | 28252.5 (290.75) | 28260 (233) | 20247 (621.75) | 20182 (456) |
| **Predicted function** | | | | | |
| In-frame indels | 15694 (47.36) | 276 (15) | 276 (11) | 201 (14) | 201 (10) |
| Synonymous | 468904 (44.64) | 14930 (173) | 14933 (136) | 10672 (332) | 10639 (244) |
| Missense | 828706 (50.96) | 12819 (160) | 12822 (124) | 9189 (290) | 9164 (211) |
| Likely benign | 198955 (44.49) | 9460 (119) | 9461 (92) | 6450 (186.75) | 6436 (136) |
| Possibly deleterious | 469321 (51.38) | 3246 (71) | 3248 (54) | 2623 (108) | 2617 (79) |
| Likely deleterious | 160430 (57.77) | 113 (15) | 114 (11) | 111 (16) | 111 (12) |
| pLOF | 57574 (61.35) | 229 (16) | 229 (12) | 178 (14) | 178 (11) |
| Start lost | 1854 (57.23) | 8 (3) | 9 (2) | 7 (2) | 7 (2) |
| Stop gain | 19616 (61.41) | 71 (8) | 71 (6) | 55 (7) | 55 (5) |
| Stop lost | 681 (54.19) | 4 (3) | 5 (2) | 3 (1) | 3 (1) |
| Splice donor | 6623 (60.52) | 25 (5) | 25 (4) | 21 (5) | 21 (3) |
| Frameshift | 23985 (61.9) | 100 (12) | 100 (8) | 77 (9) | 78 (7) |
| Splice acceptor | 4815 (62.08) | 19 (4) | 19 (3) | 14 (3) | 14 (3) |

**Supplementary Table 7**. Number of variants available in MCPS WGS, gnomAD, and TOPMed. MCPS variants were annotated with VEP, with function defined as the most deleterious consequence based on overlap with any protein-coding transcript in Ensembl v100. Variants for TOPMED Freeze 8 and gnomAD v3.1.2 were annotated with SNPeff, with function defined as the most deleterious consequence based on overlap with any protein-coding transcript in Ensembl v85.

| Variant Type | MAF | MCPS WGS All ancestries (N=9950) | | TOPMed Freeze 8[a] All ancestries (N=132,345) | gnomAD 3.1[a] All ancestries (N=76,156) |
|---|---|---|---|---|---|
| | | Total Variants | Unique to MCPS | | |
| Synonymous | All | 484,938 | 97,431 | 2,396,982 | 2,062,223 |
| | Singleton | 216,645 | 74,608 | 996,855 | 981,579 |
| | Doubleton-0.1% | 204,579 | 22,809 | 1,309,070 | 984,865 |
| | 0.1-1% | 37,232 | 12 | 50,742 | 53,974 |
| | 1-5% | 8,328 | 1 | 19,111 | 19,495 |
| | >5% | 18,154 | 1 | 21,204 | 22,310 |
| Missense | All | 887,628 | 227,561 | 5,063,772 | 4,231,655 |
| | Singleton | 449,715 | 177,023 | 2,343,133 | 2,237,366 |
| | Doubleton-0.1% | 362,573 | 50,505 | 2,616,882 | 1,881,080 |
| | 0.1-1% | 48,857 | 25 | 65,209 | 70,941 |
| | 1-5% | 9,332 | 1 | 19,718 | 21,104 |
| | >5% | 17,151 | 7 | 18,830 | 21,164 |
| LOF | All | 71,238 | 28,485 | 507,022 | 426,399 |
| | Singleton | 42,075 | 22,969 | 289,368 | 268,983 |
| | Doubleton-0.1% | 25,462 | 5,502 | 213,540 | 150,603 |
| | 0.1-1% | 2,535 | 10 | 2,795 | 4,554 |
| | 1-5% | 424 | 1 | 730 | 1,192 |
| | >5% | 742 | 3 | 589 | 1,067 |
| All Variants | All | 131,851,585 | 31,533,601 | 705,482,499 | 643,434,862 |
| | Singleton | 57,885,022 | 23,866,451 | 323,651,578 | 317,422,028 |
| | Doubleton-0.1% | 54,213,931 | 7,621,741 | 354,185,173 | 287,971,390 |
| | 0.1-1% | 10,811,295 | 33,835 | 14,418,471 | 20,236,181 |
| | 1-5% | 2,635,764 | 4,868 | 5,928,971 | 8,153,113 |
| | >5% | 6,305,573 | 6,706 | 7,298,306 | 9,652,150 |

**Supplementary Table 8**. Overlap of coding variants captured in canonical coding transcripts by WES and WGS. Variants were annotated with VEP and defined as coding based on overlap with a canonical protein-coding transcript in Ensembl v100. Counts are restricted to the same set of 9,950 individuals with both WGS and WGS available.  All variants passed QC for the respective platform. AAF = Alternate Allele Frequency.

| Class | WES Count | WGS Count | Shared-WES Count | Shared-WGS Count | WES Unique | WGS Unique |
|---|---|---|---|---|---|---|
| All | 1340335 | 1370878 | 1307688 | 1307688 | 32647 | 63190 |
| AAF > 1% | 50430 | 51570 | 49662 | 49674 | 768 | 1896 |
| 0.1% > AAF ≥ 1% | 80163 | 82449 | 79376 | 79410 | 787 | 3039 |
| AAC > 1 & AAF ≤ 0.1% | 548712 | 562420 | 541829 | 541783 | 6883 | 20637 |
| Singleton | 661030 | 674439 | 636821 | 636821 | 24209 | 37618 |

**Supplementary Table 9** : Comparison of WES and WGS datasets in coding genes. Variants were annotated with VEP. Predicted function is defined as the most deleterious consequence for a given variant based on overlap with any protein-coding transcript in Ensembl v100. Counts are restricted to the same set of 9,950 individuals with both WGS and WGS available. All variants passed QC for the respective platform. AAF = Alternate Allele Frequency, IQR = Inter Quartile Range, SD = Standard Deviation.

| Variant category | MCPS WGS - All Coding Regions (N=9950) | | | | | MCPS WES Downsampled - All Coding Regions (N=9950) | | | | |
|---|---|---|---|---|---|---|---|---|---|---|
| (All transcripts) | # Variants (All AAF) | Median number of alternate alleles per participant (IQR) | Mean number of alternate alleles per participant (SD) | Median number of unique variants per participant (IQR) | Mean number of unique variants per participant (SD) | # Variants (All AAF) | Median number of alternate alleles per participant (IQR) | Mean number of alternate alleles per participant (SD) | Median number of unique variants per participant (IQR) | Mean number of unique variants per participant (SD) |
| Coding regions | 1460499 | 30674 (309.75) | 30683 (249) | 21990 (685) | 21919 (500) | 1396076 | 29062 (295) | 29069 (238) | 20839.5 (634) | 20770 (467) |
| **Predicted function** | | | | | | | | | | |
| In-frame indels | 16695 | 300 (16) | 300 (12) | 218 (14) | 218 (10) | 15540 | 281 (16) | 281 (12) | 207 (13) | 207 (10) |
| Synonymous | 484938 | 15370 (176.75) | 15373 (138) | 10989 (339) | 10957 (250) | 469121 | 14882.5 (173) | 14886 (136) | 10640.5 (323.75) | 10609 (240) |
| Missense | 887628 | 14496 (175) | 14500 (135) | 10386 (334) | 10354 (243) | 849431 | 13545.5 (167) | 13548 (128) | 9709 (305.75) | 9681 (223) |
| Likely benign | 230281 | 10707 (128) | 10709 (100) | 7326 (219) | 7308 (158) | 203447 | 9868 (120) | 9869 (93) | 6731 (195) | 6716 (142) |
| Possibly deleterious | 496576 | 3676 (78) | 3677 (58) | 2942 (121) | 2935 (88) | 484611 | 3563 (76) | 3564 (56) | 2859 (116) | 2853 (84) |
| Likely deleterious | 160771 | 114 (15) | 114 (11) | 111 (16) | 111 (12) | 161373 | 114 (15) | 115 (12) | 112 (16) | 112 (12) |
| pLOF | 71238 | 510 (23) | 510 (17) | 389 (22) | 389 (16) | 61984 | 354 (20) | 354 (15) | 273 (19) | 273 (14) |
| Start lost | 3219 | 31 (5) | 31 (4) | 23 (5) | 24 (3) | 2944 | 27 (5) | 27 (4) | 21 (4) | 21 (3) |
| Stop gain | 22679 | 109 (10) | 109 (8) | 86 (9) | 86 (7) | 20988 | 84 (9) | 84 (7) | 67 (8) | 67 (6) |
| Stop lost | 1342 | 18 (3) | 18 (3) | 13 (3) | 13 (2) | 1068 | 13 (3) | 13 (3) | 10 (3) | 10 (2) |
| Splice donor | 9567 | 84 (9) | 84 (7) | 64 (7) | 65 (5) | 6890 | 38 (6) | 38 (5) | 30 (5) | 30 (4) |
| Frameshift | 27856 | 203 (16) | 203 (12) | 155 (14) | 155 (10) | 24756 | 146 (14) | 146 (10) | 114 (13) | 114 (9) |
| Splice acceptor | 6575 | 65 (7) | 65 (6) | 47 (6) | 47 (4) | 5338 | 44 (6) | 44 (5) | 32 (5) | 32 (3) |

**Supplementary Table 10**. Overlap of coding variants captured in all coding regions by WES and WGS. Variants were annotated with VEP and defined as coding based on overlap with any protein-coding transcript in Ensembl v100. Counts are restricted to the same set of 9,950 individuals with both WGS and WGS available.  All variants passed QC for the respective platform. AAF = Alternate Allele Frequency.

| Class | WES Count | WGS Count | Shared-WES Count | Shared-WGS Count | WES Unique | WGS Unique |
|---|---|---|---|---|---|---|
| All | 1396076 | 1460499 | 1361790 | 1361790 | 34286 | 98709 |
| AAF > 1% | 53047 | 55991 | 52243 | 52262 | 804 | 3729 |
| 0.1% > AAF ≥ 1% | 84010 | 88821 | 83176 | 83215 | 834 | 5606 |
| AAC > 1 & AAF ≤ 0.1% | 571756 | 599420 | 564526 | 564479 | 7230 | 34941 |
| Singleton | 687263 | 716267 | 661845 | 661834 | 25418 | 54433 |

**Supplementary Table 11** : Comparison of WES and WGS datasets in coding genes. Variants were annotated with VEP. Predicted function is defined by canonical transcript consequence in Ensembl v100. Counts are restricted to the same set of 9,950 individuals with both WGS and WGS available. Only variants within the WES targeted capture region were evaluated. All variants passed QC for the respective platform. AAF = Alternate Allele Frequency, IQR = Inter Quartile Range, SD = Standard Deviation.

| Variant category (Canonical transcripts) | MCPS WGS - WES Capture Region (N=9950) | | | | | MCPS WES Downsampled - WES Capture Region (N=9950) | | | | |
|---|---|---|---|---|---|---|---|---|---|---|
| | # Variants (All AAF) | Median number of alternate alleles per participant (IQR) | Mean number of alternate alleles per participant (SD) | Median number of unique variants per participant (IQR) | Mean number of unique variants per participant (SD) | # Variants (All AAF) | Median number of alternate alleles per participant (IQR) | Mean number of alternate alleles per participant (SD) | Median number of unique variants per participant (IQR) | Mean number of unique variants per participant (SD) |
| Coding regions | 1339880 | 27529 (288) | 27536 (229) | 19734 (604.75) | 19671 (444) | 1331652 | 27497 (287) | 27503 (229) | 19722 (597) | 19657 (440) |
| **Predicted function** | | | | | | | | | | |
| In-frame indels | 15283 | 266 (15) | 266 (11) | 194 (13) | 194 (10) | 14799 | 264 (15) | 264 (11) | 195 (13) | 195 (9) |
| Synonymous | 461814 | 14653 (171) | 14657 (134) | 10477 (326) | 10445 (241) | 460764 | 14661 (170) | 14665 (134) | 10483 (325.75) | 10452 (239) |
| Missense | 814192 | 12429.5 (161) | 12432 (121) | 8917 (276.75) | 8892 (204) | 808837 | 12395 (157) | 12397 (121) | 8897 (277) | 8873 (202) |
| Likely benign | 188289 | 9103 (118) | 9104 (90) | 6203 (176) | 6189 (129) | 185067 | 9032 (115) | 9032 (89) | 6154.5 (174) | 6141 (127) |
| Possibly deleterious | 465663 | 3213 (71) | 3215 (53) | 2598 (106) | 2592 (78) | 462652 | 3250 (72) | 3251 (54) | 2626 (107) | 2620 (78) |
| Likely deleterious | 160240 | 113 (15) | 113 (11) | 111 (15) | 111 (12) | 161118 | 114 (16) | 114 (12) | 111 (15) | 112 (12) |
| pLOF | 48591 | 180 (14) | 180 (11) | 139 (12) | 139 (9) | 47252 | 176 (15) | 176 (11) | 137 (13) | 138 (9) |
| Start lost | 1782 | 8 (3) | 8 (2) | 7 (2) | 7 (2) | 1800 | 8 (3) | 8 (2) | 7 (2) | 7 (2) |
| Stop gain | 19143 | 64 (8) | 64 (6) | 49 (7) | 49 (5) | 18864 | 60 (8) | 60 (6) | 47 (7) | 47 (5) |
| Stop lost | 636 | 4 (2) | 4 (2) | 3 (1) | 3 (1) | 623 | 4 (2) | 4 (2) | 3 (1) | 3 (1) |
| Splice donor | 2485 | 8 (3) | 8 (2) | 6 (2) | 6 (2) | 2447 | 5 (2) | 5 (2) | 4 (2) | 4 (2) |
| Frameshift | 23372 | 87 (10) | 87 (8) | 67 (9) | 68 (7) | 22360 | 89 (11) | 90 (8) | 70 (10) | 70 (7) |
| Splice acceptor | 1173 | 9 (2) | 9 (2) | 6 (2) | 6 (1) | 1158 | 9 (2) | 9 (2) | 6 (2) | 6 (1) |

**Supplementary Table 12**. Overlap of coding variants captured in canonical coding transcripts within WES targeted capture regions by WES and WGS. Variants were annotated with VEP and defined as coding based on overlap with a canonical protein-coding transcript in Ensembl v100. Counts are restricted to the same set of 9,950 individuals with both WGS and WGS available. All variants passed QC for the respective platform. AAF = Alternate Allele Frequency.

| Class | WES Count | WGS Count | Shared-WES Count | Shared-WGS Count | WES Unique | WGS Unique |
|---|---|---|---|---|---|---|
| All | 1331652 | 1339880 | 1299487 | 1299487 | 32165 | 40393 |
| AAF > 1% | 50238 | 50265 | 49489 | 49501 | 749 | 764 |
| 0.1% > AAF ≥ 1% | 79824 | 80476 | 79057 | 79088 | 767 | 1388 |
| AAC > 1 & AAF ≤ 0.1% | 545591 | 549818 | 538833 | 538785 | 6758 | 11033 |
| Singleton | 655999 | 659321 | 632108 | 632113 | 23891 | 27208 |

**Supplementary Table 13** : Comparison of WES and WGS datasets in coding genes. Variants were annotated with VEP. Predicted function is defined as the most deleterious consequence for a given variant based on overlap with any protein-coding transcript in Ensembl v100. Counts are restricted to the same set of 9,950 individuals with both WGS and WGS available. Only variants within the WES targeted capture region were evaluated. All variants passed QC for the respective platform. AAF = Alternate Allele Frequency, IQR = Inter Quartile Range, SD = Standard Deviation.

| Variant category | MCPS WGS - WES Capture Region (N=9950) | | | | | MCPS WES Downsampled - WES Capture Region (N=9950) | | | | |
|---|---|---|---|---|---|---|---|---|---|---|
| (All transcripts) | # Variants (All AAF) | Median number of alternate alleles per participant (IQR) | Mean number of alternate alleles per participant (SD) | Median number of unique variants per participant (IQR) | Mean number of unique variants per participant (SD) | # Variants (All AAF) | Median number of alternate alleles per participant (IQR) | Mean number of alternate alleles per participant (SD) | Median number of unique variants per participant (IQR) | Mean number of unique variants per participant (SD) |
| Coding regions | 1382760 | 28638 (293.75) | 28644 (236) | 20526 (632) | 20461 (464) | 1374220 | 28601 (292) | 28607 (235) | 20512 (626) | 20443 (459) |
| **Predicted function** | | | | | | | | | | |
| In-frame indels | 15823 | 278 (16) | 278 (11) | 203 (13) | 204 (10) | 15308 | 277 (15) | 277 (12) | 203 (13) | 204 (10) |
| Synonymous | 466529 | 14753 (170.75) | 14756 (135) | 10548 (326) | 10518 (240) | 465402 | 14757 (172) | 14761 (135) | 10554 (323) | 10523 (239) |
| Missense | 845964 | 13329 (167) | 13331 (127) | 9553 (300) | 9525 (221) | 840475 | 13287.5 (164) | 13290 (126) | 9528 (300) | 9501 (219) |
| Likely benign | 201228 | 9743 (122) | 9744 (93) | 6643 (195) | 6628 (141) | 197948 | 9662 (119) | 9663 (92) | 6589 (192) | 6573 (139) |
| Possibly deleterious | 484156 | 3472 (74) | 3474 (56) | 2793 (113) | 2786 (83) | 481192 | 3511 (75) | 3512 (56) | 2822 (114) | 2816 (83) |
| Likely deleterious | 160580 | 113 (15) | 114 (11) | 111 (16) | 111 (12) | 161335 | 114 (15) | 115 (12) | 112 (16) | 112 (12) |
| pLOF | 54444 | 278 (18) | 278 (13) | 214 (16) | 214 (12) | 53035 | 278 (18) | 278 (13) | 215 (17) | 216 (12) |
| Start lost | 2855 | 23 (5) | 23 (4) | 18 (4) | 18 (3) | 2874 | 24 (5) | 24 (4) | 19 (4) | 19 (3) |
| Stop gain | 20942 | 80 (10) | 80 (7) | 63 (8) | 63 (6) | 20649 | 76 (9) | 76 (7) | 61 (8) | 61 (6) |
| Stop lost | 920 | 12 (3) | 13 (2) | 9 (2) | 9 (2) | 907 | 12 (3) | 12 (2) | 9 (2) | 9 (2) |
| Splice donor | 2999 | 19 (4) | 19 (3) | 14 (3) | 14 (2) | 2964 | 17 (4) | 17 (3) | 13 (3) | 13 (2) |
| Frameshift | 25105 | 120 (12) | 120 (10) | 93 (12) | 93 (8) | 24025 | 123 (13) | 123 (10) | 96 (11) | 96 (8) |
| Splice acceptor | 1623 | 23 (4) | 23 (3) | 16 (3) | 16 (2) | 1616 | 25 (4) | 25 (3) | 18 (3) | 18 (2) |

**Supplementary Table 14**. Overlap of coding variants captured in WES targeted capture regions by WES and WGS. Variants were annotated with VEP and defined as coding based on overlap with any protein-coding transcript in Ensembl v100. Counts are restricted to the same set of 9,950 individuals with both WGS and WGS available. Only variants within the WES targeted capture region were evaluated. All variants passed QC for the respective platform. AAF = Alternate Allele Frequency.

| Class | WES Count | WGS Count | Shared-WES Count | Shared-WGS Count | WES Unique | WGS Unique |
|---|---|---|---|---|---|---|
| All | 1374220 | 1382760 | 1340879 | 1340879 | 33341 | 41881 |
| AAF > 1% | 52223 | 52269 | 51454 | 51470 | 769 | 799 |
| 0.1% > AAF ≥ 1% | 82758 | 83432 | 81962 | 81990 | 796 | 1442 |
| AAC > 1 & AAF ≤ 0.1% | 563087 | 567480 | 556059 | 556007 | 7028 | 11473 |
| Singleton | 676152 | 679579 | 651404 | 651412 | 24748 | 28167 |

**Supplementary Table 15** : Comparison of WES and WGS datasets in coding genes for variants with AAF ≤ 1%. Variants were annotated with VEP. Predicted function is defined by canonical transcript consequence in Ensembl v100. Counts are restricted to the same set of 9,950 individuals with both WGS and WGS available. All variants passed QC for the respective platform. AAF = Alternate Allele Frequency, IQR = Inter Quartile Range, SD = Standard Deviation.

| Variant category (Canonical transcripts) | MCPS WGS - All Coding Regions (N=9950) | | | | | MCPS WES Downsampled - All Coding Regions (N=9950) | | | | |
|---|---|---|---|---|---|---|---|---|---|---|
| | # Variants (AAF ≤ 1%) | Median number of alternate alleles per participant (IQR) | Mean number of alternate alleles per participant (SD) | Median number of unique variants per participant (IQR) | Mean number of unique variants per participant (SD) | # Variants (AAF ≤ 1%) | Median number of alternate alleles per participant (IQR) | Mean number of alternate alleles per participant (SD) | Median number of unique variants per participant (IQR) | Mean number of unique variants per participant (SD) |
| Coding regions | 1319308 | 791 (323) | 799 (235) | 787 (320) | 794 (233) | 1289905 | 772 (312) | 779 (228) | 767 (310.75) | 773 (226) |
| **Predicted function** | | | | | | | | | | |
| In-frame indels | 15136 | 10 (5) | 10 (4) | 9 (5) | 10 (4) | 14383 | 9 (5) | 9 (4) | 9 (6) | 9 (4) |
| Synonymous | 442659 | 320 (151) | 326 (113) | 318 (149.75) | 323 (112) | 435531 | 313 (147) | 319 (111) | 311 (147) | 316 (110) |
| Missense | 804517 | 440 (163) | 443 (117) | 437 (163) | 440 (117) | 786436 | 429 (159) | 432 (113) | 427 (157) | 429 (113) |
| Likely benign | 185049 | 130 (61) | 133 (46) | 129 (61) | 131 (45) | 172388 | 122 (58) | 124 (43) | 121 (57) | 123 (43) |
| Possibly deleterious | 459722 | 246 (89) | 248 (64) | 245 (88) | 246 (63) | 453580 | 243 (88) | 245 (63) | 242 (87) | 243 (62) |
| Likely deleterious | 159746 | 62 (17) | 63 (13) | 62 (18) | 62 (13) | 160468 | 63 (17) | 63 (13) | 62 (17) | 63 (13) |
| pLOF | 56996 | 20 (7) | 21 (6) | 20 (8) | 21 (6) | 53555 | 19 (8) | 19 (5) | 19 (7) | 19 (5) |
| Start lost | 1829 | 1 (1) | 1 (1) | 1 (1) | 1 (1) | 1792 | 1 (1) | 1 (1) | 1 (1) | 1 (1) |
| Stop gain | 19444 | 6 (3) | 7 (3) | 6 (3) | 7 (3) | 18744 | 6 (4) | 6 (3) | 6 (4) | 6 (3) |
| Stop lost | 670 | 0 (1) | 0 (1) | 0 (1) | 0 (1) | 634 | 0 (1) | 0 (1) | 0 (1) | 0 (1) |
| Splice donor | 6554 | 2 (3) | 3 (2) | 2 (3) | 3 (2) | 5819 | 2 (2) | 2 (2) | 2 (2) | 2 (2) |
| Frameshift | 23729 | 8 (5) | 9 (3) | 8 (5) | 9 (3) | 22178 | 8 (4) | 8 (3) | 8 (4) | 8 (3) |
| Splice acceptor | 4770 | 1 (1) | 2 (1) | 1 (1) | 2 (1) | 4388 | 1 (1) | 2 (1) | 1 (1) | 2 (1) |

**Supplementary Table 16** : Comparison of WES and WGS datasets in coding genes for variants with AAF ≤ 1%. Variants were annotated with VEP. Predicted function is defined as the most deleterious consequence for a given variant based on overlap with any protein-coding transcript in Ensembl v100. Counts are restricted to the same set of 9,950 individuals with both WGS and WGS available. All variants passed QC for the respective platform. AAF = Alternate Allele Frequency, IQR = Inter Quartile Range, SD = Standard Deviation.

| Variant category (All transcripts) | MCPS WGS - All Coding Regions (N=9950) | | | | | MCPS WES Downsampled - All Coding Regions (N=9950) | | | | |
|---|---|---|---|---|---|---|---|---|---|---|
| | # Variants (AAF ≤ 1%) | Median number of alternate alleles per participant (IQR) | Mean number of alternate alleles per participant (SD) | Median number of unique variants per participant (IQR) | Mean number of unique variants per participant (SD) | # Variants (AAF ≤ 1%) | Median number of alternate alleles per participant (IQR) | Mean number of alternate alleles per participant (SD) | Median number of unique variants per participant (IQR) | Mean number of unique variants per participant (SD) |
| Coding regions | 1404508 | 851 (351) | 859 (255) | 846 (349) | 853 (253) | 1343029 | 809 (329) | 816 (240) | 804 (326) | 810 (238) |
| **Predicted function** | | | | | | | | | | |
| In-frame indels | 16091 | 10 (5) | 11 (4) | 10 (5) | 11 (4) | 14965 | 9 (5) | 10 (4) | 9 (5) | 10 (4) |
| Synonymous | 457879 | 331 (157) | 337 (117) | 329 (156) | 334 (116) | 442953 | 319 (149) | 324 (113) | 317 (150) | 322 (112) |
| Missense | 860477 | 479 (182) | 483 (131) | 476 (181) | 480 (130) | 823980 | 455 (170) | 458 (122) | 452 (169) | 455 (121) |
| Likely benign | 214372 | 151 (72) | 154 (54) | 150 (71) | 153 (53) | 189013 | 133 (63) | 136 (47) | 133 (62) | 135 (47) |
| Possibly deleterious | 486022 | 264 (98) | 266 (69) | 262 (96.75) | 264 (69) | 474285 | 257 (94) | 259 (67) | 255 (93) | 257 (66) |
| Likely deleterious | 160083 | 62 (17) | 63 (13) | 62 (18) | 63 (13) | 160682 | 63 (18) | 63 (13) | 62 (17) | 63 (13) |
| pLOF | 70061 | 28 (11) | 29 (8) | 28 (11) | 29 (8) | 61131 | 23 (9) | 24 (7) | 23 (9) | 24 (7) |
| Start lost | 3142 | 1 (1) | 1 (1) | 1 (1) | 1 (1) | 2874 | 1 (2) | 1 (1) | 1 (2) | 1 (1) |
| Stop gain | 22407 | 8 (5) | 9 (3) | 8 (5) | 8 (3) | 20777 | 7 (5) | 8 (3) | 7 (4) | 8 (3) |
| Stop lost | 1308 | 0 (1) | 1 (1) | 0 (1) | 1 (1) | 1044 | 0 (1) | 1 (1) | 0 (1) | 1 (1) |
| Splice donor | 9369 | 4 (3) | 4 (2) | 4 (3) | 4 (2) | 6800 | 3 (3) | 3 (2) | 3 (3) | 3 (2) |
| Frameshift | 27380 | 11 (6) | 11 (4) | 11 (6) | 11 (4) | 24383 | 9 (5) | 9 (4) | 9 (5) | 9 (4) |
| Splice acceptor | 6455 | 3 (3) | 3 (2) | 3 (3) | 3 (2) | 5253 | 2 (2) | 2 (1) | 2 (2) | 2 (1) |

**Supplementary Table 17** : Comparison of WES and WGS datasets in coding genes for variants with AAF ≤ 1%. Variants were annotated with VEP. Predicted function is defined by canonical transcript consequence in Ensembl v100. Counts are restricted to the same set of 9,950 individuals with both WGS and WGS available. Only variants within the WES targeted capture region were evaluated. All variants passed QC for the respective platform. AAF = Alternate Allele Frequency, IQR = Inter Quartile Range, SD = Standard Deviation.

| Variant category (Canonical transcripts) | MCPS WGS - WES Capture Region (N=9950) | | | | | MCPS WES Downsampled - WES Capture Region (N=9950) | | | | |
|---|---|---|---|---|---|---|---|---|---|---|
| | # Variants (AAF ≤ 1%) | Median number of alternate alleles per participant (IQR) | Mean number of alternate alleles per participant (SD) | Median number of unique variants per participant (IQR) | Mean number of unique variants per participant (SD) | # Variants (AAF ≤ 1%) | Median number of alternate alleles per participant (IQR) | Mean number of alternate alleles per participant (SD) | Median number of unique variants per participant (IQR) | Mean number of unique variants per participant (SD) |
| Coding regions | 1289615 | 772 (315) | 781 (229) | 768 (312) | 775 (227) | 1281414 | 769 (311) | 776 (227) | 763 (309) | 770 (225) |
| **Predicted function** | | | | | | | | | | |
| In-frame indels | 14745 | 9 (5) | 10 (4) | 9 (5) | 9 (4) | 14262 | 9 (5) | 9 (4) | 9 (5) | 9 (4) |
| Synonymous | 436028 | 314 (148) | 320 (111) | 312 (147) | 318 (110) | 434993 | 313 (147) | 319 (111) | 311 (146) | 316 (110) |
| Missense | 790696 | 431 (159) | 434 (114) | 428 (159) | 431 (114) | 785353 | 429 (158) | 431 (113) | 426 (158) | 428 (112) |
| Likely benign | 174981 | 123 (59) | 126 (44) | 122 (57) | 125 (43) | 171886 | 121 (57) | 124 (43) | 120 (57) | 122 (42) |
| Possibly deleterious | 456157 | 244 (88) | 246 (63) | 243 (87.75) | 244 (63) | 453036 | 243 (88) | 245 (63) | 242 (87) | 243 (62) |
| Likely deleterious | 159558 | 62 (18) | 63 (13) | 62 (18) | 62 (13) | 160431 | 63 (17) | 63 (13) | 62 (17) | 63 (13) |
| pLOF | 48146 | 17 (6) | 17 (5) | 17 (7) | 17 (5) | 46806 | 16 (7) | 17 (5) | 16 (7) | 17 (5) |
| Start lost | 1758 | 0 (1) | 1 (1) | 0 (1) | 1 (1) | 1776 | 0 (1) | 1 (1) | 0 (1) | 1 (1) |
| Stop gain | 18987 | 6 (4) | 6 (3) | 6 (4) | 6 (3) | 18712 | 6 (4) | 6 (3) | 6 (4) | 6 (3) |
| Stop lost | 626 | 0 (1) | 0 (1) | 0 (1) | 0 (1) | 615 | 0 (1) | 0 (1) | 0 (1) | 0 (1) |
| Splice donor | 2472 | 1 (1) | 1 (1) | 1 (1) | 1 (1) | 2436 | 1 (1) | 1 (1) | 1 (1) | 1 (1) |
| Frameshift | 23144 | 8 (4) | 8 (3) | 8 (4) | 8 (3) | 22123 | 8 (4) | 8 (3) | 8 (4) | 8 (3) |
| Splice acceptor | 1159 | 0 (1) | 0 (1) | 0 (1) | 0 (1) | 1144 | 0 (1) | 0 (1) | 0 (1) | 0 (1) |

**Supplementary Table 18** : Comparison of WES and WGS datasets in coding genes for variants with AAF ≤ 1%. Variants were annotated with VEP. Predicted function is defined as the most deleterious consequence for a given variant based on overlap with any protein-coding transcript in Ensembl v100. Counts are restricted to the same set of 9,950 individuals with both WGS and WGS available. Only variants within the WES targeted capture region were evaluated. All variants passed QC for the respective platform. AAF = Alternate Allele Frequency, IQR = Inter Quartile Range, SD = Standard Deviation.

| Variant category (All transcripts) | MCPS WGS - WES Capture Region (N=9950) | | | | | MCPS WES Downsampled - WES Capture Region (N=9950) | | | | |
|---|---|---|---|---|---|---|---|---|---|---|
| | # Variants (AAF ≤ 1%) | Median number of alternate alleles per participant (IQR) | Mean number of alternate alleles per participant (SD) | Median number of unique variants per participant (IQR) | Mean number of unique variants per participant (SD) | # Variants (AAF ≤ 1%) | Median number of alternate alleles per participant (IQR) | Mean number of alternate alleles per participant (SD) | Median number of unique variants per participant (IQR) | Mean number of unique variants per participant (SD) |
| Coding regions | 1330491 | 800.5 (327) | 809 (238) | 797 (324) | 803 (236) | 1321997 | 797 (324) | 803 (236) | 791 (322) | 797 (234) |
| **Predicted function** | | | | | | | | | | |
| In-frame indels | 15260 | 10 (6) | 10 (4) | 10 (6) | 10 (4) | 14748 | 9 (5) | 10 (4) | 9 (5) | 9 (4) |
| Synonymous | 440533 | 318 (150) | 323 (113) | 316 (149) | 321 (112) | 439434 | 316 (148) | 322 (112) | 314 (148) | 319 (111) |
| Missense | 820923 | 452 (168) | 455 (121) | 449 (168) | 452 (120) | 815453 | 449 (168) | 452 (120) | 446 (167) | 449 (119) |
| Likely benign | 186978 | 132 (63) | 135 (47) | 131 (61) | 133 (46) | 183830 | 130 (61) | 132 (46) | 129 (61) | 131 (46) |
| Possibly deleterious | 474051 | 255 (94) | 258 (67) | 254 (93) | 256 (66) | 470979 | 254 (93) | 256 (66) | 253 (92) | 255 (66) |
| Likely deleterious | 159894 | 62 (17) | 63 (13) | 62 (18) | 62 (13) | 160644 | 63 (18) | 63 (13) | 62 (17) | 63 (13) |
| pLOF | 53775 | 20 (8) | 20 (6) | 20 (8) | 20 (6) | 52362 | 20 (8) | 20 (6) | 19 (8) | 20 (6) |
| Start lost | 2794 | 1 (2) | 1 (1) | 1 (2) | 1 (1) | 2812 | 1 (2) | 1 (1) | 1 (2) | 1 (1) |
| Stop gain | 20739 | 7 (4) | 7 (3) | 7 (4) | 7 (3) | 20451 | 7 (4) | 7 (3) | 7 (4) | 7 (3) |
| Stop lost | 897 | 0 (1) | 0 (1) | 0 (1) | 0 (1) | 886 | 0 (1) | 0 (1) | 0 (1) | 0 (1) |
| Splice donor | 2966 | 1 (2) | 1 (1) | 1 (2) | 1 (1) | 2932 | 1 (2) | 1 (1) | 1 (2) | 1 (1) |
| Frameshift | 24793 | 9 (4) | 9 (4) | 9 (4) | 9 (4) | 23706 | 9 (5) | 9 (3) | 9 (5) | 9 (3) |
| Splice acceptor | 1586 | 1 (1) | 1 (1) | 1 (1) | 1 (1) | 1575 | 1 (1) | 1 (1) | 1 (1) | 1 (1) |

**Supplementary Table 19**: Number (and %) of coding variants captured only by WES or WGS that are represented in TOPMED Freeze 8 or Gnomad v3.1.2. Variants were annotated with VEP and defined as coding based on the canonical protein-coding transcript in Ensembl v100. Counts are restricted to the same set of 9,950 individuals with both WGS and WGS available. Only variants within the WES targeted capture region were evaluated. All variants passed QC for the respective platform. AAF = Alternate Allele Frequency.

| Source | AAF | N | In Topmed | In Gnomad | In Either |
|---|---|---|---|---|---|
| WGS unique | All | 63190 | 33356 (52.79) | 34378 (54.4) | 40661 (64.35) |
| | AAF > 1% | 1896 | 1587 (83.7) | 1875 (98.89) | 1883 (99.31) |
| | AAF ≤ 1% | 61294 | 31769 (51.83) | 32503 (53.03) | 38778 (63.27) |
| | Singleton | 37618 | 15199 (40.4) | 14346 (38.14) | 18266 (48.56) |
| WES unique | All | 32647 | 16643 (50.98) | 17558 (53.78) | 21507 (65.88) |
| | AAF > 1% | 768 | 290 (37.76) | 703 (91.54) | 714 (92.97) |
| | AAF ≤ 1% | 31879 | 16353 (51.3) | 16855 (52.87) | 20793 (65.22) |
| | Singleton | 24209 | 12259 (50.64) | 11612 (47.97) | 14668 (60.59) |

**Supplementary Table 20:** Number (and %) of coding variants captured only by WES or WGS that are represented in TOPMED Freeze 8 or Gnomad v3.1.2. Variants were annotated with VEP and defined as coding based on overlap with any protein-coding transcript in Ensembl v100. Counts are restricted to the same set of 9,950 individuals with both WGS and WGS available. Only variants within the WES targeted capture region were evaluated. All variants passed QC for the respective platform. AAF = Alternate Allele Frequency.

| Source | AAF | N | In Topmed | In Gnomad | In Either |
|---|---|---|---|---|---|
| WGS unique | All | 98709 | 57997 (58.76) | 57065 (57.81) | 67490 (68.37) |
| | AAF > 1% | 3729 | 3342 (89.62) | 3710 (99.49) | 3720 (99.76) |
| | AAF ≤ 1% | 94980 | 54655 (57.54) | 53355 (56.17) | 63770 (67.14) |
| | Singleton | 54433 | 24172 (44.41) | 22183 (40.75) | 28282 (51.96) |
| WES unique | All | 34286 | 17504 (51.05) | 18471 (53.87) | 22613 (65.95) |
| | AAF > 1% | 804 | 304 (37.81) | 738 (91.79) | 749 (93.16) |
| | AAF ≤ 1% | 33482 | 17200 (51.37) | 17733 (52.96) | 21864 (65.3) |
| | Singleton | 25418 | 12896 (50.74) | 12213 (48.05) | 15419 (60.66) |

**Supplementary Table 21**: Number (and %) of target region canonical coding variants captured only by WES or WGS that are represented in TOPMED Freeze 8 or Gnomad v3.1.2. Variants were annotated with VEP and defined as coding based on the canonical protein-coding transcript in Ensembl v100. Counts are restricted to the same set of 9,950 individuals with both WGS and WGS available. Only variants within the WES targeted capture region were evaluated. All variants passed QC for the respective platform. AAF = Alternate Allele Frequency.

| Source | AAF | N | In Topmed | In Gnomad | In Either |
|---|---|---|---|---|---|
| WGS unique | All | 40393 | 17406 (43.09) | 19515 (48.31) | 23254 (57.57) |
| | AAF > 1% | 764 | 513 (67.15) | 752 (98.43) | 756 (98.95) |
| | AAF ≤ 1% | 39629 | 16893 (42.63) | 18763 (47.35) | 22498 (56.77) |
| | Singleton | 27208 | 9552 (35.11) | 9356 (34.39) | 11970 (43.99) |
| WES unique | All | 32165 | 16479 (51.23) | 17320 (53.85) | 21217 (65.96) |
| | AAF > 1% | 749 | 288 (38.45) | 687 (91.72) | 697 (93.06) |
| | AAF ≤ 1% | 31416 | 16191 (51.54) | 16633 (52.94) | 20520 (65.32) |
| | Singleton | 23891 | 12154 (50.87) | 11498 (48.13) | 14516 (60.76) |

**Supplementary Table 22**: Number (and %) of target region coding variants captured only by WES or WGS that are represented in TOPMED Freeze 8 or Gnomad v3.1.2. Variants were annotated with VEP and defined as coding based on overlap with any protein-coding transcript in Ensembl v100. Counts are restricted to the same set of 9,950 individuals with both WGS and WGS available. Only variants within the WES targeted capture region were evaluated. All variants passed QC for the respective platform. AAF = Alternate Allele Frequency.

| Source | AAF | N | In Topmed | In Gnomad | In Either |
|---|---|---|---|---|---|
| WGS unique | All | 41881 | 18049 (43.1) | 20247 (48.34) | 24136 (57.63) |
| | AAF > 1% | 799 | 528 (66.08) | 786 (98.37) | 791 (99) |
| | AAF ≤ 1% | 41082 | 17521 (42.65) | 19461 (47.37) | 23345 (56.83) |
| | Singleton | 28167 | 9897 (35.14) | 9673 (34.34) | 12396 (44.01) |
| WES unique | All | 33341 | 17090 (51.26) | 17970 (53.9) | 22003 (65.99) |
| | AAF > 1% | 769 | 297 (38.62) | 707 (91.94) | 717 (93.24) |
| | AAF ≤ 1% | 32572 | 16793 (51.56) | 17263 (53) | 21286 (65.35) |
| | Singleton | 24748 | 12598 (50.91) | 11919 (48.16) | 15039 (60.77) |

**Supplementary Table 23**: Concordance comparison of WES, WGS and ARRAY datasets

| Comparison | # samples | #autosomal SNPs | # SNPs with discordance > 0 | # SNPs with discordance > 1% | mean discordance % |
|---|---|---|---|---|---|
| WES vs WGS | 9950 | 3,005,612 | 107,998 | 5,935 | 0.00006388 |
| WES vs ARRAY | 138,200 | 47,261 | 25,528 | 350 | 0.03239 |
| WGS vs ARRAY | 9,950 | 498,929 | 93,994 | 1,024 | 0.03233 |

**Supplementary Table 24:** QC steps applied to MCPS array data. Input array data from the RGC Sequencing Lab consisted of 140,831 samples and 650,380 variants and were passed through multiple steps of quality control. The number of samples (N) and number of variants (M) of array data at each step are reported in columns 3 and 4, respectively. A set of 81,747 unrelated samples up to the 3rd degree, used at step 7, was defined in the

| Step | PLINK command | N | M | Comments |
|---|---|---|---|---|
| 0 | | 140,831 | 650,380 | Input data from the RGC Sequencing Lab |
| 1 | --split-x hg38 no-fail | | | no PAR1/2 variants found |
| 2 | --set-hh-missing | | | 46,014 samples, 26,603 variants affected |
| 3 | --zero-cluster samples-sex-discordant --within variants-chr-23-24-25 | | | 128 samples, 30,100 variants affected |
| 4 | (manually check for sex-discordant samples) | 140,831 | 650,380 | No discordant samples found |
| 5 | --autosome --mind 0.05 | (-2,320) | | |
| 6 | --geno 0.02 --mac 1 | | (-80,280) | |
| 7 | --keep-fam unrel.fam --hwe 1e-30 | | (-9,645) | |
| | --make-bed | 138,511 | 560,455 | |
| 8 | --fam nuc.fam --set-me-missing --mendel-duos | | | 244,100 Mendel errors fixed |
| 9 | Repeat steps 5-7 | 138,511 | 559,923 | |

**Supplementary Table 25** : Comparison of family network sizes in MCPS, GHS and UKB datasets. Numbers in brackets are percentages of MCPS participants included in pairwise relationships.

| | MCPS 150k | GHS 145k | UKB 450k |
|---|---|---|---|
| Parent-child relationships | 31,597 (33.1%) | 29,599 (31.1%) | 6,270 (2.3%) |
| Full-sibling relationships | 29,482 (26.9%) | 18,540 (18.9%) | 22,657 (8.5%) |
| 2nd-degree relationships | 47,080 (31.2%) | 71,766 (42.4%) | 11,169 (4.2%) |
| 3rd-degree relationships | 120,180 (45.2%) | 129,782 (48.0%) | 66,847 (19.8%) |
| Number of pedigrees | 22,766 | 19,770 | 24,560 |
| Pedigrees with >2 individuals | 9,889 | 7,998 | 2590 |
| Individuals with both parents (trios) | 5,603 | 5,448 | 1,065 |
| Pedigrees >2 generations | 600 | 2,299 | 0 |

**Supplementary Table 26** : Distribution of family sizes in the 3,595 nuclear familes inferred from the genetic relatedness estiimation

| number of children | number of families |
|---|---|
| 1 | 2268 |
| 2 | 869 |
| 3 | 208 |
| 4 | 100 |
| 5 | 34 |
| 6 | 11 |
| 7 | 3 |
| 8 | 2 |

**Supplementary Table 27** : Summary of relationship types stratified by residence location for each pair of individuals.

| Relationship Type | Residence | Count | % of relationship type |
|---|---|---|---|
| Unrelated parents | Coyoacán | 1354 | 38% |
| | Iztapalapa | 2211 | 62% |
| | Mixed | 30 | <1% |
| Parent-child | Coyoacán | 11183 | 35% |
| | Iztapalapa | 19472 | 62% |
| | Mixed | 942 | 3% |
| Full siblings | Coyoacán | 10088 | 34% |
| | Iztapalapa | 17460 | 59% |
| | Mixed | 1934 | 7% |
| 2nd degree | Coyoacán | 11860 | 25% |
| | Iztapalapa | 28940 | 61% |
| | Mixed | 6280 | 13% |
| 3rd degree | Coyoacán | 22718 | 19% |
| | Iztapalapa | 76805 | 64% |
| | Mixed | 20657 | 17% |

**Supplementary Table 28** : Estimates of male and female ancestry contributions stratified by three ancestry components in the MCPS cohort, African, European and Native Mexican. The estimates are based on a simplified population mixture event model that best fits the observed X ancestry proportions (Bryc et al., 2015). The estimates of parameters (male and females ancestry contributions denoted as Pm and Pf, respectively) are obtained via a grid search with step of 0.005%. The proportion of female contribution per ancestry is calculated as Pf / (Pm + Pf) and reported in the last column.

| Ancestry | Autosomes | Chr. X | Pm | Pf | Pf / (Pm + Pf) |
|---|---|---|---|---|---|
| African | 3.50% | 3.20% | 2.10% | 1.50% | 41.40% |
| European | 31.80% | 22.70% | 29.40% | 2.40% | 7.50% |
| Native Mexican | 64.70% | 73.80% | 18.60% | 46.20% | 71.30% |

| **Supplementary Table 29** : Summary of runs of homozygosity | | | | | | |
|---|---|---|---|---|---|---|
| All 138,200 Individuals | | | | | | |
| | **Minimum** | **1st Quantile** | **Median** | **Mean** | **3rd Quantile** | **Maximum** |
| **% genome** | 0 | 0 | 0 | 0.343327 | 0.264189 | 34.05982 |
| **# segments** | 0 | 0 | 0 | 1.153466 | 1 | 51 |
| **Total length segments (Mb)** | 0 | 0 | 0 | 9.863901 | 7.590254 | 978.5518 |
| **Average length segment (Mb)** | 0 | 0 | 0 | 3.302302 | 4.681624 | 248.1359 |
| All 60,722 individuals with at least one 4cM ROH segment | | | | | | |
| | **Minimum** | **1st Quantile** | **Median** | **Mean** | **3rd Quantile** | **Maximum** |
| **% genome** | 0.010875 | 0.121064 | 0.356337 | 0.781393 | 0.958705 | 34.05982 |
| **# segments** | 1 | 1 | 2 | 2.625226 | 3 | 51 |
| **Total length segments (Mb)** | 0.312442 | 3.47822 | 10.23771 | 22.44971 | 27.54397 | 978.5518 |
| **Average length segment (Mb)** | 0.312442 | 2.889543 | 5.501573 | 7.515861 | 9.725092 | 248.1359 |

**Supplementary Table 30** : Number of rare homozygote pLoF variants within ROH segments and genome-wide, assigned by ancestry. Variants were assigned ancestries when haplotypes were concordant for ancestry and has posterior probability >0.8. AAF = Alternate Allele Frequency.

| | Within ROH | | | | Genome-wide | | | |
|---|---|---|---|---|---|---|---|---|
| Alternate allele frequency range | African | Native Mexican | European | Unassigned | African | Native Mexican | European | Unassigned |
| AAF<0.001 | 166 | 1186 | 543 | 71 | 219 | 1762 | 818 | 964 |
| 0.001<AAF<0.01 | 313 | 819 | 591 | 77 | 993 | 2318 | 1947 | 1715 |
| 0.01<AAF<0.05 | 189 | 1120 | 1100 | 91 | 1364 | 11339 | 13980 | 9550 |
| AAF>0.05 | 1264 | 48586 | 12953 | 3344 | 16101 | 4870834 | 1059360 | 4957683 |
| Total | 1932 | 51711 | 15187 | 3583 | 18677 | 4886253 | 1076105 | 4969912 |

**Supplementary Table 31** : Exome variant survey stratified by ancestry. Alternate allele counts on ancestry-specific haplotypes give three counts per-ancestry, multiplied by a scalar to interpolate to the whole-genome scale. The overall counts are calculated on all haplotypes and it is nearly equivalent to mean count numbers reported in the main Table 1. The number of samples is 138,092, for which phased WES data on autosomes and Chr. X is available.

| Variant category | Interpolated ancestry-specific mean number of alternate alleles per participant | | | |
|---|---|---|---|---|
| | Native Mexican | African | European | Overall |
| Coding regions | 29140 | 34411 | 28590 | 29140 |
| **Predicted function** | | | | |
| In-frame indels | 284 | 330 | 274 | 283 |
| Synonymous | 14922 | 17643 | 14587 | 14907 |
| Missense | 13587 | 16030 | 13375 | 13596 |
| Likely benign | 9917 | 11510 | 9768 | 9914 |
| Possibly deleterious | 3565 | 4283 | 3507 | 3567 |
| Likely deleterious | 107 | 215 | 123 | 115 |
| pLOF (any transcript) | 347 | 427 | 361 | 354 |
| Start lost | 27 | 31 | 27 | 27 |
| Stop gain | 86 | 98 | 81 | 85 |
| Stop lost | 13 | 18 | 13 | 13 |
| Splice donor | 38 | 46 | 37 | 38 |
| Frameshift | 138 | 190 | 159 | 146 |
| Splice acceptor | 44 | 45 | 44 | 44 |
